# A novel lineage of the Capra genus discovered in the Taurus Mountains of Turkey using ancient genomics

**DOI:** 10.1101/2022.04.08.487619

**Authors:** Kevin G. Daly, Benjamin S. Arbuckle, Conor Rossi, Valeria Mattiangeli, Phoebe A. Lawlor, Marjan Mashkour, Eberhard Sauer, Joséphine Lesur, Levent Atici, Cevdet Merih Erek, Daniel G. Bradley

## Abstract

Direkli Cave, located in the Taurus Mountains of southern Turkey, was occupied by Late Epipaleolithic hunters-gatherers for the seasonal hunting and processing of game including large numbers of wild goats. We report genomic data from new and published *Capra* specimens from Direkli Cave and, supplemented with historic genomes from multiple *Capra* species, find a novel lineage best represented by a ∼14,000 year old 2.59X genome sequenced from specimen Direkli4. This newly discovered Capra lineage is a sister clade to the Caucasian tur species (*Capra cylindricornis* and *Capra caucasica*), both now limited to the Caucasus region. We identify genomic regions introgressed in domestic goats with high affinity to Direkli4, and find that West Eurasian domestic goats in the past, but not those today, appear enriched for Direkli4-specific alleles at a genome-wide level. This forgotten “Taurasian tur” likely survived Late Pleistocene climatic change in a Taurus Mountain refugia and its genomic fate is unknown.

## Introduction

The genus *Capra* includes the domestic goat (*Capra hircus*) as well as a variety of wild mountain-dwelling goat/ibex species distributed across Eurasia and North Africa including several listed as endangered or vulnerable (Shackleton 1997; Pidancier et al. 2006). Nine species are currently recognized by the IUCN; however, taxonomic relationships are still under revision (Pidancier et al. 2006; Zheng et al. 2020). Among these, the status of the two species endemic to the Caucasus Mountains has been debated (Parrini, Cain, and Krausman 2009; Groves and Grubb 2011). The East Caucasian tur (*Capra cylindricornis*) has been considered either a species distinct from the West Caucasian tur (*Capra caucasica*) or they comprise a single species of two potentially-hybridizing populations (Heptner, Nasimovich, and Bannikov 1961). Moreover the bezoar (*Capra aegagrus*), progenitor of domestic goat, has also been reported to hybridize with both tur varieties with which it shares seasonal grazing territories in the Caucasus region (Pfitzenmayer 1915; Sarkisov 1953; Weinberg 2002). Interspecies *Capra* gene flow is well known (Manceau et al. 1999; Pidancier et al. 2006; Kazanskaia, Kuznetsova, and Danilkin 2007), and may explain discordant phylogenies across loci (Pidancier et al. 2006; Ropiquet and Hassanin 2006). Such admixture may have shaped the evolution of domestic goat; for example, the tur has been identified as a putative source of a *MUC6* allele driven to fixation in domestic populations and likely selected for gastrointestinal parasite resistance (Zheng et al. 2020; Grossen et al. 2020). Tur additionally shows differing affinity to domestic and wild goat genomes indicating a complex evolutionary history of the genus.

Although tur are currently restricted to the Caucasus region, ancient wild goat specimens recovered from Direkli Cave, a camp site used by Late Pleistocene hunters in the Central Taurus Mountains of southern Turkey (Figures 1A, S1 and S2) (Arbuckle and Erek 2012), were found to carry a tur-like mitochondrial lineage, designated T (Daly et al. 2018). Three of these four reported ancient *Capra* specimen fall within the bezoar autosomal diversity, but a fourth - Direkli4, dated to 12,164-11,864 cal BCE (Ramsey 2009; Reimer et al. 2020) and sequenced here to 2.59X mean genome coverage (Table 1) - shows an excess of ancestral alleles in *D* statistic tests (Green et al. 2010) when paired with domestic/bezoar goat (Figure S3, Table S2) implying Direkli4 carries ancestry basal to that clade. To explore this signal further we generated low coverage genomes from historic and rare *Capra* samples, including a 20th century CE zoo-born East Caucasian tur (Tur2), tur specimens from the Dariali-Tamara Fort archaeological site near Kazbegi, Georgia (Mashkour et al. 2020), a zoo-born Walia ibex, and supplemented with published modern and ancient *Capra* genomes (Table S1, S3, Figure S4) (Zheng et al. 2020; Grossen et al. 2020). Surprisingly, a neighbour joining tree from nuclear genome identity-by-state (IBS) information places Direkli4 as sister to a clade of both Caucasian tur taxa, a signal obtained using either goat- or sheep-aligned data (Figures 1B and S5). The Direkli4 genome thus suggests a previously-unrecognized *Capra* lineage sister to both Caucasian tur inhabited the Taurus Mountains ∼14,000 years ago.

**Figure 1.**
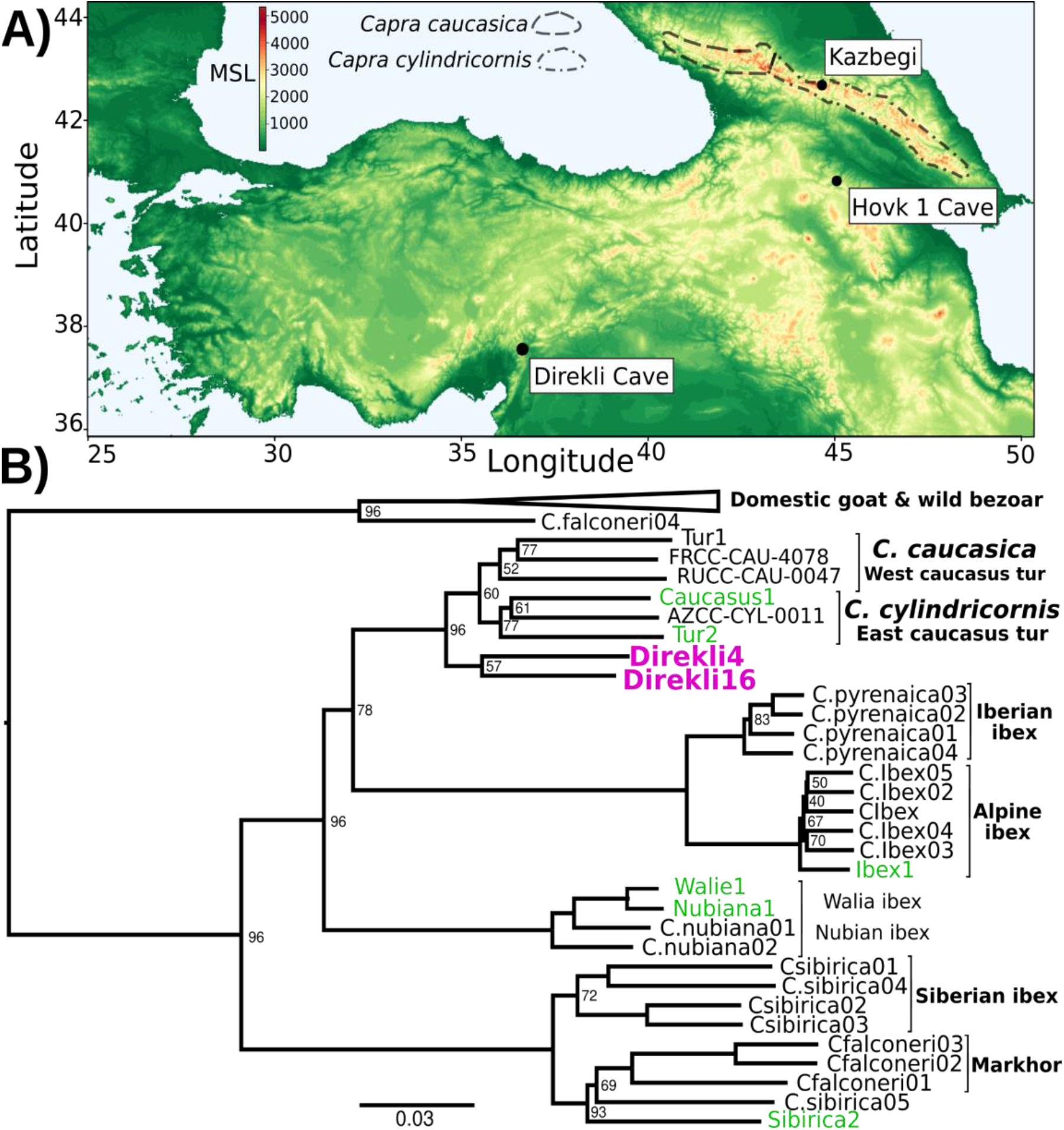
A) Elevation map of southwest Asia. Key sites are indicated, with *C. caucasica* and *C. cylindricornis* distributions from (Gavashelishvili et al. 2018) displayed. MSL=metres above mean sea level. B) Neighbour joining phylogeny of genomes >0.5X and the lower coverage Tur2 (0.02X) and Direkli16 (0.01X) genomes using 625,495 transversion sites and pairwise IBS, rooted on Sheep (not shown, as well as a likely Barbary Sheep sample Falconeri1, see Supplementary Methods). Node support from 100 replicates using 50 5Mb regions sampled without replacement shown when <100. Pink=Direkli4, green=other genomes first reported here.

**Table 1.** Sample provenance and sequencing summary.

| Sample | Morphological Species | Origin | Age | Sex<sup>b</sup> | Nuclear Cov.<sup>c</sup> | mtDNA Cov. |
| --- | --- | --- | --- | --- | --- | --- |
| Direkli4 | *Capra spc.* | E4/8A, Direkli Cave, Turkey | 12,164-11,864 cal BCE (2σ) | M | 2.59 | 642.23 |
| Direkli9 | *Capra spc.* | B6/5, Direkli Cave, Turkey | Est. 12,100-8,900 BCE<sup>a</sup> | M | 0.0003 | 9.5 |
| Direkli12 | *Capra spc.* | B13/4A, Direkli Cave, Turkey | Est. 12,100-8,900 BCE | M | 0.07 | 44.06 |
| Direkli13 | *Capra spc.* | B13/7B, Direkli Cave, Turkey | Est. 12,100-8,900 BCE | M | 0.0055 | 76.99 |
| Direkli14 | *Capra spc.* | D3/7, Direkli Cave, Turkey | Est. 12,100-8,900 BCE | M | 0.0005 | 13.24 |
| Direkli15 | *Capra spc.* | B6/5, Direkli Cave, Turkey | Est. 12,100-8,900 BCE | F | 0.0002 | 24.21 |
| Direkli16 | *Capra spc.* | B8/7, Direkli Cave, Turkey | Est. 12,100-8,900 BCE | F | 0.01 | 15.11 |
| Direkli17 | *Capra spc.* | E5/5, Direkli Cave, Turkey | Est. 12,100-8,900 BCE | F | 0.0001 | 0.0845 |
| Caucasus1 | *Capra caucasica* | Tamara Fort, Kazbegi, Georgia | 4th-21st c. CE, probably 5th-15th c. CE | M | 0.55 | 64.63 |
| Caucasus2 | *Capra caucasica* | Tamara Fort, Kazbegi, Georgia | 4th-21st c. CE, probably 5th-15th c. CE | M | 0.0021 | 379.71 |
| Caucasus3 | *Capra caucasica* | Tamara Fort, Kazbegi, Georgia | 4th-21st c. CE, probably 5th-15th c. CE | M | 0.004 | 464.51 |
| Falconeri1 | *Capra falconeri hepteneri*<sup>d</sup> | Unknown, via Parc de la Haute-Touche | 20th Century CE | M | 0.58 | 72.11 |
| Falconeri2 | *Capra falconeri* | Born at MNHN Zoo, Paris | 20th Century CE | M | 0.06 | 45.78 |
| Ibex1 | *Capra ibex* | Unknown | 20th Century CE | F | 3.93 | 179.03 |
| Ibex2 | *Capra ibex* | Pointe de Calabre, Savoie | 20th Century CE | M | 0.05 | 21.4 |
| Sibirica1 | *Capra sibirica* | Born at MNHN Zoo, Paris | 20th Century CE | M | 0.04 | 163.16 |
| Sibirica2 | *Capra sibirica* | Born at MNHN Zoo, Paris | 20th Century CE | F | 1.48 | 205.78 |
| Tur2 | *Capra cylindricornis* | Unknown, via Vincennes Zoo | 20th Century CE | F | 0.02 | 4.96 |
| Walie1 | *Capra walie* | Born at MNHN Zoo, Paris | 20th Century CE | M | 0.75 | 103.12 |
| Pyrenaica2 | *Capra pyrenaica* | Unknown | 20th Century CE | M | 0.16 | 51.2 |
| Nubiana1 | *Capra nubiana* | Unknown | 20th Century CE | M | 1.25 | 211.32 |

<sup>a</sup>Estimated ages for Direkli material is based on calibrated ages from the cave stratigraphy. <sup>b</sup>M=Male, F=Female
<sup>c</sup>Cov. = coverage.
<sup>d</sup>Falconeri1, a likely Barbary sheep (see Supplementary Methods).

## Materials and methods

DNA from 7 postcranial bone elements from Direkli Cave and 13 historic *Capra* specimen was extracted via standard aDNA protocols (Yang et al. 1998), with a 0.5% sodium hypochlorite pre-wash (Korlević et al. 2015) performed for the Direkli material. Following uracil excision (Rohland et al. 2015) and dsDNA library construction (Meyer and Kircher 2010), libraries were subject to shotgun sequencing (Illumina HiSeq 2000 and NovaSeq 6000) or RNA-bait enrichment of mtDNA reads prior to shotgun sequencing. Additional sequencing data was also generated for specimen Direkli4.

Using bwa aln (Li, Ruan, and Durbin 2008) a relaxed alignment (-n 0.01 -o 2, Meyer et al. 2012) was performed against the goat reference ARS1 (Bickhart et al. 2017) or an outgroup genome (Oar_rambouillet_v1.0). Subsequent analyses were primarily performed in the ANGSD environment (Korneliussen, Albrechtsen, and Nielsen 2014) using single read sampling. A more detailed methodology is provided in the Supplementary Material, available at OSF (https://doi.org/10.17605/OSF.IO/3ECQD).

## Results

Our additional screening of Direkli Cave *Capra* remains identified seven with surviving DNA (Table S4); two genomes show greater affinity to Direkli4 than bezoar from the same site (Figure 2A, Table S5). An MDS plot of IBS distances (Figure S6) places two Direkli samples with sufficient coverage (Direkli4 and Direkli16) close to East and West Caucasian tur genome clusters, with a slight bias to the former. This tur affinity is unlikely to be driven by error as Direkli specimens have low error rates (0.026-0.195%, Table S1 and S3) and do not show inflated distance-to-the-outgroup relative to modern genomes (Figure S7). A total of three out of the eleven Direkli Cave *Capra* specimens therefore are assigned to the tur-related clade, implying that while less numerous than bezoar, members of this clade were not rare in the region in the Late Pleistocene. Nuclear genome types (tur-like or bezoar) do not necessarily co-associate with mitochondrial lineages (tur T and bezoar F), with all combinations except “tur-like genome, tur-like mitochondria” observed (Figure 2B, Figures S8 and S9, Table S5), establishing that there was gene flow between these lineages. Additionally, there is little variation among Direkli T mtDNA (average 4.07 pairwise differences, compared to 67 among Direkli F mtDNA), suggesting a limited population size for this Direkli tur-like matrilineage.

**Figure 2.**
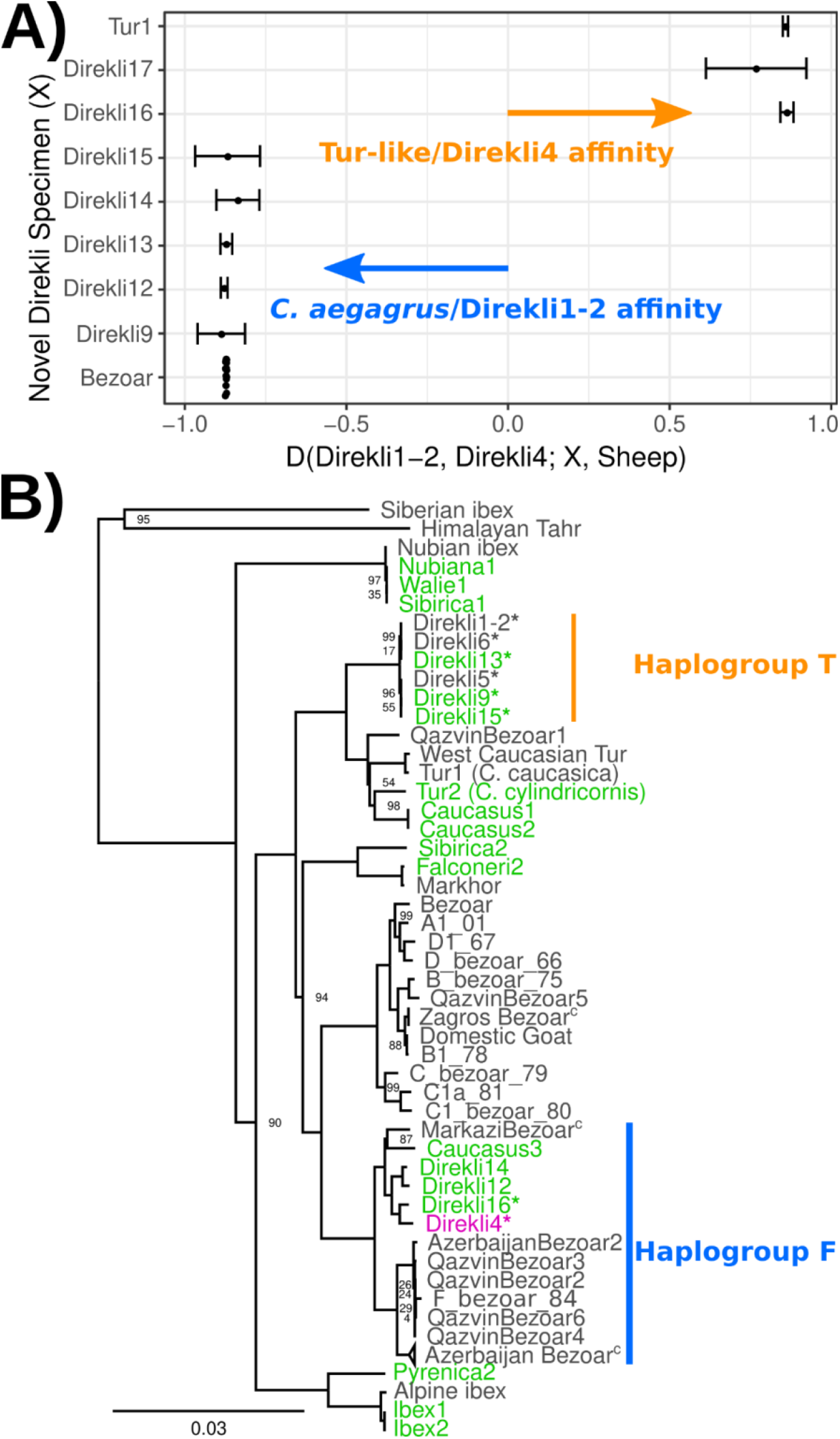
A) *D* statistic test of affinity using specimens from Direkli Cave and a historic *C. caucasica* individual for reference. Positive values indicate sample X has greater affinity with the Tur-like Direkli4 genome; negative values indicate greater affinity with the *C. aegagrus* Direkli1-2. Error bars represent 3 standard errors, underlying site counts are presented in Table S5. B) ML phylogeny of mtDNA, abbreviated. Bootstrap node support values (100 replicates) are displayed when <100. The complete phylogeny including likely Barbary sheep Falconeri1 is displayed in Figure S8. T haplogroup is as defined by (Daly et al. 2018) Low coverage sample Direkli17 is displayed in a highly reduced phylogeny Figure S9, C=collapsed. *=Direkli sample with discordant mtDNA and nuclear genome affinity.

A tur-like population in the Taurus Mountains is consistent with the high variability in body size of the *Capra* material at Direkli Cave where extremely large *Capra* remains have been reported alongside smaller bezoar-size individuals (Figure S10). Extant tur exhibit body weights 20-50% larger than bezoar (Masseti 2009; Castelló, Huffman, and Groves 2016) and it is plausible that the ‘large’ *Capra* from Direkli represent tur-lineage animals. Although there are clear differences between bezoar and tur horn morphologies (Pidancier et al. 2006), unfortunately, diagnostic horncore remains have not been recovered from Direkli. The cave was initially inferred to be occupied primarily during summer months (Arbuckle and Erek 2012), with subsequent discoveries of architectural remains and zooarchaeological analyses indicating more intensive use (Arbuckle 2019). The presence of tur-related goats may reflect use of the cave in the winter months when, based on Caucasian analogs (Gavashelishvili 2009), tur would be expected to descend from the higher elevations surrounding the cave (>2000m above sea level).

Given the Direkli4 genome was recovered together with bezoar specimens, the two lineages of *Capra* likely had proximate ranges and hybridised. We use *D* statistics (Green et al. 2010) to measure Direkli4 derived allele sharing relative to either a likely-hunted (Table S7) or likely-herded (Table S8, Pearson’s r >0.99) ∼10,000 year old goat from the Zagros Mountains. A Late Pleistocene wild goat from the Armenian Lesser Caucasus, Hovk1, shows highest affinity with Direkli4 (Figure S11). Bezoar goats from Direkli Cave also show high Direkli4 allele sharing, mirroring affinity measures with west Caucasian tur (Zheng et al. 2020). While directionality is uncertain, these statistics imply gene flow between the tur-like lineage and wild bezoar.

Examining domestic goats we find that Neolithic genomes from Europe show greater affinity to Direkli4 (Figure S11), but Neolithic Iranian goats do not, echoing the distribution of Direkli bezoar-related ancestry in West Eurasian populations (Daly et al. 2018). We account for possible gene flow from Caucasian tur into modern European goat using the statistic *D*(Tur1, Direkli4; X, Sheep) to compare relative affinity with Tur1 and Direkli4 (Table S9). With the exception of two other tur samples, all examined domestic/bezoar goats show either a bias towards Direkli4 or gave a non-significant result, consistent with Direkli4-related admixture or a more complicated genetic history.

Genetic exchange between bezoar and the ancestors of Direkli4 could confound these measures of shared variation among domestic populations. We identified variants specific to Direkli4, conditioned on ancestral allele fixation in a range of defined groups (Figure 3A, Supplementary). Using this we calculate a statistic analogous to the *D* statistic, here termed the extended *D* or *D*_*ex*_. *D*_*ex*_ measures the relative degree of allele sharing, derived specifically in a selected genome or group of genomes, and may have some utility in genera with complex, admixture histories or admixture from ghost lineages. Relative to Neolithic Zagros goats, ancient domestic genomes from western Eurasia have an excess of Direkli4-specific variants (Figure 3B, Figure S12, Table S10). This “Direkli4-specific” allele sharing signal is absent in ancient goats from Iran-eastwards, and in all tested modern goats (Figure 3C). To control for possible reference biases, we calculated *D*_*ex*_ ascertaining on variants segregating in sheep (Table S11) and recovered similar results (Pearson’s *r* = 0.9935). Repeating the analysis using other ancient/historic *Capra* “specific” alleles shows somewhat correlated results (Table S12), but the distinct patterns of allele sharing (Figure S13, Table S13) imply that Direkli4 ancestry in domestic goat varies temporally and geographically.

**Figure 3.**
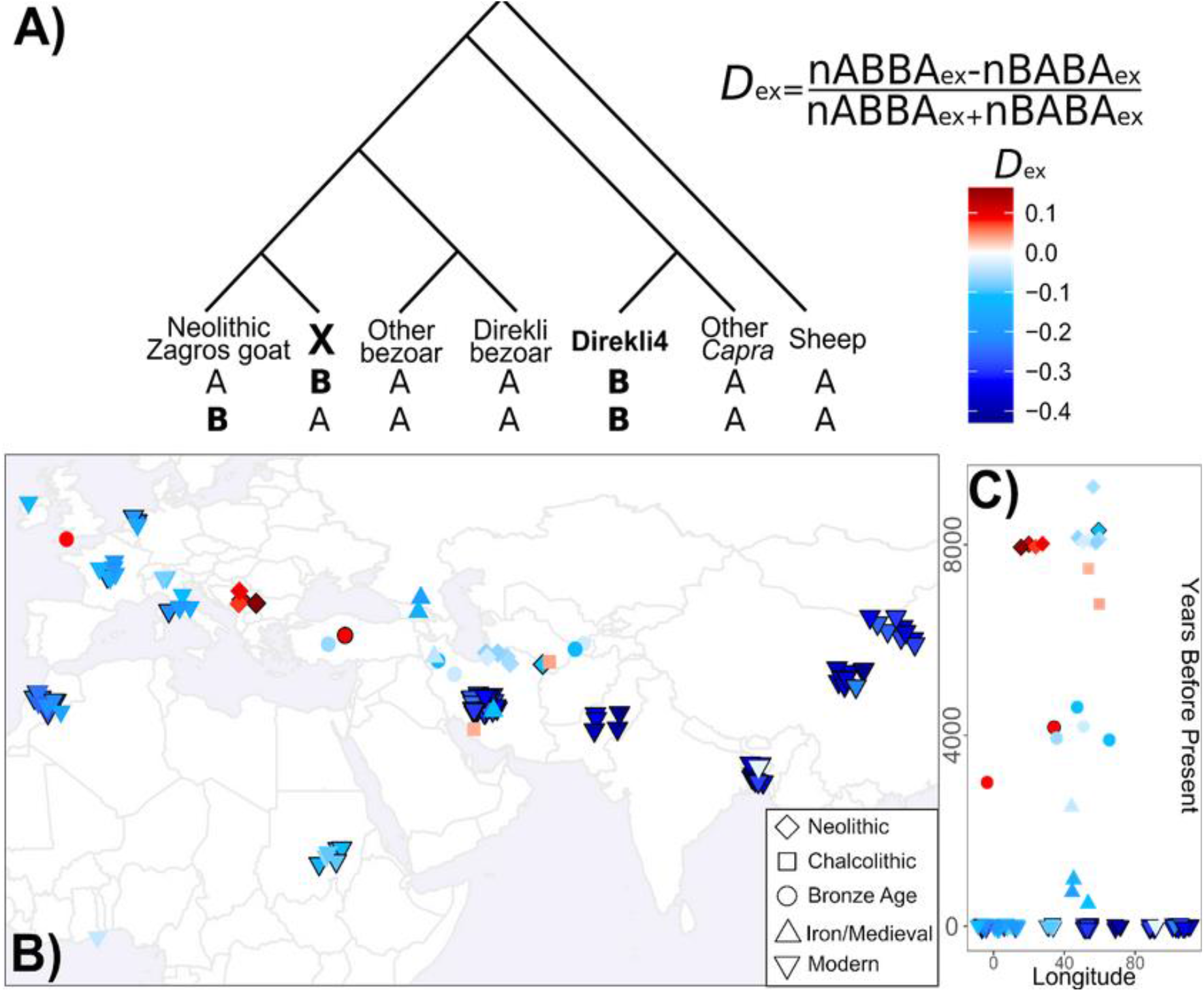
A) Extended D statistic. To control for gene flow with the *Capra* genus, we condition on variants derived in Direkli4 (H3) and a genome X (H2), but ancestral in other populations (here: Sheep, non-bezoar *Capra* genomes, bezoar from Direkli Cave, and other bezoar). Values are calculated relative to a set of Neolithic goat from the Zagros Mountains, and normalized similarly to the D statistic. B) Extended D statistic values for Direkli4 using transversions, C) plotted through time. Each symbol is a test genome, with shape denoting time period. Black borders indicates a |Z| score ≥ 3, using 1000 bootstrap replicates and 5Mb blocks.

We next identify alleles derived in Direkli4 also at a low frequency (>0%, ≤10%) in other *Capra* and bezoar, and then measure their abundance in domestic goats. The west Caucasian tur (Tur1) most frequently shares derived alleles with Direkli4 and domestic goat (Figure S14, consistent with their cladal relationship (although this measure is sensitive to genome depth, Supplementary Methods). Ancient European domestic goats share a higher proportion of alleles with both Direkli4 and the high-coverage bezoar from Direkli Cave, Direkli1-2. In comparison, Modern European and African goats carry variation present in Direkli4 plus one of the two Caucasian tur (Tur1 and Caucasus1). This discrepancy could be explained by either gene flow from domestic goats into tur during the last 8,000 years, or alternatively an increase in tur-Direkli4 related ancestry in European populations over time.

Investigating gene flow events within *Capra*, automated tree-based model exploration (Pickrell and Pritchard 2012) detects admixture between the Direkli4/Tur lineage and the ancestors of the Late Pleistocene bezoar Hovk-1 (Figure S15). Residuals of this graph point to unmodelled affinity between Direkli4 and both Direkli Cave bezoar and with Neolithic Serbian domestic goat (Figure S16). Modern European goats do not show unmodeled Direkli4 affinity, supporting the interpretation that Direkli4-related ancestry has declined with time in west Eurasian goats. A reduced set of populations explored using ML network orientation (Molloy, Durvasula, and Sankararaman 2021) reiterates the Tur1/Direkli4 and Direkli bezoar lineages admixture, and also between Direkli bezoar and domestic goat (Figure S17). Investigating admixture graph space (Supplementary Methods, Maier et al. 2022) we find 2 admixture events best explain how a subset of populations (Sheep, Tur1, Direkli4, Direkli bezoar, Neolithic East Iran, and Neolithic Serbia) can be modelled. A majority (6/11) of graphs model Direkli bezoar as containing ancestry related to Direkli4 (median 1.5%, mean 5.2%; best fitting graph is shown in Figure S18), with a single graph modelling the opposite (2% Direkli bezoar ancestry). While the graph space explored is limited, these results suggest a greater degree of “Direkli4 to Direkli bezoar” gene flow than “Direkli bezoar to Direkli4”.

We finally identify 3 out of 112 regions introgressed from other *Capra* species to domestic goats (Zheng et al. 2020) which show high affinity with Direkli4 (Figures S18-20, S31-32, Supplementary Data Files 1 and 2). A further 7 regions appear to have most affinity with the Direkli4-tur clade (Figures S22-30), including a locus encompassing *MUC6*, a target of selection in domestic goats during the last 10,000 years (Zheng et al. 2020), implicating the Direkli4 lineage in the makeup of domestic goat gene pool.

## Discussion

Our results indicate that a lineage related to the Caucasian tur existed in the Taurus Mountains during the Late Pleistocene, as late as the 12th millennium cal BCE. Based on the current, limited genomic data from the *Capra* genus, which we improve on here, this lineage appears to be a sister group to the tur *C. caucasica* and *C. cylindricornis*. Similar to other mammalian groups (Gopalakrishnan et al. 2018; Palkopoulou et al. 2018; Zheng et al. 2020), admixture likely occurred among *Capra* lineages; the population reported here carries bezoar-associated mtDNA and a possible small amount of bezoar nuclear genome ancestry (2% from 1/12 graphs). The Taurasian tur population is itself a possible candidate for the source of Tur-like ancestry present in domestic goats, including an introgressed *MUC6* allele fixed in modern populations which increases gastrointestinal parasite resistance (Zheng et al. 2020). Given the relative paucity of *Capra* genomic data available compared to other mammalian groups, additional genomes from the genus will help refine the history of divergences and gene flow events which shaped the group’s evolution.

We suggest this novel “Taurasian tur” lineage be designated *Capra taurensis* following IUCN convention (Weinberg and Lortkipanidze 2020) or *Capra caucasica taurensis* under a subspecies classification (Wilson and Reeder 2005) if and when sufficient multi-disciplinary data allows for taxonomic delimitation (De Queiroz 2007). The Taurasian tur may have diverged from the Caucasian lineages 130-200kya based on mtDNA coalescent estimates (Bouckaert et al. 2014; Daly et al. 2018). The current distribution of *Capra* species is mostly discontinuous and is suggestive of climate-induced fragmentation (Shackleton 1997). The ancestors of Caucasian tur likely extended over a broader range in Eurasia during the Late Pleistocene but may have been poorly captured by the fossil record (Uerpmann 1987; Crégut-Bonnoure 1991; Weinberg 2002). The large variability and high upper size range of *Capra* remains are consistent with both smaller-bodied bezoar and larger-bodied tur-relatives being present within the faunal assemblage at Direkli as well as other sites in the central Taurus and Lebanese mountains (Üçagızlı cave, Ksar Akil, Saaide II), but not in the western Taurus where only bezoar are evident (Figure S10) (Arbuckle and Erek 2012). *C. taurensis* could have survived the Last Glacial Maximum within the central Taurus Mountains, a plausible refugia for *Capra* species in addition to the Pontic and Anti-Taurus ranges (Gavashelishvili et al. 2018) while experiencing a severe matrilineal bottleneck (Figure 2B). *C. taurensis* appears to have produced fertile offspring with other members of the *Capra* genus; the traces of shared ancestry in ancient bezoar and likely managed goat (Figure 3B) may be the consequence of direct gene flow or secondarily via admixed bezoar. Gene flow between early managed goats and Anatolian bezoar carrying *C. taurensis* ancestry could partially explain the divergence between Zagros and more westerly herds.

Given the tremendous pressure on *Capra* species via Anthropocene over-hunting and habitat disruption (Shackleton 1997), it is assumed that *C. taurensis* is extinct, with its existence only now revealed via palaeogenomics. The Caucasian tur’s preference for snowier habitats (Gavashelishvili et al. 2018) combined with the lower altitude of the Taurus Mountains relative to the Caucasus (Figure 1A) may have rendered the lineage vulnerable to climatic change via Holocene warming and interspecific competition with bezoar, which are still found in the Taurus mountains (Naderi et al. 2008; Gavashelishvili 2009), leading to its hypothesised extinction. As the history of *C. taurensis* following the Late Pleistocene is still unknown, further genomic surveys of Holocene *Capra* remains and present-day populations, such as the VarGoats project (Denoyelle et al. 2021), from this and adjacent regions may illuminate its genetic legacy.

## Acknowledgements

We thank Matthew Teasdale and Amelie Scheu for their advice on interpretation of results and helpful discussions. Excavations at Direkli Cave are sponsored by T.C. Kültür ve Turizm Bakanlığı. Permission to export samples from Direkli Cave provided by T.C. Kültür Varlıkları ve Müzeler Genel Müdürlüğü and T.C. Ankara Valılığı, Il Kültür ve Turizm Müdürlüğü, Anadolu Medeniyetleri Müzesi Müdürlüğü (#70583208-160.99(06)-899). We thank the VarGoats consortia for use of the modern tur sequencing data in IBS analyses. Version 5 of this preprint has been peer-reviewed and recommended by Peer Community In Genomics (https://doi.org/10.24072/pci.genomics.100020).

## Data, scripts and codes availability

Raw sequencing reads, aligned QCed final bam files, and mitochondrial fasta files have been deposited in ENA under the project accession PRJEB51668. Admixture graphs “as good as” the best fitting graph are available at https://osf.io/3ecqd/. *Capra taurensis* has been registered under the Zoobank LSID urn:lsid:zoobank.org:act:1261A42B-B0C0-4571-87F4-8EC3B5381A88. Scripts for extended D calculation/disentangling derived allele sharing are available at https://osf.io/3ecqd/.

## Supplementary material

Supplementary material, including supplementary files 1 and 2 (Figure S31 and S32) are available online: https://doi.org/10.17605/OSF.IO/3ECQD.

## Conflict of interest disclosure

The authors of this preprint declare that they have no financial conflict of interest with the content of this article.

## Funding

Zooarchaeological work at Direkli has been supported by grants from the Office of Vice Provost for Research, Baylor University and a URC grant from the Office of the Vice Chancellor for Research at the University of North Carolina at Chapel Hill. This work was supported by the European Research Council under the European Union’s Horizon 2020 research and innovation programme (grant numbers 885729-AncestralWeave, 295729-CodeX, 295375-Persia and its Neighbours); and supported in part by a Grant from Science Foundation Ireland under grant number 21/PATH-S/9515.

## Notes

https://osf.io/3ecqd/files/

https://github.com/Xevkin/direkli_caprid_extended_ds

### Competing Interest Statement

The authors have declared no competing interest.

### Summary of Updates

Updated to fit with PCI Genomics recommended template

## References

Arbuckle, B. S. (2019). Zooarchaeology at Epipaleolithic Direkli Cave, Kahramanmaras, Turkey. Journal of Anatolian Prehistoric Research/Anadolu Prehistorya Araştırmaları Dergisi, 5, 1–14.

Arbuckle, B. S., & Erek, C. M. (2012). Late Epipaleolithic hunters of the central Taurus: Faunal remains from Direkli Cave, Kahramanmaraş, Turkey. International Journal of Osteoarchaeology, 22(6), 694–707. https://doi.org/10.1002/oa.1230

Bickhart, D. M., Rosen, B. D., Koren, S., Sayre, B. L., Hastie, A. R., Chan, S., Lee, J., Lam, E. T., Liachko, I., Sullivan, S. T., Burton, J. N., Huson, H. J., Nystrom, J. C., Kelley, C. M., Hutchison, J. L., Zhou, Y., Sun, J., Crisà, A., Ponce de León, F. A., … Smith, T. P. L. (2017). Single-molecule sequencing and chromatin conformation capture enable de novo reference assembly of the domestic goat genome. Nature Genetics, 49(4), 643–650. https://doi.org/10.1038/ng.3802

Bouckaert, R., Heled, J., Kühnert, D., Vaughan, T., Wu, C.-H., Xie, D., Suchard, M. A., Rambaut, A., & Drummond, A. J. (2014). BEAST 2: a software platform for Bayesian evolutionary analysis. PLoS Computational Biology, 10(4), e1003537. https://doi.org/10.1371/journal.pcbi.1003537

Castelló, J. R., Huffman, B., & Groves, C. (2016). Bovids of the World: Antelopes, Gazelles, Cattle, Goats, Sheep, and Relatives (Vol. 104). Princeton University Press.

Crégut-Bonnoure, E. (1991). Intérêt biostratigraphique de la morphologie dentaire de Capra (Mammalia, Bovidae). Annales Zoologici Fennici, 28(3/4), 273–290.

Daly, K. G., Maisano Delser, P., Mullin, V. E., Scheu, A., Mattiangeli, V., Teasdale, M. D., Hare, A. J., Burger, J., Verdugo, M. P., Collins, M. J., Kehati, R., Erek, C. M., Bar-Oz, G., Pompanon, F., Cumer, T., Çakırlar, C., Mohaseb, A. F., Decruyenaere, D., Davoudi, H., … Bradley, D. G. (2018). Ancient goat genomes reveal mosaic domestication in the Fertile Crescent. Science, 361(6397), 85–88. https://doi.org/10.1126/science.aas9411

Denoyelle, L., Talouarn, E., Bardou, P., Colli, L., Alberti, A., Danchin, C., Del Corvo, M., Engelen, S., Orvain, C., Palhière, I., Rupp, R., Sarry, J., Salavati, M., Amills, M., Clark, E., Crepaldi, P., Faraut, T., Masiga, C. W., Pompanon, F., … VarGoats Consortium. (2021). VarGoats project: a dataset of 1159 whole-genome sequences to dissect Capra hircus global diversity. Genetics, Selection, Evolution: GSE, 53(1), 86. https://doi.org/10.1186/s12711-021-00659-6

De Queiroz, K. (2007). Species concepts and species delimitation. Systematic Biology, 56(6), 879–886. https://doi.org/10.1080/10635150701701083

Gavashelishvili, A. (2009). GIS-based Habitat Modeling of Mountain Ungulate species in the Caucasus Hotspot. In N. Zazanashvili & D. Mallon (Eds.), Status and Protection of Globally Threatened Species in the Caucasus (pp. 74–82).

Gavashelishvili, A., Yarovenko, Y. A., Babayev, E. A., Mikeladze, G., Gurielidze, Z., Dekanoidze, D., Kerdikoshvili, N., Ninua, L., & Paposhvili, N. (2018). Modeling the distribution and abundance of eastern tur (Capra cylindricornis) in the Caucasus. Journal of Mammalogy. https://doi.org/10.1093/jmammal/gyy056

Gopalakrishnan, S., Sinding, M.-H. S., Ramos-Madrigal, J., Niemann, J., Samaniego Castruita, J. A., Vieira, F. G., Carøe, C., Montero, M. de M., Kuderna, L., Serres, A., González-Basallote, V. M., Liu, Y.-H., Wang, G.-D., Marques-Bonet, T., Mirarab, S., Fernandes, C., Gaubert, P., Koepfli, K.-P., Budd, J., … Gilbert, M. T. P. (2018). Interspecific Gene Flow Shaped the Evolution of the Genus Canis. Current Biology: CB, 28(21), 3441–3449.e5. https://doi.org/10.1016/j.cub.2018.08.041

Green, R. E., Krause, J., Briggs, A. W., Maricic, T., Stenzel, U., Kircher, M., Patterson, N., Li, H., Zhai, W., Fritz, M. H.-Y., Hansen, N. F., Durand, E. Y., Malaspinas, A.-S., Jensen, J. D., Marques-Bonet, T., Alkan, C., Prüfer, K., Meyer, M., Burbano, H. A., … Pääbo, S. (2010). A draft sequence of the Neandertal genome. Science, 328(5979), 710–722. https://doi.org/10.1126/science.1188021

Grossen, C., Guillaume, F., Keller, L. F., & Croll, D. (2020). Purging of highly deleterious mutations through severe bottlenecks in Alpine ibex. Nature Communications, 11(1), 1001. https://doi.org/10.1038/s41467-020-14803-1

Groves, C., & Grubb, P. (2011). Ungulate Taxonomy. JHU Press.

Heptner, V. G., Nasimovich, A. A., & Bannikov, A. G. (1961). Mammals of the Soviet Union (V. G. Heptner & N. P. Naumov (eds.); pp. 679–679). Vysshaya Shkola. https://doi.org/10.2307/1381452

Kazanskaia, E. I., Kuznetsova, M. V., & Danilkin, A. A. (2007). [Phylogenetic reconstructions in the genus Capra (Bovidae, Artiodactyla) based on the mitochondrial DNA analysis]. Genetika, 43(2), 245–253.

Korneliussen, T. S., Albrechtsen, A., & Nielsen, R. (2014). ANGSD: Analysis of Next Generation Sequencing Data. BMC Bioinformatics, 15, 356. https://doi.org/10.1186/s12859-014-0356-4

Li, H., Ruan, J., & Durbin, R. (2008). Mapping short DNA sequencing reads and calling variants using mapping quality scores. Genome Research, 18(11), 1851–1858. https://doi.org/10.1101/gr.078212.108

Maier, R., Flegontov, P., Flegontova, O., Changmai, P., & Reich, D. (2022). On the limits of fitting complex models of population history to genetic data. bioRxiv 2022.05.08.491072. https://doi.org/10.1101/2022.05.08.491072

Manceau, V., Després, L., Bouvet, J., & Taberlet, P. (1999). Systematics of the genus Capra inferred from mitochondrial DNA sequence data. Molecular Phylogenetics and Evolution, 13, 504–510. https://doi.org/10.1006/mpev.1999.0688

Mashkour, M., Amiri, S., Fathi, H., Khazaeli, R., Debue, K., Decruyenaere, D., Doost, S. B., Kamjan, S., & Sauer, E. W. (2020). Herding and Hunting in the highlands from the Sasanian to Late Medieval periods: The archaeozoology of the Dariali Gorge. In E. W. Sauer (Ed.), Dariali: The “Caspian Gates” in the Caucasus from Antiquity to the Age of the Huns and the Middle Ages: The Joint Georgian-British Dariali Gorge Excavations and Surveys of 2013–2016 (Vol. 6, pp. 729–780). Oxbow.

Masseti, M. (2009). The wild goatsCapra aegagrus Erxleben, 1777 of the Mediterranean Sea and the Eastern Atlantic Ocean islands. Mammal Review, 39(2), 141–157. https://doi.org/10.1111/j.1365-2907.2009.00141.x

Meyer, M., & Kircher, M. (2010). Illumina sequencing library preparation for highly multiplexed target capture and sequencing. Cold Spring Harbor Protocols, 2010(6), db.prot5448. https://doi.org/10.1101/pdb.prot5448

Meyer, M., Kircher, M., Gansauge, M.-T., Li, H., Racimo, F., Mallick, S., Schraiber, J. G., Jay, F., Prüfer, K., de Filippo, C., Sudmant, P. H., Alkan, C., Fu, Q., Do, R., Rohland, N., Tandon, A., Siebauer, M., Green, R. E., Bryc, K., … Pääbo, S. (2012). A high-coverage genome sequence from an archaic Denisovan individual. Science, 338(6104), 222–226. https://doi.org/10.1126/science.1224344

Molloy, E. K., Durvasula, A., & Sankararaman, S. (2021). Advancing admixture graph estimation via maximum likelihood network orientation. Bioinformatics, 37(Suppl_1), i142–i150. https://doi.org/10.1093/bioinformatics/btab267

Naderi, S., Rezaei, H.-R., Pompanon, F., Blum, M. G. B., Negrini, R., Naghash, H.-R., Balkiz, O., Mashkour, M., Gaggiotti, O. E., Ajmone-Marsan, P., Kence, A., Vigne, J.-D., & Taberlet, P. (2008). The goat domestication process inferred from large-scale mitochondrial DNA analysis of wild and domestic individuals. Proceedings of the National Academy of Sciences of the United States of America, 105, 17659–17664. https://doi.org/10.1073/pnas.0804782105

Palkopoulou, E., Lipson, M., Mallick, S., Nielsen, S., Rohland, N., Baleka, S., Karpinski, E., Ivancevic, A. M., To, T.-H., Kortschak, R. D., Raison, J. M., Qu, Z., Chin, T.-J., Alt, K. W., Claesson, S., Dalén, L., MacPhee, R. D. E., Meller, H., Roca, A. L., … Reich, D. (2018). A comprehensive genomic history of extinct and living elephants. Proceedings of the National Academy of Sciences of the United States of America. https://doi.org/10.1073/pnas.1720554115

Parrini, F., Cain, J. W., & Krausman, P. R. (2009). Capra ibex (Artiodactyla: Bovidae). Mammalian Species, 830, 1–12. https://doi.org/10.1644/830.1

Pfitzenmayer, E. V. (1915). Nekotoryye interesnyye ublyudki semeistva polorogikh iz Zakavkaz’ya [Some interesting bovid mongrels from Transcaucasia]. Izvestiya Kavkazskogo Muzeya, 8, 251–266.

Pickrell, J. K., & Pritchard, J. K. (2012). Inference of population splits and mixtures from genome-wide allele frequency data. PLoS Genetics, 8(11), e1002967. https://doi.org/10.1371/journal.pgen.1002967

Pidancier, N., Jordan, S., Luikart, G., & Taberlet, P. (2006). Evolutionary history of the genus Capra (Mammalia, Artiodactyla): discordance between mitochondrial DNA and Y-chromosome phylogenies. Molecular Phylogenetics and Evolution, 40(3), 739–749. https://doi.org/10.1016/j.ympev.2006.04.002

Ramsey, C. B. (2009). Bayesian Analysis of Radiocarbon Dates. Radiocarbon, 51(1), 337–360. https://doi.org/10.1017/S0033822200033865

Reimer, P. J., Austin, W. E. N., Bard, E., Bayliss, A., Blackwell, P. G., Ramsey, C. B., Butzin, M., Cheng, H., Lawrence Edwards, R., Friedrich, M., Grootes, P. M., Guilderson, T. P., Hajdas, I., Heaton, T. J., Hogg, A. G., Hughen, K. A., Kromer, B., Manning, S. W., Muscheler, R., … Talamo, S. (2020). The intcal20 northern hemisphere radiocarbon age calibration curve (0–55 cal kbp). Radiocarbon, 1–33. https://doi.org/10.1017/RDC.2020.41

Rohland, N., Harney, E., Mallick, S., Nordenfelt, S., & Reich, D. (2015). Partial uracil-DNA-glycosylase treatment for screening of ancient DNA. Philosophical Transactions of the Royal Society of London. Series B, Biological Sciences, 370(1660), 20130624. https://doi.org/10.1098/rstb.2013.0624

Ropiquet, A., & Hassanin, A. (2006). Hybrid origin of the Pliocene ancestor of wild goats. Molecular Phylogenetics and Evolution, 41(2), 395–404. https://doi.org/10.1016/j.ympev.2006.05.033

Sarkisov, A. A. (1953). 0 pomesyakh polorogikh [On bovid crossbreeds]. Priroda, 42, 113.

Shackleton, D. M. (1997). Wild sheep and goats and their relatives: status survey and conservation action plan for Caprinae. IUCN.

Supplementary Material: available at https://doi.org/10.17605/OSF.IO/3ECQD

Uerpmann, H.-P. (1987). The ancient distribution of ungulate mammals in the Middle East: fauna and archaeological sites in Southwest Asia and Northeast Africa: Vol. Nr. 27. L. Reichert Verlag.

Weinberg, P. J. (2002). Capra cylindricornis. Mammalian Species, 1–9. https://doi.org/10.2307/0.695.1

Weinberg, P., & Lortkipanidze, B. (2020). IUCN Red List of Threatened Species: Capra cylindricornis. IUCN Red List of Threatened Species.

Wilson, D. E., & Reeder, D. M. (2005). Mammal Species of the World: A Taxonomic and Geographic Reference. JHU Press.

Yang, D. Y., Eng, B., Waye, J. S., Dudar, J. C., & Saunders, S. R. (1998). Technical note: improved DNA extraction from ancient bones using silica-based spin columns. American Journal of Physical Anthropology, 105(4), 539–543. https://doi.org/10.1002/(SICI)1096-8644(199804)105:4<539::AID-AJPA10>3.0.CO;2-1

Zheng, Z., Wang, X., Li, M., Li, Y., Yang, Z., Wang, X., Pan, X., Gong, M., Zhang, Y., Guo, Y., Wang, Y., Liu, J., Cai, Y., Chen, Q., Okpeku, M., Colli, L., Cai, D., Wang, K., Huang, S., … Jiang, Y. (2020). The origin of domestication genes in goats. Science Advances, 6(21), eaaz5216. https://doi.org/10.1126/sciadv.aaz5216

