## Supplemental Information for "A novel lineage of the Capra genus discovered in the Taurus Mountains of Turkey using ancient genomics"

### Supplementary Materials and Methods

#### Direkli Cave

Direkli Cave (37°51'28.08"N, 36°39'13.98"E) is located in the Central Taurus mountains, north-west of Kahramanmaraş Province, near the village of Döngel, Turkey (Erek 2010; Erek 2012). It was discovered and first excavated by Kılıç Kökten in 1959 (Kökten 1960). The cave sits in a south-facing limestone escarpment at an elevation of 1100 metres above sea level. It is located at the base of the Deli Höbek mountain and adjacent peaks which rise up to 2200 metres immediately to the northeast. The cave is located above a small alluvial fan on the eastern side of the north-south trending valley draining the Tekir river (and the D825 highway), itself a tributary to the Ceyhan, a major river system which flows into the Mediterranean. The site is located on the edge of a highly dissected upland and within a corridor providing easy access both to the Mediterranean coast as well as the interior of southern Turkey. It is therefore situated along a convenient route for seasonal movements between higher and lower elevations for both humans and other animal species.

In 2007, new excavations under the direction of C. M. Erek were initiated with the support of the Turkish Ministry of Culture and Tourism in order to further explore the late Pleistocene deposits in the cave, a period that is poorly documented for the central Taurus mountains. Work carried out since that time has identified a stratigraphic sequence rich in artifacts and features dating to the late Pleistocene with lithic parallels both with the Levantine Natufian as well as sites on the Turkish Mediterranean coast. Radiocarbon dates derived from charcoal and animal bone from layers containing microlithic tools, date the main prehistoric occupation layers of the site to between 12,100-8900 years calibrated BCE. The majority of these dates place the occupation in the late Pleistocene prior to the Younger Dryas although the cave was used as late as the late 10th millennium BCE as well.

Excavations were carried out following the natural and cultural stratigraphic layers. Sediments of different colours in each plan square were collected in different buckets and passed through a triple screening system. The sediments, which were all subjected to water screening, were sieved through fine (1mm), medium (2mm) and coarse (4mm) sieves and the materials in each sieve were dried in the shade. These recovery techniques have produced a rich faunal assemblage indicative of the local fauna exploited by the cave's occupants. This faunal community is dominated by the remains of wild goats (*Capra spp*–the focus of this study) but also deer, boar and black bear as well as a variety of fur bearing carnivores.

*Capra* specimens sequenced in this study derive from excavation areas located in the North portion (interior) of the cave (Figure S1). Stratigraphic affiliation of each specimen is reported in Table 1 in the main text. Specimens were recovered from stratigraphic layers 4-8 and all derive from deposits containing material culture (especially microliths) consistent with a late Pleistocene date. In general, *Capra* specimens are highly fragmented and many exhibit cut and percussion marks clearly indicating an anthropogenic origin for the assemblage. There is no evidence that *Capra* remains accumulated through natural processes or carnivore denning behaviours. Direct AMS dates were acquired for two specimens (Direkli4 [Beta-432464:

12130+/-40] and Direkli2 [Beta-425280: 11370+/-40]) confirming their late Pleistocene provenience (12,200-11,200 cal BCE). Four specimens derive from grid square B/13 (Figure S2) including stratigraphic levels 4A, 4C, 7B, 7C providing a temporal sequence from youngest to oldest including Direkli2, Direkli15, Direkli16, Direkli1.

**Sample preparation**

Material from the Muséum national d'Histoire naturelle collections in Paris were sampled on-site. Newly screened specimens from Direkli Cave and Dariali-Tamara Fort were sampled in dedicated ancient DNA facilities in TCD, Dublin, following standard protocols (Pääbo et al. 2004).

Sampled materials were cleaned with a drill bit and then subject to UV for 30 minutes, flipping midway to decontaminate both sides, followed by pulverization using a Mixer Mill (MM 200, Retsch).

**DNA extraction and library preparation**

For MNHN and Tamara Fort material, ~150mg of bone powder was subject to EDTA-prewash and proteinase K 3-day extraction as previously described (Yang et al. 1998; MacHugh et al. 2000; Gamba et al. 2014; Daly et al. 2018). For newly screened Direkli Cave material, samples were subjected to a 0.5% hypochlorite wash prior to EDTA wash and proteinase K-based extraction (Boessenkool et al. 2017; Daly et al. 2021). Previously described protocols for DNA cleanup (Daly et al. 2018; Daly et al. 2021), Uracil-DNA-glycosylase treatment (Rohland et al. 2015) of 16.25ul DNA followed by dsDNA library construction (Meyer and Kircher 2010). Control tubes were included for extraction and library steps and kept for subsequent analysis, but did not show evidence of contamination.

**Screening, sequencing and nuclear genome alignment**

Libraries were amplified (Accuprime Pfx, Thermofisher) with single indexes (MNHN and Tamara Fort specimen) or double indexes (newly-screened Direkli specimen) as described (Daly et al. 2018; Daly et al. 2021). Initial screening was performed using an Illumina MiSeq platform (50bp SE; TrinSeq, Dublin), HiSeq 2000 (100bp SE; Macrogen, Seoul) or NovaSeq 6000 (100bp PE; TrinSeq, Dublin); sample-platform mix is presented in Table S4.

Following de-multiplexing, single-end fastq files were filtered for adaptors which were also trimmed, and reads <30 bp in length removed using cutadapt v1.9.1 (Martin 2011): cutadapt -a AGATCGGAAGAGCACACGTCTGAACTCCAGTCAC -O 1 -m 30.

AdaptorRemoval v2.3.1 (Schubert et al. 2016) was used for pair-end data, for the above QC steps and also to collapse overlapping read pairs: AdapterRemoval -collapse --minadapteroverlap 1 --adapter1 AGATCGGAAGAGCACACGTCTGAACTCCAGTCAC --adapter2 AGATCGGAAGAGCGTCGTGTAGGGAAAGAGTGT --minlength 30 --gzip --trimns --trimqualities

Trimmed (SE) or collapsed (PE) reads were aligned to ARS1 (Bickhart et al. 2017) using bwa aln (Li et al. 2008) relaxing parameters (Meyer et al. 2012) and handling sam/bam files using samtools v1.11 (Li et al. 2009). Read groups were assigned using bwa samse/sampe to differentiate sequencing libraries and PCR reactions. To estimate endogenous DNA %age, duplicate removal was performed using picard v2.22.1 (The Broad Institute 2018) and mapQ ≥30 filtering with samtools, dividing QCed aligned reads by raw reads. Substitutions associated with ancient DNA were examined using mapDamage2 (Jónsson et al. 2013), and are presented in Figure S4.

Samples selected for deeper sequencing (typically >10% endogenous DNA, and including additional sequencing of Direkli4 originally reported in (Daly et al. 2018)) were sequenced on either a Illumina HiSeq 2000 (100bp SE; Macrogen, Seoul) or NovaSeq 6000 (100bp PE; TrinSeq, Dublin). Deeper sequenced data was aligned as above, with a subsequent indel realignment step (The Broad Institute 2018) and softclipping of reads by reducing the base quality of the first and last four bases to zero using a custom python script. Genome depth was estimated using GATK and presented along with alignment statistics in Table S4.

For alignment to the sheep reference, the above pipeline was used substituting ARS1 with the sheep reference Oar_rambouillet_v1.0 available at https://www.ncbi.nlm.nih.gov/assembly/GCF_002742125.1/. Samples aligned to sheep are indicated in Table S3.

**Mitochondrial capture**

Samples not selected for deep genomic sequencing were enriched for mammalian mtDNA sequences using an in-solution RNA hybridization approach using custom baits as previously described (Gnirke et al. 2009; Maricic et al. 2010; O'Sullivan et al. 2016; Daly et al. 2018); Daicel Arbor Biosciences, Ann Arbor, USA). Libraries enriched for mtDNA were subsequently sequenced using a MiSeq platform (50bp SE; TrinSeq, Dublin).

**Mitochondrial genome alignment and phylogeny**

Reads were aligned to circularized (15bp either end) mitochondrial reference genomes as previously described (Daly et al. 2021), realigning to closer mtDNA references to maximize sequence recovery. Sample fasta files were producing using ANGSD v0.922 (-doFasta2 -setMinDepth 3 -minQ 20 -minMapQ 30 -trim 4; (Korneliussen et al. 2014)) and decircularized by removing 15bp at both ends of the resulting fasta sequence. Due to the low coverage of Direkli17, this sample fasta was produced with -setMinDepth 1 and analyzed independently of others.

A multiple sequence alignment of data generated here plus published *Capra* mtDNA sequences (Table S5) was generated using MUSCLE (Edgar 2004). A ML phylogeny with 100 bootstrap replicates was then computed using phyML (Guindon et al. 2010) using a BIC-selected substitution model, and visualized using Figtree (Rambaut 2009).

Pairwise differences among aligned mtDNA sequences was calculated using a custom Python script, ignoring sites where one of the two samples were missing data (“N”). For F lineage comparisons, Direkli17 was excluded due to its low coverage and the higher error rate implicit in the -setMinDepth 1 option used for that sample.

**Modern alignment**

Modern genomes (Table S3) were aligned to ARS1 or Oar_rambouillet_v1.0 using bwa mem v0.7.5a-r405 (Li et al. 2008; Li et al. 2009; Li 2013) and following GATK Best Practices, removing duplicates and reads with mapQ ≥30. Among these were 3 modern Tur genomes, which were included in the IBS calculation and nj tree (below) with permission from the VarGoats consortia (Denoyelle et al. 2021) prior to these samples’ in-depth analyses. Samples aligned to sheep are indicated in Table S3.

**ANGSD-based analyses**

All ANGSD (v0.922; (Korneliussen et al. 2014)) analyses were performed using the following parameters: -minMapQ 30 -minQ 20 -C 50 -remove_bads 1 -uniqueOnly 1 -rmTrans 1 and restricting analyses only to the autosomes. When necessary (-doMajorMinor 5 or -doAbbababa 1), the ancestral sequence was defined using a fasta file from a goat-aligned sheep (Daly et al. 2021). Genotype likelihoods were computed using -GL 2 -rmTrans 1 -SNP_pval 1e-6 -skipTriallelic 1.

**Error estimation**

To estimate the error rates of the Direkli specimen and *Capra* genomes reported here, the ANGSD “perfect genome” approach was employed, using a high coverage Old Irish Goat (“IOG”, ~42X): -doAncError 1 -ref IOG.fa, where IOG.fa was generated using ANGSD -doFasta 1 -doCounts 1 -setMinDepth 13 -setMaxDepth 76 -C 50 -minQ 20 -minMapQ 30. Sheep was used as the outgroup. Error rates were reasonable, with all falling below 0.5% and the highest reaching 0.3428%; the highest Direkli genome error was just 0.1947%. Error rates are shown in Tables S1 and S3 (for previously published Direkli bezoar and the Tur1 specimen).

We additionally assessed error rates of ancient vs modern individuals by plotting distance from the outgroup as calculated from Identity-By-State statistics (see below). We expect high error individuals, particularly ancient ones, to show inflated distances from the outgroup (sheep). Plotting distances (Figure S5), the majority ancient Direkli specimen do not show excessive distance to the outgroup, with the highest distances being observed in historic and modern European (alpine and Iberian) ibex. A single Direkli wild goat / *Capra aegagrus* shows elevated distance to the outgroup relative to modern *Capra aegagrus* (0.240308), but all other Direkli specimens have unremarkable distance values.

**D statistics**

D statistics were computed using -doAbbaBaba 1 -rmTrans 1 -doCounts 1 and using sheep as above to define the ancestral allele. Correlation between *D* statistical tests was measured by calculating Pearson’s correlation coefficient *r* using the cor() function of R (R Core Team 2021). *D* test results are presented in Figure S11 and Tables S7-9. A subset of *D* statistic tests were also repeated using sheep aligned-data (see above); Pearson's $r$ for the test indicated in Table S6-8 were 0.927, 0.950, and 0.494 respectively. The latter is not concerning as the $D$ statistics in question (Direkli4, Tur1; X, Sheep) are overwhelmingly large for goat and sheep-aligned data (i.e. $|D| >> 0.05$) and are consistent in directionality.

**Identity-by-state (IBS)**

Pairwise IBS matrices were computed for reported genomes ≥0.5X coverage, the lower coverage Tur2 genome (*C. cylindricornis*), a subset of modern and ancient *Capra* genomes, and a sheep outgroup using ANGSD (-makeMatrix 1 -doCov 1 -doMaf 1 -minFreq 0.05 -minInd 56), allowing ~5% missingness per site. Neighbour-joining phylogenies were constructed using the nj() function of the ape R package (Paradis and Schliep 2019). Published genomes included in IBS calculations are indicated in Table S3; sheep was used as the outgroup.

To calculate node support, pseudo-bootstrap datasets were generated by sampling-without-replacement 50 5Mb regions a total of 100 times, and calculating IBS values for each of the 100 pseudoreplicates. Neighbour-joining trees were calculated as above and node supports applied to the base tree using RAxML (Stamatakis 2014), and are presented in Figure 1B.

To control for possible reference genome effects, an additional IBS neighbour-joining tree was computed using sheep-aligned data using the following ANGSD settings, allowing for 90% site missingness due to the smaller number of genomes used: -doMajorMinor 4 -GL 2 -minFreq 0.05 -minInd 41. Node support values were calculated as above, using 50 x 5Mb regions per replicate. This tree is presented in Figure S5, with (A) and without (B) the lower coverage Direkli16 sample which falls within the "taurasian tur" clade. Site counts for both trees are 96,930 and 110,672 respectively.

We additionally constructed a MDS plot using the sheep-aligned MDS data and R functions cmdscale(as.dist()) on the IBS .ibsMat file (Figure S6A), with an insert of the corner of the plot containing the taurensis tur lineage. The MDS places the nubian, alpine, and european ibex specimens, as well as tur, in a distinct part of the MDS plot away from a *Capra aegagrus/ hircus* group and a *Capra sibirica/falconeri* (Siberian ibex and markhor). Within the west Eurasian ibex & tur genomes, the relative affinities seen in the nj tree (Figure 1B) are reiterated, with the tur group being closer to the European ibex than to Nubian ibex. The taurasian tur genomes Direkli4 and Direkli16 clearly fall within the Tur group. Both eastern and western tur genomes form distinct groups in MDS space, with Direkli tur showing somewhat higher affinity to the eastern tur group. We additionally computed IBS using sheep-aligned, ancients & historic samples only, allowing 20% missingness to account for the relatively high number of low coverage samples (-minInd 21). From the resulting IBS matrix (computed from 464,799 biallelic transversion sites), an MDS plot was produced (Figure S6B). This plot offers less discriminatory power within the "ibex / tur" group, as expected given the lower number of *Capra* genomes and overall lower coverage among the non *Capra hircus / Capra aegagrus* genomes. However, both Direkli "taurasian tur" genomes fall clearly within the "ibex / tur" group

**Haploid calling**
To produce a haploid (random read sampling) callset of genomes with at least 0.5X coverage for subsequent analyses the ANGSD -doHaplocall function was employed: -doHaploCall 1 -minInd 2 -minQ 20 -minMapQ 30 -remove_bads 1 -uniqueOnly -doCounts 1 -C 50 -trim 4. Output files were then screened using a custom python to retain biallelic sites only and remove CpG sites (232,685,198 remaining), and then converted to plink format (Purcell et al. 2007) using the haploToPlink tool provided with ANGSD. An additional dataset consisting of genomes ≥2X coverage was also generated for Treemix and Orientagraph analyses (below).

**Outgroup/sheep ascertained sites**
To create an outgroup-ascertained set of variant sites for Treemix/Orientagraph, Minor Allele Frequencies were computed dataset of 11 ARS1-aligned sheep (Table S3):
-doMaf 1 -doCounts 1 -minMaf 0.05 -minQ 20 -minMapQ 30 -C 50 -doMajorMinor 3 -GL 2 -remove_bads 1 -uniqueOnly 1 -SNP_pval 1e-6 -setMinDepthInd 4 -minInd 6 -sites sites.txt; where sites.txt was a file describing biallelic sites in the 2X haploid-called dataset (above). These 223,772 outgroup-varying sites were then extracted from the 2X dataset using plink --extract. Twelve modern domestic goat samples (Alashan2, 3, and 4; Erlangshan1, 3, and 4; Aerbasi1, 3 and 6; Liaoning2, 3, and 5; see Table S3) were then removed due to an excess of missingness (>0.15).

**Introgressed regions**
To examine possible sources of 112 haplotypes identified as introgressed in domestic goat (Zheng et al. 2020), IBS values were calculated using ANGSD (-doIBS 1 -doMajorMinor 5 -minInd 157 -minMaf 0.02). The -minInd parameter was set to ensure only sites with ≤10% missing data across the 175 individuals were considered, and -minMaf so that alleles found at least one *Capra* source and two domestic genomes were included. Genomes with ≥1X mean coverage were included, plus the *Capra* samples of interest. Introgressed region coordinates were obtained from Data File S1 of (Zheng et al. 2020).

Heatmaps from pairwise ibs data were constructed using the R gplots heatmap.2() function (Warnes et al. 2019), first removing Nubiana1 and Caucasus1 due to coverage effects, and sheep samples due to distortion of ibs distance value scales. Heatmaps and hierarchical clustering of each putatively-introgressed region are displayed in Figures S19-30. Additionally, nj trees were constructed for each IBS matrix using the nj() function and visually assessed to determine if the Direkli4 lineage is a possible source of introgressed haplogroup. All 10 nj trees are shown in Supplementary Data Files 1 and 2. Assessing nj trees and heatmaps, 3 regions out of the 112 total are plausible as Direkli4 being the closest-to-domestic sample (1:104,150,173-104,349,720; 2:24,324,410-24,369,675; 13:66,710,508-66,749,824); a further 7 may fit with the Direkli4-Tur1 clade being closest to domestic. These assignments should be considered preliminary in lieu of higher quality genomic data for the two Caucasian tur and the Direkli4 lineages.

**Identity of MNHN sample Falconeri1**

Falconeri1, thought to be a *Capra falconeri hepteneri* (Tadjik or Bukharan markhor) from the Muséum national d'Histoire naturelle (MNHN-ZM-AC-2009-243), shows the genetic profile aberrant relative to its supposed species. The mitochondrial sequence of Falconeri1 groups with the Barbary sheep or aoudad (*Ammotragus lervia*), a member of the *Caprini* tribe but not of the genus *Capra* (Figure S8). The mtDNA does not form a clade with the two other markhor sequences included in the phylogeny. Consistent with this is the position of Falconeri1 in IBS-based nj trees, which places the sample basal to all other non-outgroup samples, and does not group with other markhor samples (Figure 1B, Figure S5). We conclude that the sample was misidentified during storage or sampling, and that the genetic sample labelled here as Falconeri1 is more likely to be a Barbary sheep. As such the sample was excluded from subsequent analyses.

**Extended *D*/Direkli4-specific alleles**

We extended the general idea of the *D* statistic (Green et al. 2010; Patterson et al. 2012) and group *D* statistic (Soraggi et al. 2018) to identify variants derived in a genome/population of interest, H3, and ancestral in a set of other defined genomes/populations and an outgroup (i.e. multiple "outgroups", see Figure 3A). The number of times the derived allele (i.e. the H3 "specific" allele) was observed in a reference individual/genome H1 was counted ($nBABA_{ex}$), as was the number of times the derived allele was observed in the target individual/genome H2 ($nABBA_{ex}$). The difference between $nABBA_{ex}$ and $nBABA_{ex}$ was then divided by the sum of $nABBA_{ex}$ and $nBABA_{ex}$, analogous to the *D* formula:

$$D_{ex} = (nABBA_{ex} - nBABA_{ex}) / (nABBA_{ex} + nBABA_{ex})$$

A Z value can then be computed from bootstraps estimate of the standard error using 5Mb genome blocks and 1000 bootstrap replicantes, to normalize the $D_{ex}$ value.

Sites must be covered at least once in each defined group ("outgroups" or otherwise). The ancestral allele was also conditioned on being at 100% frequency in each of the "outgroup" groups, but conceptually this criteria could be slackened to investigate patterns of derived allele sharing (i.e. derived variants in H3 that are present at some frequency e.g. >0%,≤10% in one of the defined "outgroups"). Calculations of $D_{ex}$ were based on the biallelic CpG-removed haploid-called sites (232,685,198 total), computing with and without transitions, and performed using a custom python script.

Initially we examined the sharing of Direkli4-specific variants relative to a population of ~10,000 year old Neolithic domestic-like genomes from the Zagros Mountains (Daly et al. 2021), requiring the Direkli4-derived allele to be fixed for ancestral and covered at least once each in the following groups:

-Direkli bezoar
-Other bezoar (ancient and modern)
-Other *Capra*
-Sheep (using the 11 genomes defined in Table S3) ultimately defining the ancestral state

This would therefore identify variants specific to the Direkli4 lineage, and exclude those shared with the other *Capra* genomes analyzed here (e.g. Caucasian Tur) or the Direkli bezoar (which have likely experienced gene flow with the Direkli4 lineage, see Figure S11). $D_{ex}$ values were highly similar when computed using transversions only (Figure 3B) or all variants (Figure S11, for both see Table S10). To control for reference genome effects we also calculated $D_{ex}$ requiring the Direkli4-derived allele to segregate in sheep, with highly correlated results (Pearson's $r$=0.9935, Table S10).

We repeated the $D_{ex}$ calculation but varying H3 to be a different genome of interest (Figure S13, Table S13). *Capra aegagrus* from Direkli Cave (Direkli1-2, Direkli5, Direkli6) show highest levels of shared-specific ancestry, occurring in European and African goat; this ancestry is only somewhat correlated with Direkli4-specific ancestry ($r$ = 0.6562-0.5731) and likely reflects gene flow from Anatolian bezoar to the ancestors of west Eurasian domestic goat (e.g. Figure S15, S15). A single markhor (Cfalconeri04, SAMN10736157) may have east Asian related gene flow in its ancestry, while derived alleles signals of multiple *Capra* genomes in sub-Saharan goat imply gene flow from a *Capra* source into these domestic populations.

We observed positive $D_{ex}$ values for Direkli4 and ancient west Eurasian goats, but not modern goats (Figure 3B). To assess whether technical biases may artificially cause Direkli4 to share more alleles with ancient samples, we examined the correlation with $D_{ex}$ when other *Capra* genomes define the "specific" allele (Figure S13, Table S12). Correlations of specific allele sharing values with *C. caucasica*, the Palaeolithic Armenian bezoar Hovk1, and medieval *C. cylindricornis* Caucasus1 were highest (Pearson's $r$ = 0.9258, 0.9231, 0.8824 respectively) (Table S12). The relatively high range of correlations with historic *Capra* genomes ($r$ = 0.7194-0.8363) suggests some technical bias, but does not completely explain the pattern of Direkli4-specific allele sharing.

As mentioned above this "extended *D*" approach was amenable to allowing the H3-specific variant to also segregate at a defined rate among "outgroups", effectively "disentangling" patterns of shared derived alleles. We investigated Direkli4-specific allele sharing, cycling through different target H2 genomes, for variants also >0%, ≤10% across "outgroups" (but fixed as ancestral in sheep). For each target H2, the total number of times a given "Outgroup" individual shared the "Direkli4-specific" allele was recorded, expressing as a proportion of the total >0%, ≤10% shared variants in a heatmap format (Figure S14). While sensitive to coverage (as deeper sequenced samples will on-average make up a greater proportion of the total number of shared "Direkli4-specific" variants), differences are apparent between Direkli4-specific allele sharing between ancient and modern European goat; the former share more Direkli4 alleles with Direkli bezoar, while the latter share more Direkli4 alleles with the Caucasian tur genomes Tur1 and Caucasus1. This pattern could be explained by a turnover in European goat populations to one with a greater Caucasian tur-related ancestry, a technical bias that increases affinity between the Direkli bezoar and ancient European goat or Tur genomes and modern European goat, or gene flow between a population related to modern European goat and Caucasian tur. These hypotheses could be tested with finer temporal sampling of European goats, or a time series of Caucasian tur to determine if gene flow from domestic populations occurred.

Python scripts to run the extended *D* calculations and to “disentangle” the pattern of derived allele sharing are available on github.com:
https://github.com/Xevkin/direkli_caprid_extended_ds/tree/main

To confirm our results were unlikely due to choice of reference genome, we recalculated a subset of $D_{ex}$ statistics with data aligned to sheep, with and without ascertaining in the sheep outgroup population and examining tests related to Direkli4-specific variants in domestic groups (Tables S10 and S11). Correlation with goat-aligned data was high ($r$ = 0.897 and 0.906 respectively using transversions-only), suggesting that a reference bias towards ARS1 was unlikely driving the observed patterns in Direkli4-sharing allele sharing. However, some Z scores differed by crossing the chosen significance threshold (| Z | > 3) while retaining their $D_{ex}$ directionality. For example, with sheep-ascertained sites the modern European goats IOG and Italian4 obtain significant negative scores (having fewer Direkli4-specific variants than the reference Neolithic Zagros group) when using sheep-aligned data but not goat aligned (negative but not |Z| > 3, Table S11). This is likely due to the lower number of genomes included in the $D_{ex}$ calculation when using sheep-aligned data, a necessity due to the computational limitations of aligning available data to the sheep genome. Each additional genome include in the H4 “outgroup” (e.g. diverse bezoar, *Capra* specimen, other Direkli bezoar) should reduce the total number of sites in the $D_{ex}$ calculation, by filtering out variants otherwise assigned as “Direkli4-specific”. Fewer sites will reduce the sensitivity of the test (by decreasing the number of sites included in each bootstrap iteration) while increasing the specificity of the variants. As such it is the consistency of directionality and correlation of the $D_{ex}$ values between different data sets which should lend confidence to the goat-aligned results presented in Figures S12 and S13.

**Graph-based modelling**

To generate a graph-based admixture model of how individual genomes relate, Treemix (Pickrell and Pritchard 2012) was employed on the 2X-sheep ascertained dataset. The number of migration events *m* was varied from 0 to 5, with other parameters set as -root Sheep -k 1000 --noss --global. Node support values were estimated using 50 bootstrap replicates and the -boot option, applying node support values to the base tree using RAxML (Stamatakis 2014).
To maximize the possibility of detecting gene flow between the Direkli4/Tur lineage, Orientagraph (Molloy et al. 2021) was employed on a reduced set of populations and at a group level. Sites were filtered to retain those with at least one call per group and a MAF of ≥0.05, leaving 104,550 sites. *m* was varied from 0 to 4 due to computational limitations, running with Treemix settings as above but with the addition of -mlno to find the Maximum Likelihood Network Orientation, and *k* of 500 due to the lower SNP number.

Finally ADMIXTOOLS2 (Maier et al. 2022) was employed to explore admixture graph space in a complementary manner. To investigate the admixture status of the “taurasian tur” lineage in a limited graph space, 6 populations/genomes were included:

1. West Caucasus Tur *Capra caucasica* (**Tur1**)
2. Direkli4 (**DIR4**)
3. Direkli bezoar / Epipaleolithic Taurus (**ETa**)
4. PPN/PN East Iran (**NEI**)
5. Neolithic Serbia (**NSe**)
6. A group of ARS1-aligned sheep were used as the outgroup (**Sheep**).

SNPs used were the 223,772 sheep-ascertained variants described above. We followed the approach suggested by (Maier et al. 2022), fitting 50 graphs per complexity class (m=0-5) using find_graphs(stop_gen = 100, outpop = 'Sheep') using pre-computed $f_2$ statistics calculated over all goat autosomes and allowing missingness (extract_f2(maxmiss = 1, auto_only= F)). At m=2 (two admixture edges), the preponderance of graphs fit with a worst $f_4$ outlier $| Z | < 3$.

find_graph() was rerun at m=2 for 50 iterations, with the constraint that the Neolithic Serbian group had to receive at least one admixture edge in its history, reflecting the established gene flow event from a population related to the Direkli bezoar (Daly et al. 2018; Daly et al. 2021). Duplicate graphs were removed, as were graphs with 100%/0% admixture events. The best fitting graph was compared to the remaining graphs by calculating bootstrap values for graph scores (qpgraph_resample_multi(nboot=100) and compare_fits()$p_emp), with significantly worse fitting graphs removed; several as-good-as fitting graphs retained significant $f_4$ outlier $|Z| > 3$.

The remaining 11 graphs were then scored for the presence of 1) admixture edges from ETa into DIR4, an indication that the taurasian tur group has bezoar admixture, and 2) admixture edges from DIR4 into ETa, indicating that Direkli bezoar have ancestry related to the taurasian tur. In a majority of graphs (6/11), ETa is modelled as having DIR4 ancestry (median 1.5% DIR4 ancestry, mean 5.2%). Only one graph models DIR4 as a mixture of an ETa related clade and Tur1 (2% for the latter). Following the methodology suggested by (Maier et al. 2022), we examined the best fitting graph at an additional complexity class (m=3) and found ETa again to be modelled as containing DIR4 ancestry (1%), demonstrating this feature to be a robust one. These results suggest that while the data does not exclude bezoar to taurasian tur gene flow (which is implied by other results, including mtDNA lineages (Figure S8)), “Direkli taurasian tur” to “Direkli bezoar” admixture was of greater consequence, with more graphs indicating Direkli bezoar received taurasian tur admixture.

All 12 graphs “as good as” the best fitting graph for m=2 (displayed in Figure S18) are available at https://osf.io/3ecqd/, along with the distribution of log likelihood scores for m=0-5 and the best fitting graph for m=3.

References

Açıkkol A. 2006. Üçağızlı mağarası Faunasının zooarkeolojik açıdan analizi: Capra, Capreolus, Dama ve Cervusların morfometrik açıdan analiz.Güleç E, editor.

Bickhart DM, Rosen BD, Koren S, Sayre BL, Hastie AR, Chan S, Lee J, Lam ET, Liachko I, Sullivan ST, et al. 2017. Single-molecule sequencing and chromatin conformation capture enable de novo reference assembly of the domestic goat genome. *Nat. Genet.* 49:643–650. http://dx.doi.org/10.1038/ng.3802

Boessenkool S, Hanghøj K, Nistelberger HM, Der Sarkissian C, Gondek AT, Orlando L, Barrett JH, Star B. 2017. Combining bleach and mild predigestion improves ancient DNA recovery from bones. *Mol. Ecol. Resour.* 17:742–751. http://dx.doi.org/10.1111/1755-0998.12623

Bökönyi S. 1977. The animal remains from four sites in the Kermanshah Valley, Iran. Oxford: British Archaeological Reports

Churcher CS. 1994. The vertebrate fauna from the Natufian level at Jebel es-Saaïdé (Saaïdé II), Lebanon. *Paléorient* 20:35–58. http://www.jstor.org/stable/41492588

Daly KG, Maisano Delser P, Mullin VE, Scheu A, Mattiangeli V, Teasdale MD, Hare AJ, Burger J, Verdugo MP, Collins MJ, et al. 2018. Ancient goat genomes reveal mosaic domestication in the Fertile Crescent. *Science* 361:85–88. http://dx.doi.org/10.1126/science.aas9411

Daly KG, Mattiangeli V, Hare AJ, Davoudi H, Fathi H, Doost SB, Amiri S, Khazaeli R, Decruyenaere D, Nokandeh J, et al. 2021. Herded and hunted goat genomes from the dawn of domestication in the Zagros Mountains. *Proc. Natl. Acad. Sci. U. S. A.* 118. https://www.pnas.org/content/118/25/e2100901118

Denoyelle L, Talouarn E, Bardou P, Colli L, Alberti A, Danchin C, Del Corvo M, Engelen S, Orvain C, Palhière I, et al. 2021. VarGoats project: a dataset of 1159 whole-genome sequences to dissect Capra hircus global diversity. *Genet. Sel. Evol.* 53:86. http://dx.doi.org/10.1186/s12711-021-00659-6

Edgar RC. 2004. MUSCLE: multiple sequence alignment with high accuracy and high throughput. *Nucleic Acids Res.* 32:1792–1797. http://dx.doi.org/10.1093/nar/gkh340

Erek CM. 2010. A new Epi-Paleolithic Site in the Northeast Mediterranean region: Direkli Cave (Kahramanmaraş, Turkey). *Adalya*:1–17. https://app.trdizin.gov.tr/makale/TVRFNE16Z3hNUT09/a-new-epi-paleolithic-site-in-the-northeast-mediterranean-region-direkli-cave-kahramanmaras-turkey-

Erek CM. 2012. Güneybatı Asya ekolojik nişi içinde Direkli Mağarası Epi-paleolitik buluntularının değerlendirilmesi. *Anadolu / Anatolia*:53–66. https://dergipark.org.tr/tr/doi/10.1501/andl_0000000393

Gamba C, Cristina G, Jones ER, Teasdale MD, McLaughlin RL, Gloria G-F, Valeria M, László D, Ivett K, Ildikó P, et al. 2014. Genome flux and stasis in a five millennium transect of European prehistory. *Nat. Commun.* 5:5257. http://dx.doi.org/10.1038/ncomms6257

Gnirke A, Melnikov A, Maguire J, Rogov P, LeProust EM, Brockman W, Fennell T, Giannoukos G, Fisher S, Russ C, et al. 2009. Solution hybrid selection with ultra-long oligonucleotides for massively parallel targeted sequencing. *Nat. Biotechnol.* 27:182–189. http://dx.doi.org/10.1038/nbt.1523

Green RE, Krause J, Briggs AW, Maricic T, Stenzel U, Kircher M, Patterson N, Li H, Zhai W, Fritz MH-Y, et al. 2010. A draft sequence of the Neandertal genome. *Science* 328:710–722. http://dx.doi.org/10.1126/science.1188021

Guindon S, Dufayard J-F, Lefort V, Anisimova M, Hordijk W, Gascuel O. 2010. New algorithms and methods to estimate maximum-likelihood phylogenies: assessing the performance of PhyML 3.0. *Syst. Biol.* 59:307–321. http://dx.doi.org/10.1093/sysbio/syq010

Hesse BC. 1978. Evidence for husbandry from the early Neolithic site of Ganj Dareh in western Iran. https://www.worldcat.org/title/evidence-for-husbandry-from-the-early-neolithic-site-of-ganj-dareh-in-western-iran/oclc/16112914?referer=di&ht=edition

Horwitz LK. 2003. The Neolithic fauna. In: Khalaily H, Marder O, editors. The Neolithic Site of Abu Gosh. The 1995 Excavations. Vol. 19. Jerusalem: Israel Antiquities Authority Reports. p. 87–101.

Jónsson H, Ginolhac A, Schubert M, Johnson PLF, Orlando L. 2013. mapDamage2.0: fast approximate Bayesian estimates of ancient DNA damage parameters. *Bioinformatics* 29:1682–1684. http://dx.doi.org/10.1093/bioinformatics/btt193

Kersten AMP. 2020. Age and sex composition of Epipalaeolithic fallow deer and wild goat from Ksar ‘Akil. *Palaeohistoria*:119–131. https://www.taylorfrancis.com/chapters/edit/10.1201/9781003079446-8/age-sex-composition-epipalaeolithic-fallow-deer-wild-goat-ksar-akil-kersten

Kökten K. 1960. Anadolu Maras Vilayetinde tarihten dip tarihe gidis. *Türk Arkeoloji Dergisi*:42–53.

Korneliussen TS, Albrechtsen A, Nielsen R. 2014. ANGSD: Analysis of Next Generation Sequencing Data. *BMC Bioinformatics* 15:356. http://dx.doi.org/10.1186/s12859-014-0356-4

Li H. 2013. Aligning sequence reads, clone sequences and assembly contigs with BWA-MEM. *arXiv [q-bio.GN]*. http://arxiv.org/abs/1303.3997

Li H, Handsaker B, Wysoker A, Fennell T, Ruan J, Homer N, Marth G, Abecasis G, Durbin R, 1000 Genome Project Data Processing Subgroup. 2009. The Sequence Alignment/Map format and SAMtools. *Bioinformatics* 25:2078–2079. http://dx.doi.org/10.1093/bioinformatics/btp352

Li H, Ruan J, Durbin R. 2008. Mapping short DNA sequencing reads and calling variants using mapping quality scores. *Genome Res.* 18:1851–1858. http://dx.doi.org/10.1101/gr.078212.108

MacHugh DE, Edwards CJ, Bailey JF, Bancroft DR, Bradley DG. 2000. The extraction and analysis of ancient DNA from bone and teeth: a survey of current methodologies. *Anc. Biomol.* 3:81–103. http://www.academia.edu/download/45484488/The_extraction_and_analysis_of_ancient_D

20160509-18037-1un8a5m.pdf

Maier R, Flegontov P, Flegontova O, Changmai P, Reich D. 2022. On the limits of fitting complex models of population history to genetic data. *bioRxiv*:2022.05.08.491072. https://www.biorxiv.org/content/10.1101/2022.05.08.491072v2

Maricic T, Whitten M, Pääbo S. 2010. Multiplexed DNA sequence capture of mitochondrial genomes using PCR products. *PLoS One* 5:e14004. http://dx.doi.org/10.1371/journal.pone.0014004

Martin M. 2011. Cutadapt removes adapter sequences from high-throughput sequencing reads. *EMBnet.journal* 17:10–12. http://journal.embnet.org/index.php/embnetjournal/article/view/200/479%C3%AF%C2%BB%C2%BF

Masseti M. 2009. The wild goatsCapra aegagrusErxleben, 1777 of the Mediterranean Sea and the Eastern Atlantic Ocean islands. *Mamm. Rev.* 39:141–157. https://onlinelibrary.wiley.com/doi/10.1111/j.1365-2907.2009.00141.x

Meyer M, Kircher M. 2010. Illumina sequencing library preparation for highly multiplexed target capture and sequencing. *Cold Spring Harb. Protoc.* 2010:db.prot5448. http://dx.doi.org/10.1101/pdb.prot5448

Meyer M, Kircher M, Gansauge M-T, Li H, Racimo F, Mallick S, Schraiber JG, Jay F, Prüfer K, de Filippo C, et al. 2012. A high-coverage genome sequence from an archaic Denisovan individual. *Science* 338:222–226. http://dx.doi.org/10.1126/science.1224344

Molloy EK, Durvasula A, Sankararaman S. 2021. Advancing admixture graph estimation via maximum likelihood network orientation. *Bioinformatics* 37:i142–i150. http://dx.doi.org/10.1093/bioinformatics/btab267

O'Sullivan NJ, Teasdale MD, Mattiangeli V, Maixner F, Pinhasi R, Bradley DG, Zink A. 2016. A whole mitochondria analysis of the Tyrolean Iceman's leather provides insights into the animal sources of Copper Age clothing. *Sci. Rep.* 6:31279. http://dx.doi.org/10.1038/srep31279

Pääbo S, Poinar H, Serre D, Jaenicke-Despres V, Hebler J, Rohland N, Kuch M, Krause J, Vigilant L, Hofreiter M. 2004. Genetic analyses from ancient DNA. *Annu. Rev. Genet.* 38:645–679. http://dx.doi.org/10.1146/annurev.genet.37.110801.143214

Paradis E, Schliep K. 2019. ape 5.0: an environment for modern phylogenetics and evolutionary analyses in R. *Bioinformatics* 35:526–528. http://dx.doi.org/10.1093/bioinformatics/bty633

Patterson N, Moorjani P, Luo Y, Mallick S, Rohland N, Zhan Y, Genschoreck T, Webster T, Reich D. 2012. Ancient admixture in human history. *Genetics* 192:1065–1093. http://dx.doi.org/10.1534/genetics.112.145037

Pickrell JK, Pritchard JK. 2012. Inference of population splits and mixtures from genome-wide allele frequency data. *PLoS Genet.* 8:e1002967. http://dx.doi.org/10.1371/journal.pgen.1002967

Purcell S, Neale B, Todd-Brown K, Thomas L, Ferreira MAR, Bender D, Maller J, Sklar P, de

Bakker PIW, Daly MJ, et al. 2007. PLINK: a tool set for whole-genome association and population-based linkage analyses. *Am. J. Hum. Genet.* 81:559–575. http://dx.doi.org/10.1086/519795

Rambaut A. 2009. FigTree. http://tree.bio.ed.ac.uk/software/figtree/

R Core Team. 2021. R: A Language and Environment for Statistical Computing. Vienna, Austria: R Foundation for Statistical Computing https://www.R-project.org/

Rivals F. 2004. Les petits bovidés (Caprini et Rupicaprini) pléistocènes dans le bassin méditerranéen et le Caucase: Etude paléontologique, biostratigraphique, archéozoologique et paléoécologique. http://dx.doi.org/10.30861/9781841716725

Rohland N, Harney E, Mallick S, Nordenfelt S, Reich D. 2015. Partial uracil-DNA-glycosylase treatment for screening of ancient DNA. *Philos. Trans. R. Soc. Lond. B Biol. Sci.* 370:20130624. http://dx.doi.org/10.1098/rstb.2013.0624

Schubert M, Lindgreen S, Orlando L. 2016. AdapterRemoval v2: rapid adapter trimming, identification, and read merging. *BMC Res. Notes* 9:88. http://dx.doi.org/10.1186/s13104-016-1900-2

Soraggi S, Wiuf C, Albrechtsen A. 2018. Powerful Inference with the D-Statistic on Low-Coverage Whole-Genome Data. *G3* 8:551–566. http://dx.doi.org/10.1534/g3.117.300192

Stamatakis A. 2014. RAxML version 8: a tool for phylogenetic analysis and post-analysis of large phylogenies. *Bioinformatics* 30:1312–1313. https://academic.oup.com/bioinformatics/article/30/9/1312/238053

The Broad Institute. 2018. Picard Tools. The Broad Institute https://broadinstitute.github.io/picard/

Warnes GR, Bolker B, Bonebakker, Robert LG, Liaw WHA, Lumley T, Maechler M, Magnusson A, Moeller S, Schwartz M, et al. 2019. gplots: Various R Programming Tools for Plotting Data. https://CRAN.R-project.org/package=gplots

Yang DY, Eng B, Waye JS, Dudar JC, Saunders SR. 1998. Technical note: improved DNA extraction from ancient bones using silica-based spin columns. *Am. J. Phys. Anthropol.* 105:539–543. http://dx.doi.org/10.1002/(SICI)1096-8644(199804)105:4<539::AID-AJPA10>3.0.CO;2-1

Zheng Z, Wang X, Li M, Li Y, Yang Z, Wang X, Pan X, Gong M, Zhang Y, Guo Y, et al. 2020. The origin of domestication genes in goats. *Sci Adv* 6:eaaz5216. http://dx.doi.org/10.1126/sciadv.aaz5216

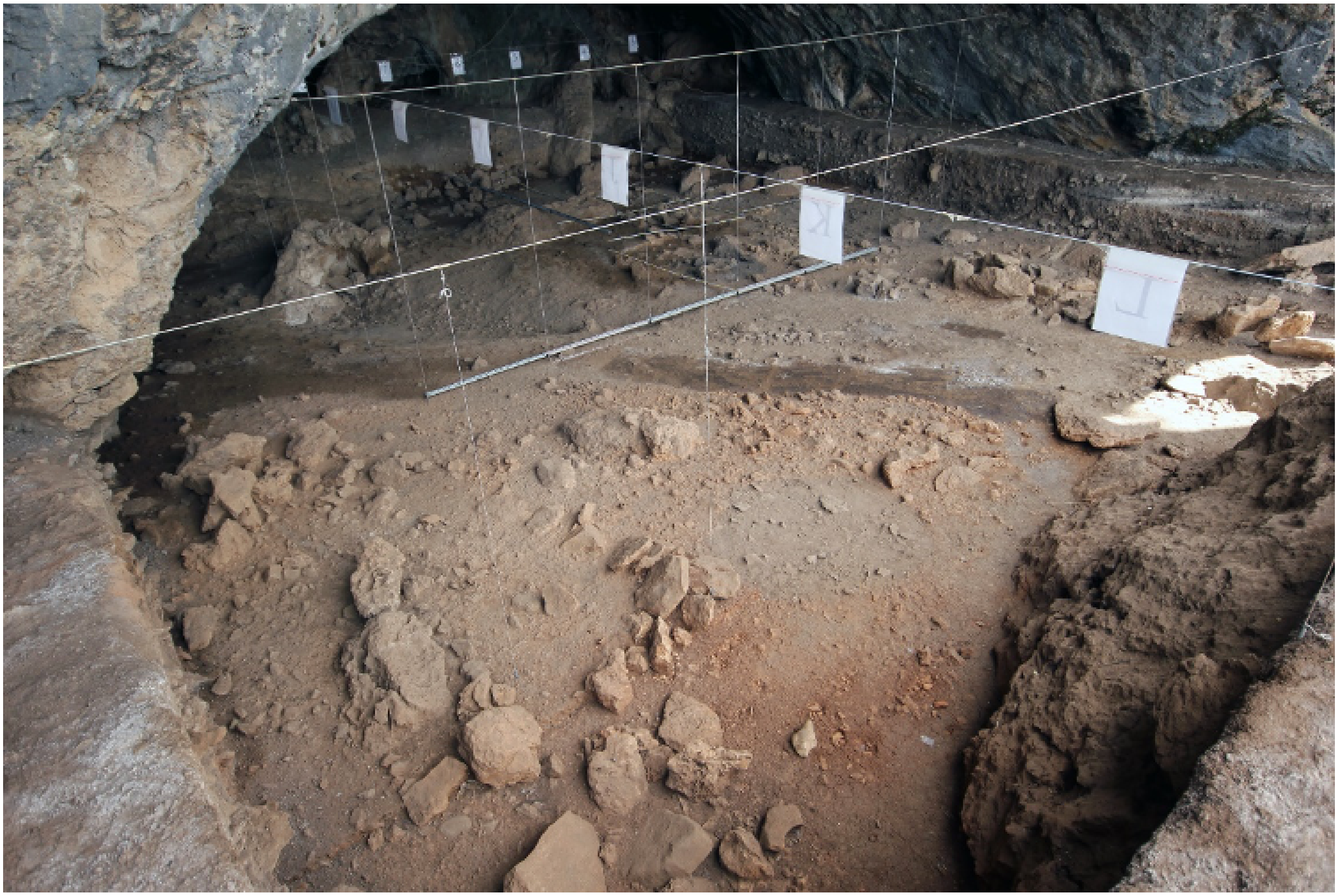

**Figure S1:** View of excavation area from SW, Direkli Cave (from Direkli Cave Excavation Archive, 2018).

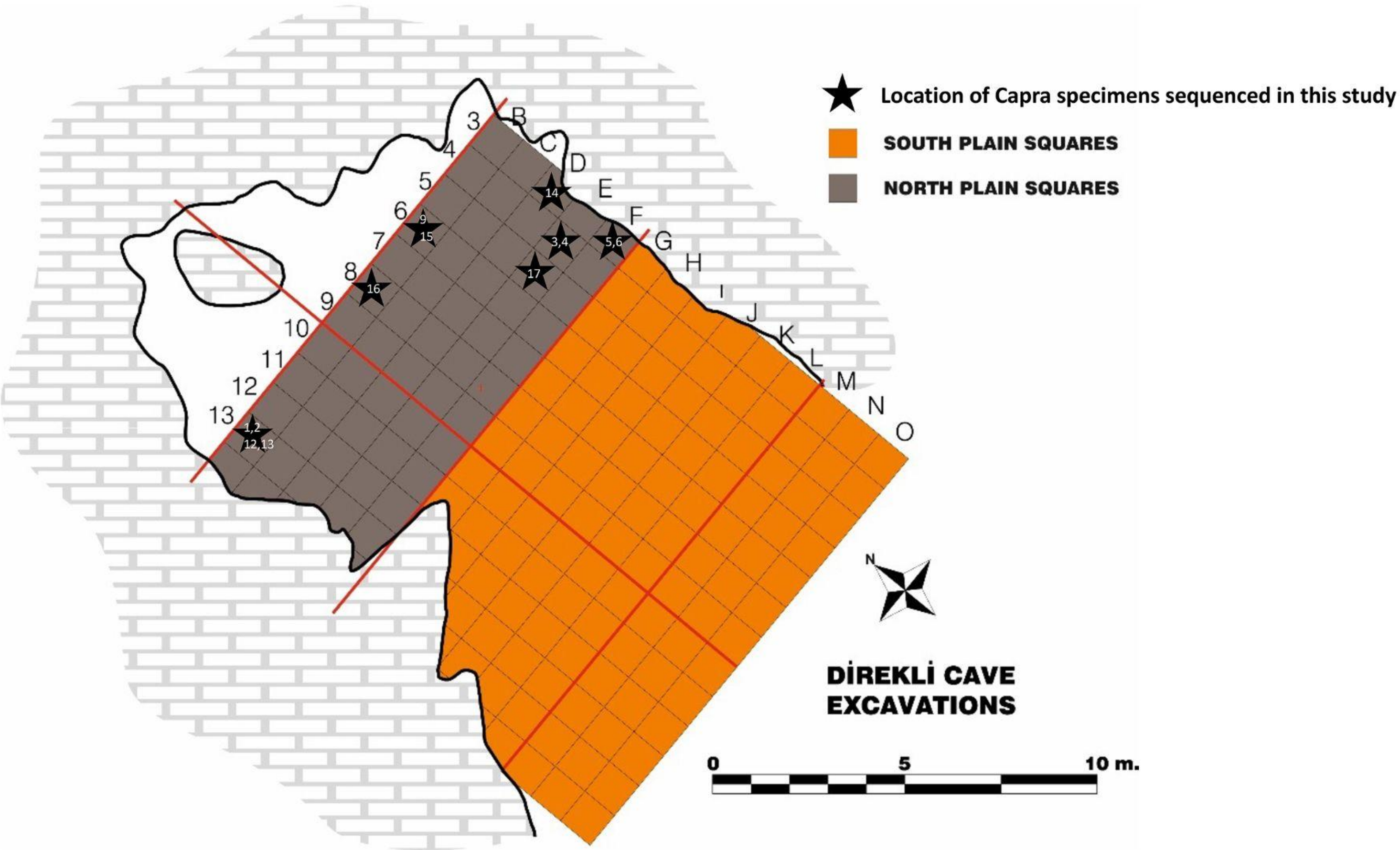


**Figure S2:** Plan map of Direkli Cave showing location of *Capra* specimens sequenced in this study.

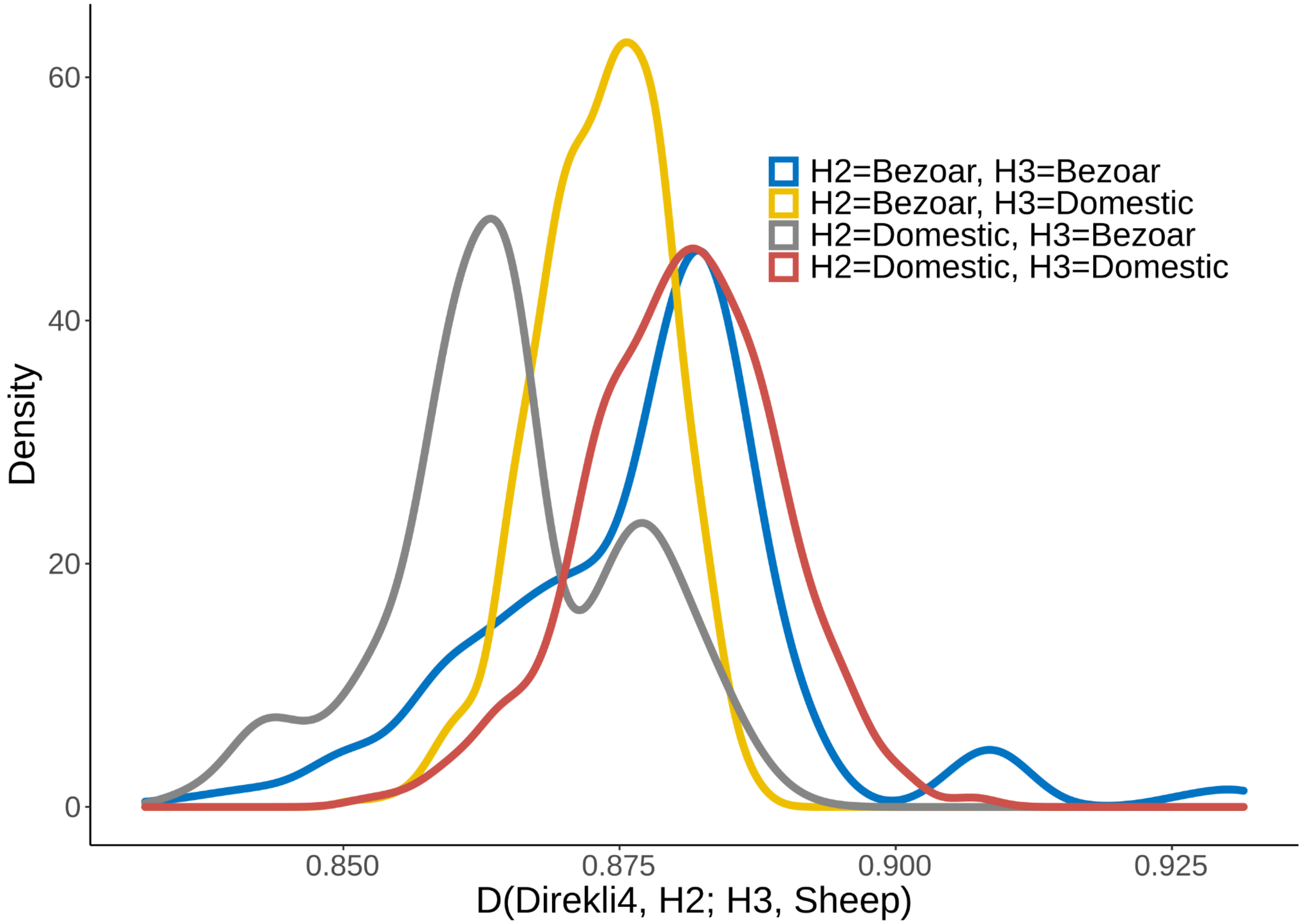


**Figure S3:** Probability density distribution of pairwise *D* statistics using 28 domestic *C. hircus* and 20 wild *C. aegagrus* in the form *D*(Direkli4, H2; H3, Sheep). In all tests positive *D* statistics are obtained, implying an excess of ancestral variation in Direkli4 relative to *C. hircus* and *C. aegagrus*.

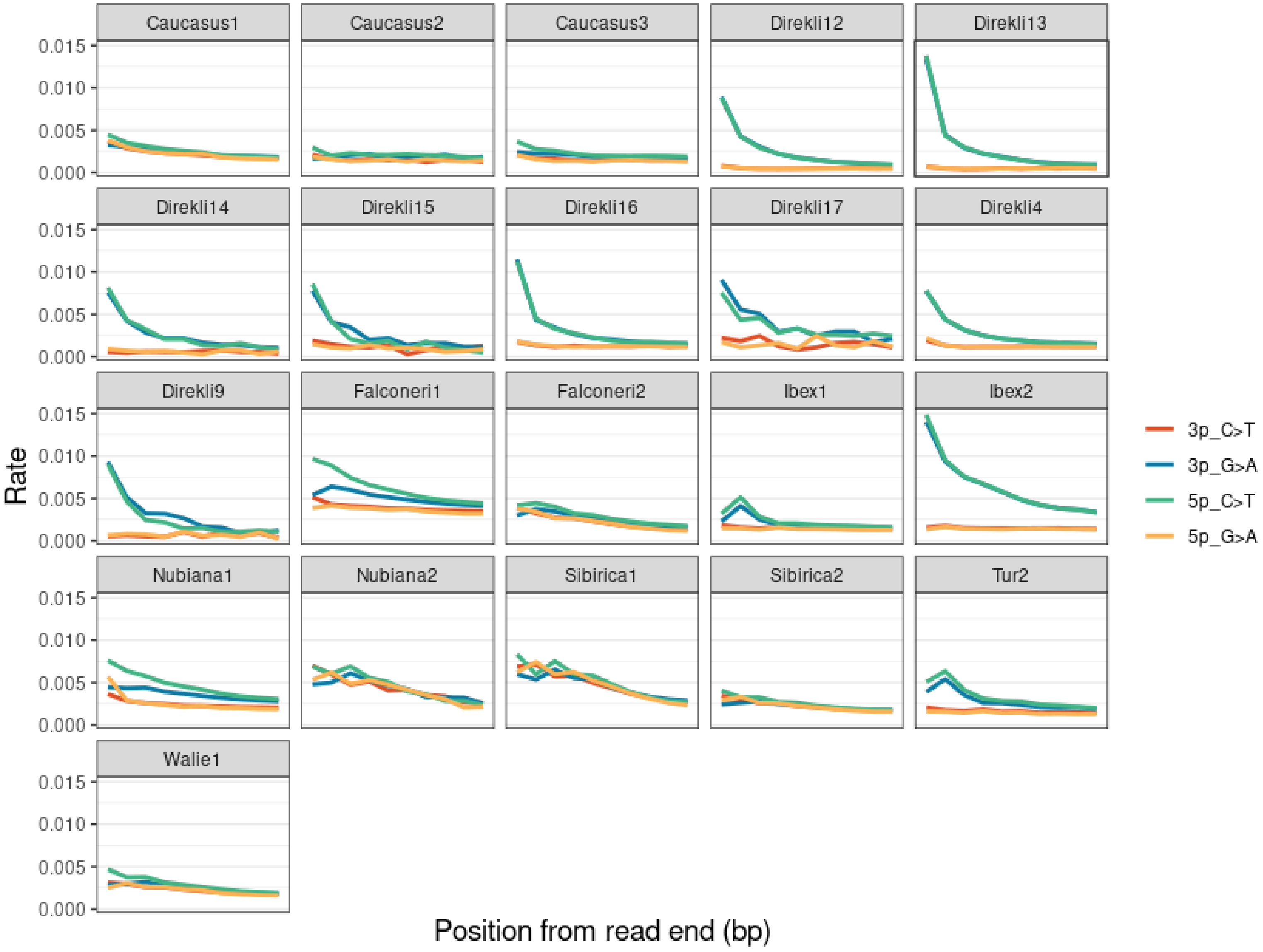


**Figure S4:** Substitution rates of C>T and G>A transitions for ancient and historic samples sequenced in the present study, relative to the 5' and 3' ends of DNA fragments.

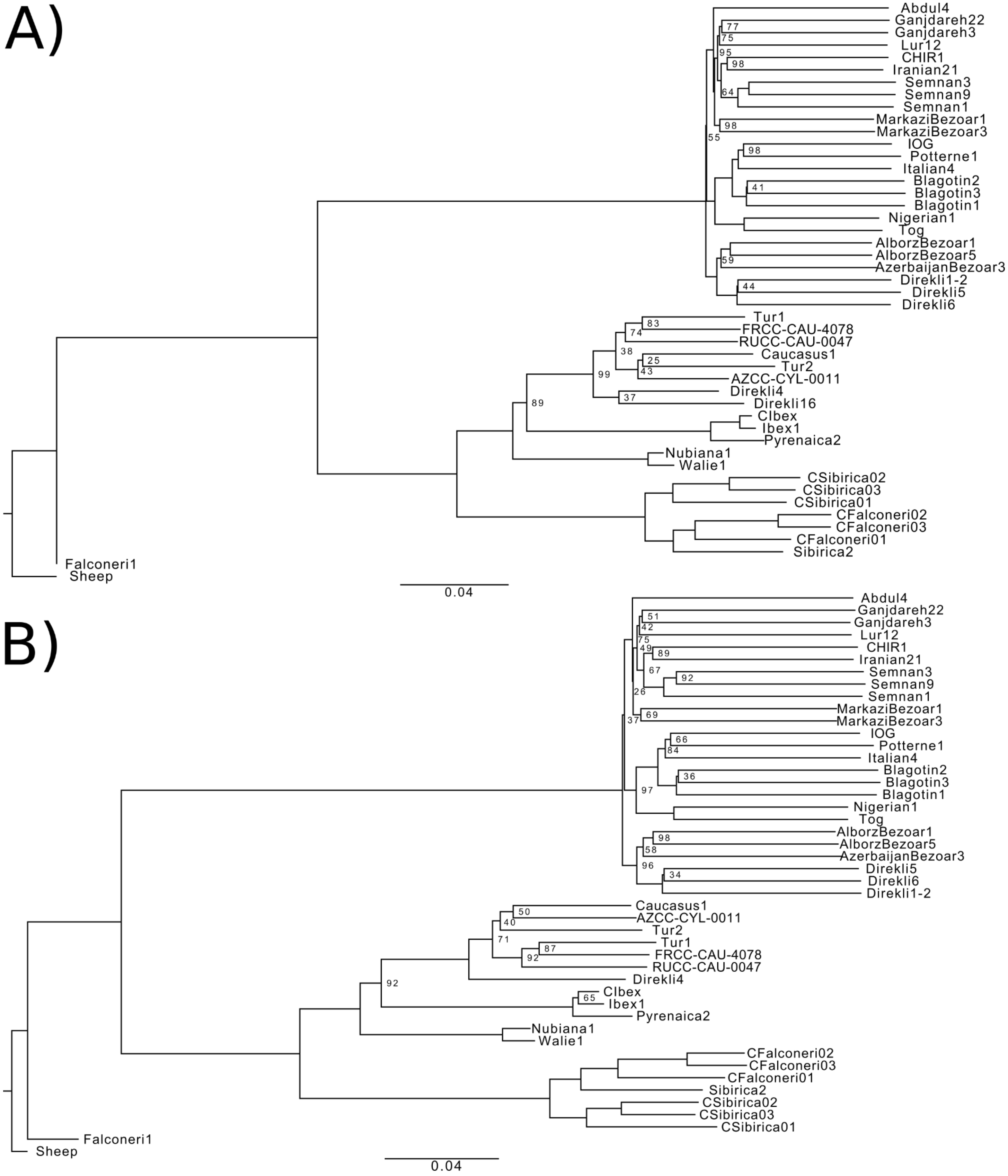


**Figure S5:** NJ Phylogeny using sheep-aligned identity-by-state data, A) with and B) without the lower coverage Direkli16 sample. 96,930 and 110,672 transversion biallelic SNPs were used in each IBS calculation respectively. Node support values are displayed when < 100.

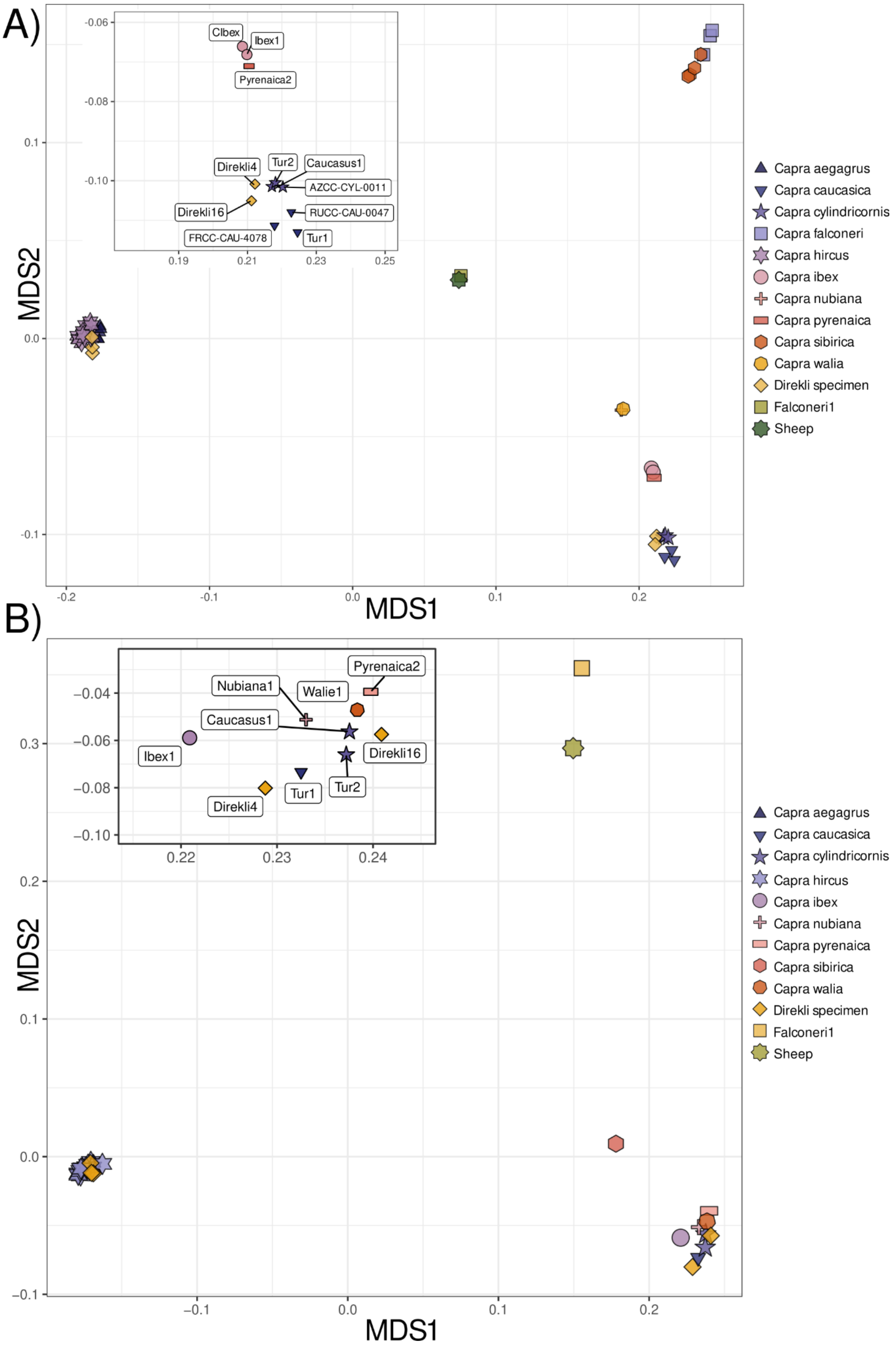


**Figure S6:** MDS plot of sheep-aligned IBS data A) underlying Figure S5A and B) using ancients & historic samples only. Insert plot gives a higher detailed view of the MDS region encapsulating the european ibex and tur specimen, including two “taurasian tur” (Direkli4 and Direkli16).

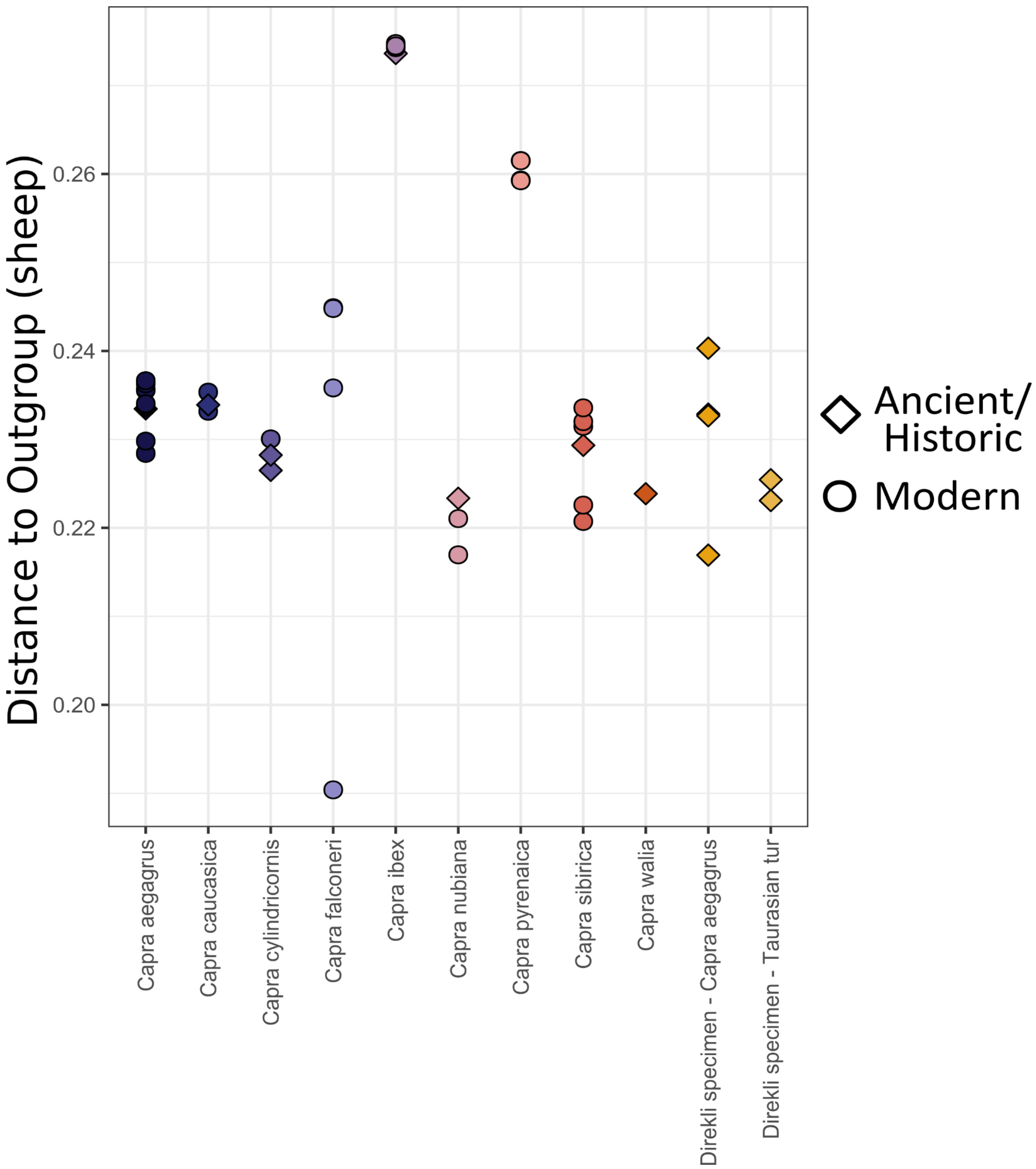


**Figure S7:** Distance-from-the-outgroup for modern and ancient/historic *Capra* genomes. Distance values are obtained from sheep-aligned IBS statistics underlying Figure S5A.

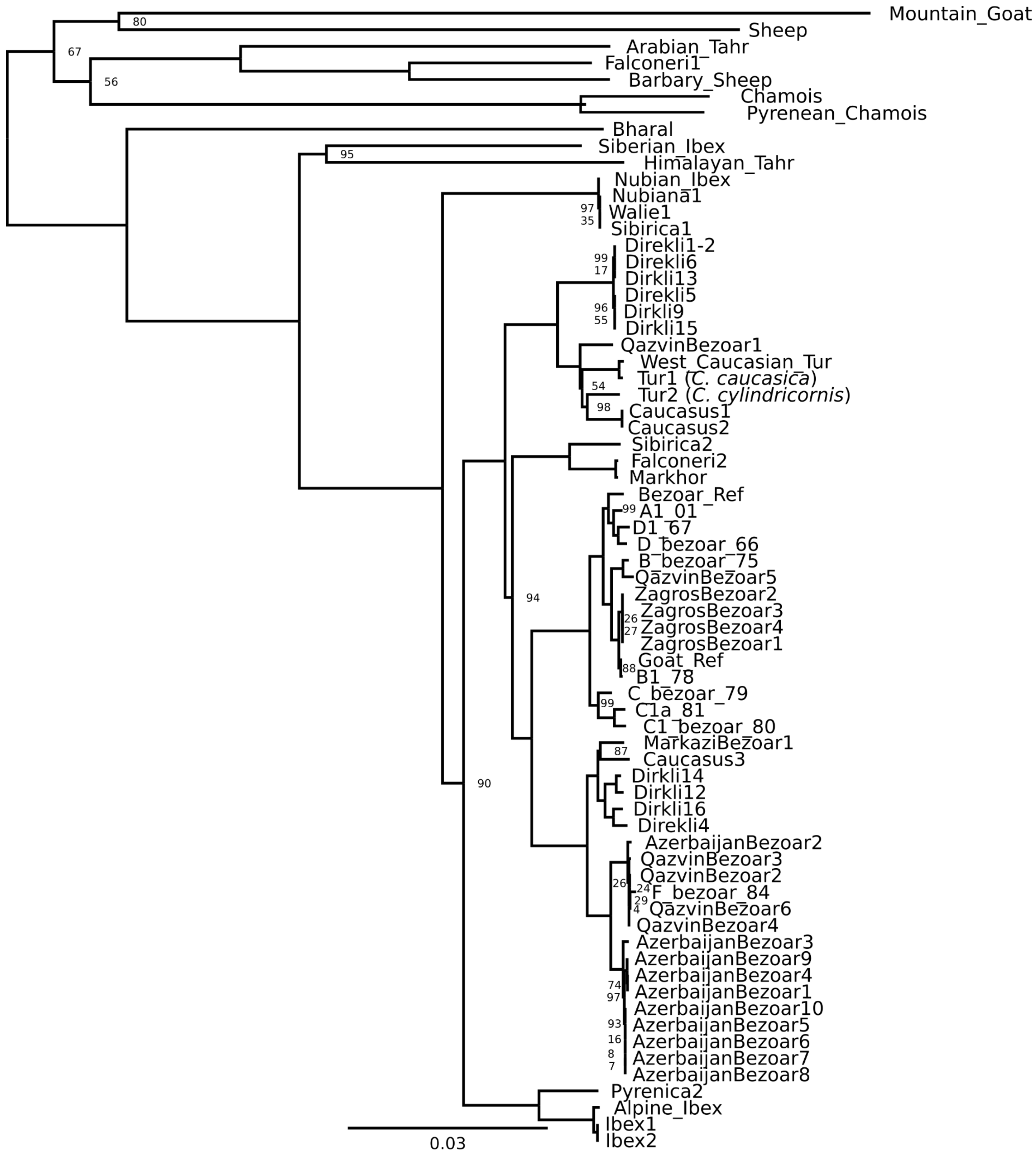


**Figure S8. ML phylogeny of mtDNA, uncollapsed.** Bootstrap node support values (100 replicates) are displayed when <100. Outgroup is Yak and not shown.

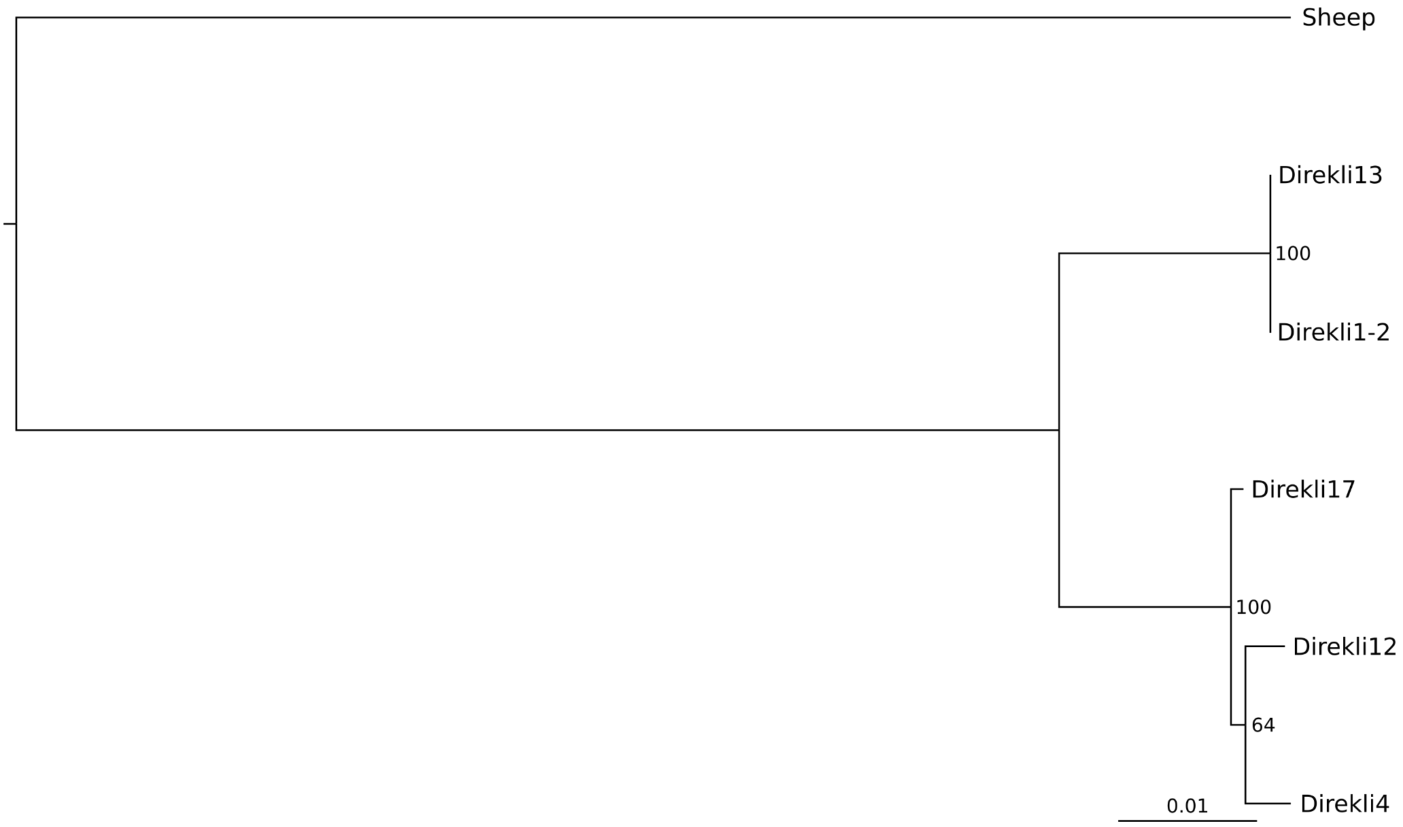


**Figure S9: Reduced mtDNA ML phylogeny including Direkli17.** Direki17's low number of called sites (1162) precluded inclusion in a larger phylogeny; Direkli1-2 and Direkli13 represent the "T" clade, while Direkli4 and Direkli12 represent the "F" clade. Bootstrap values (100 replicated) displayed at nodes.

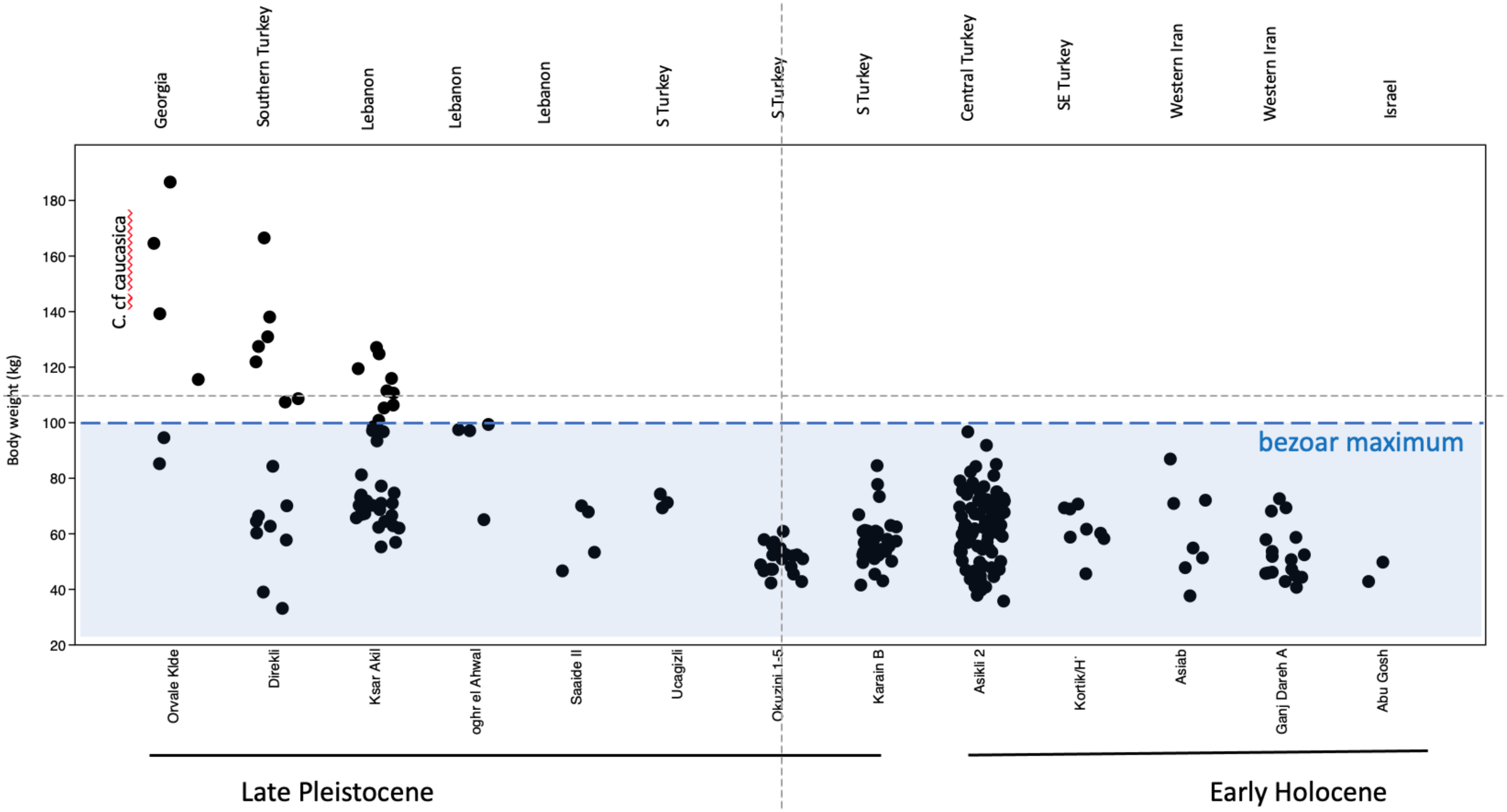


**Figure S10:** *Capra* body size estimates from archaeological assemblages, organized by region, time, and archaeological site. Body weight estimates are based on Rivals' (Rivals 2004) conversion formula of astragalus measurements to body weight (kg). The bezoar (*Capra aegagrus*) body size maximum of ~100kg (Masseti 2009) is highlighted to demonstrate the high upper range and variability observed at Direkli Cave and also Ksar Akil. Data from authors as well as the following: (Bökönyi 1977; Hesse 1978; Churcher 1994; Horwitz 2003; Açıkkol 2006; Kersten 2020). Data from Aşıklı Höyük (level 2) are used with permission of Hijlke Buitenhuis, Joris Peters and Nadja Pöllath.

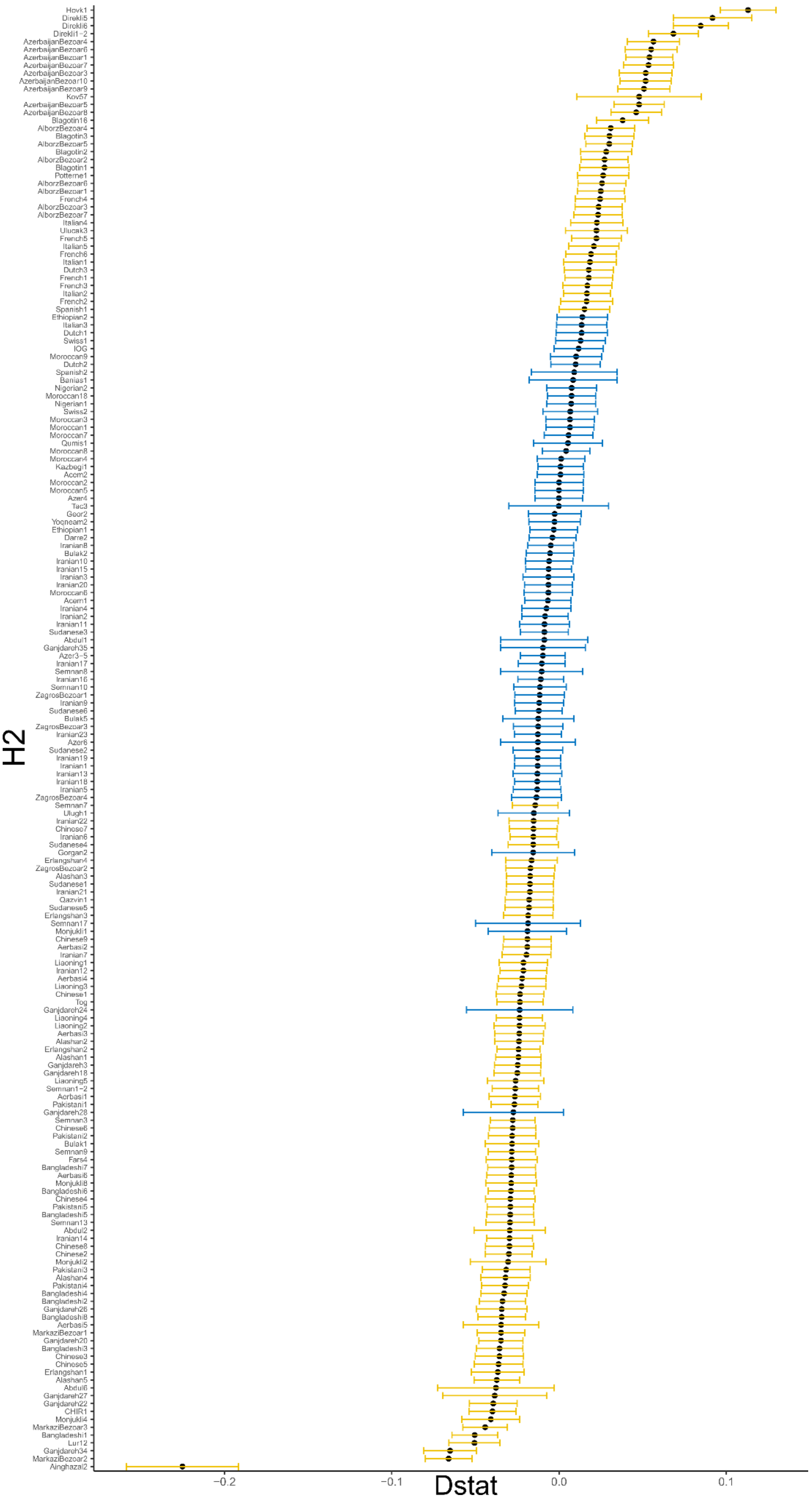


**Figure S11:** *D* statistic of the form *D*(Abdul4, Test; Direkli4, Sheep). Caption on next page.

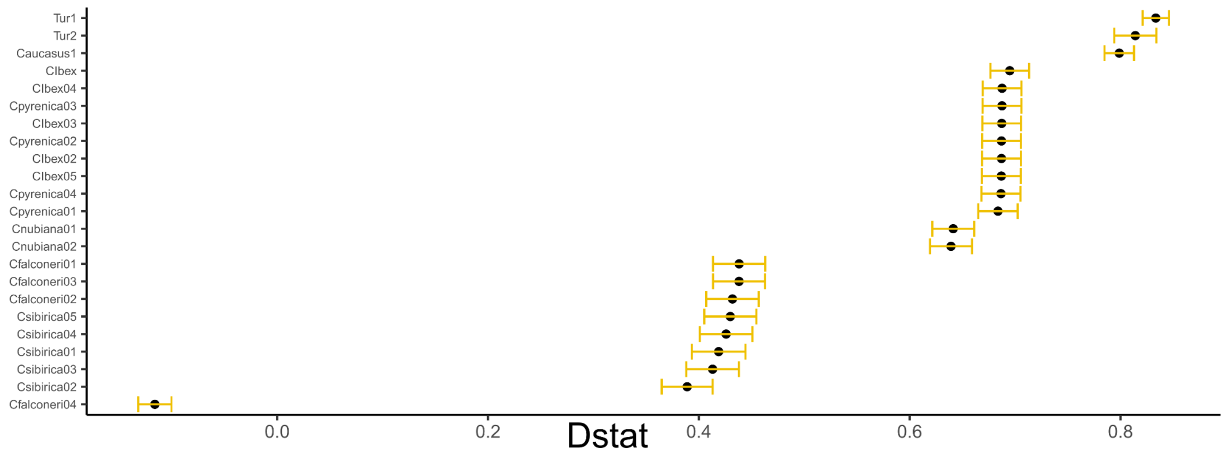


**Figure S11, continued:** *D* statistic of the form *D*(Abdul4, Test; Direkli4, Sheep), where Abdul4 is a *Capra aegagrus* from the 11th millennium BCE Zagros Mountains and Direkli4 is a *Capra caucasica* like genome from the Taurus Mountains. Significant (|Z| ≥ 3) results are indicated in yellow; non-significant (|Z| < 3) in blue. Positive *D* values indicate greater derived allele sharing between Direkli4 and the current H2 genome than between Direkli4 and Abdul4; negative indicates greater allele sharing between Direkli4 and Abdul4. These tests indicate that a Late Pleistocene *C.aegagrus* from Armenia (Hovk-1), from Direkli Cave, and modern bezoar from northwest Iran have elevated Direkli4 derived allele sharing, as well as domestic *Capra hircus* from west Eurasia and Europe.

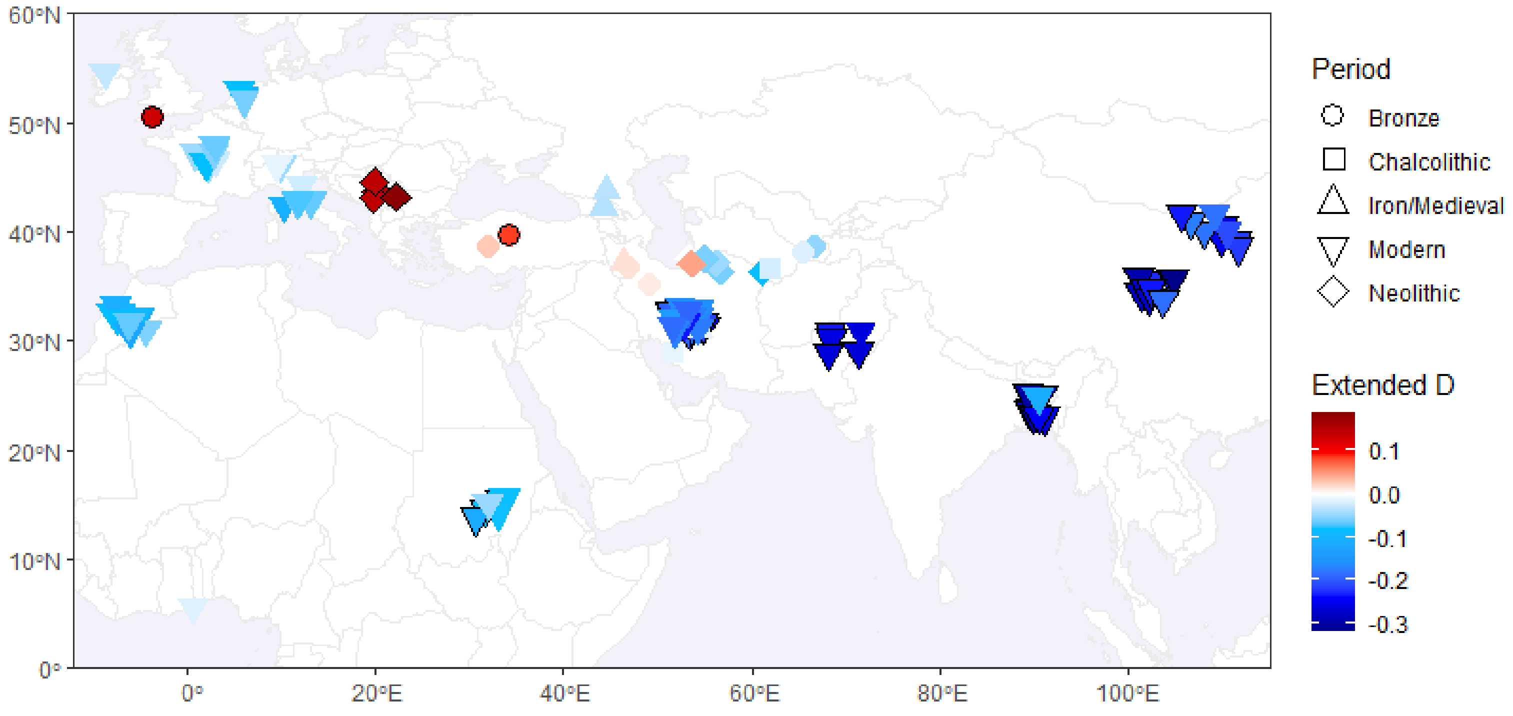


**Figure S12:** Extended *D* statistic in the form {Aceramic Neolithic Zagros, X/H2, Direkli4 (H3), Direkli Wild+Other Wild+*Capra*, Sheep}, using both transitions and transversions. Fill colour indicates extended D value measuring excess of Direkli4-specific alleles in H2, relative to Aceramic Neolithic Zagros goat. "+" demarcates different populations in which the ancestral allele (relative to the derived Direkli4 allele) must be observed and fixed. Black borders indicate a Z score ≥ |3|.

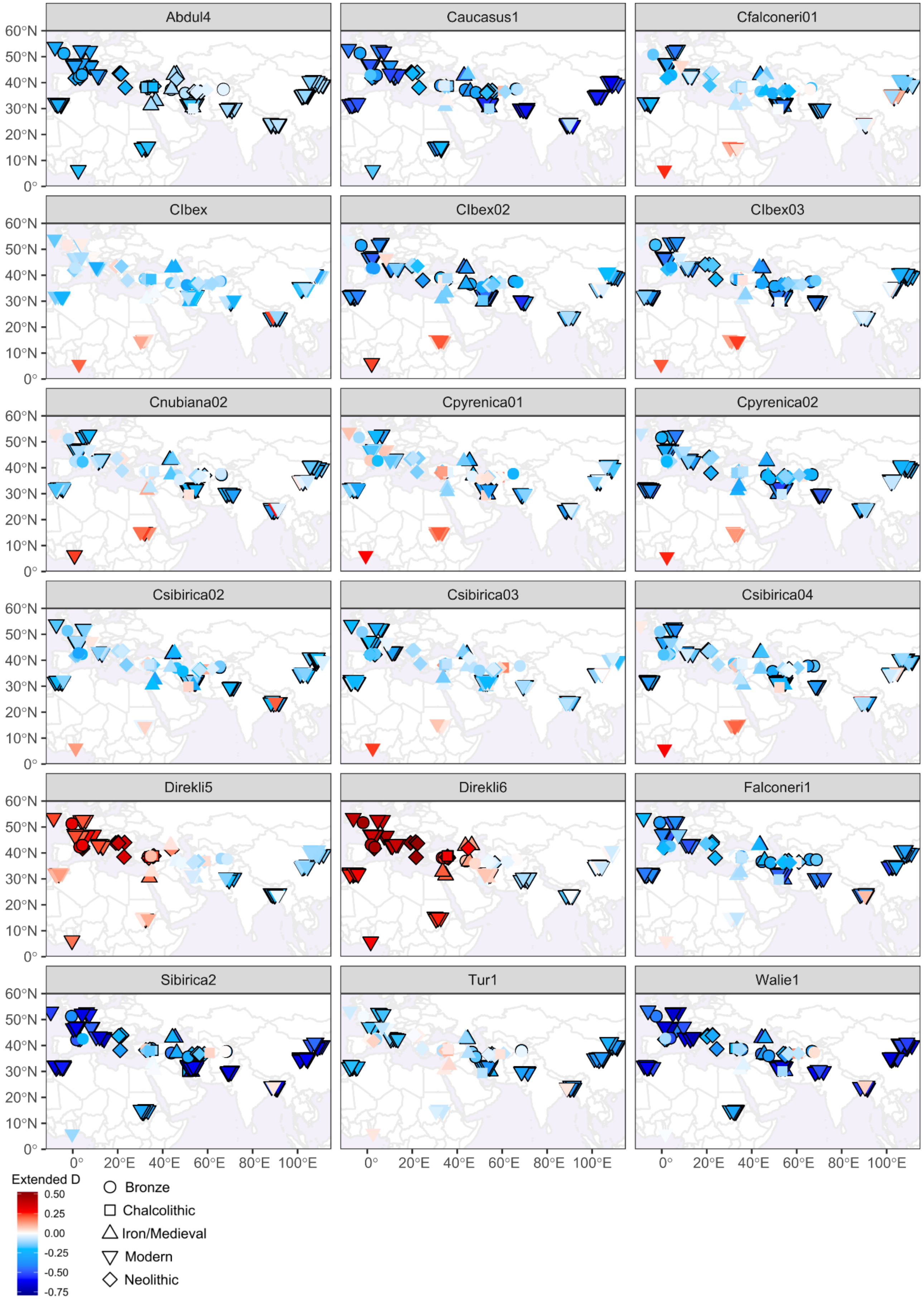


**Figure S13:** Extended *D*, rotating H3 through *Capra* genus genomes.

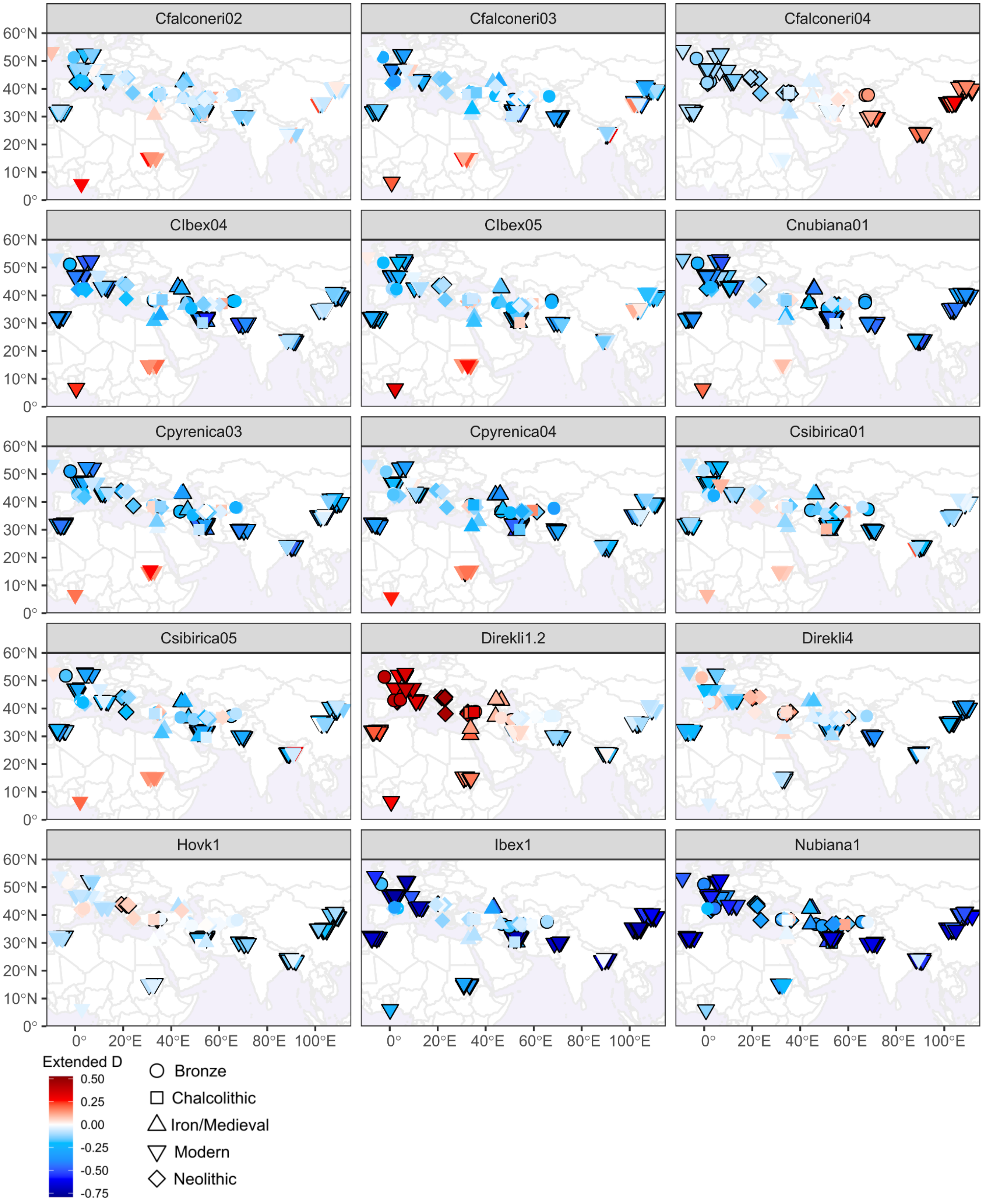


**Figure S13, continued:** Extended *D*, rotating H3 through *Capra* genus genomes.

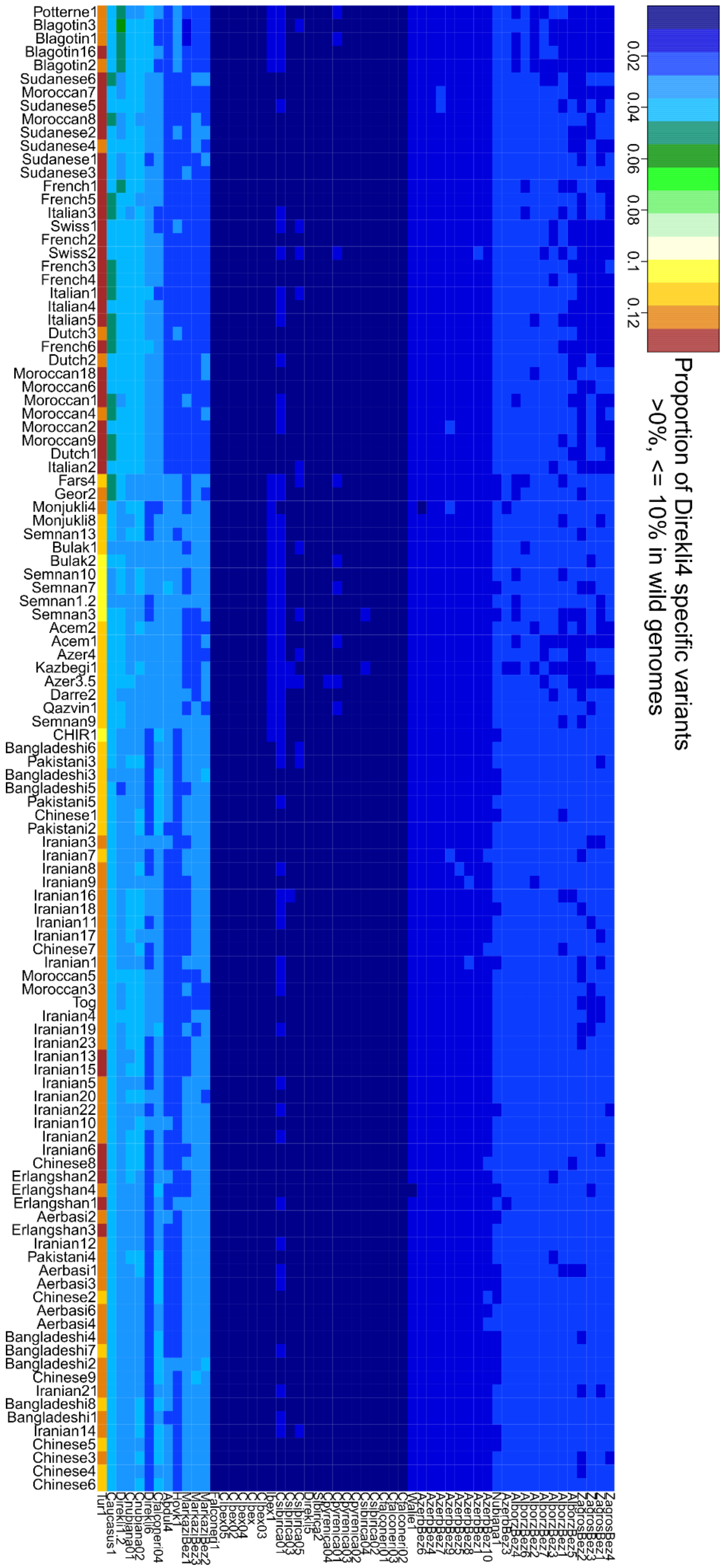


**Figure S14:** Figure caption provided on next page.

**Figure S14:** Distribution of Direkli4 specific variants also present at >0%, ≤10% in wild genomes, expressed as a proportion of the total number of these variants in the tested domestic genomes. The majority of "shared Direkli4" variants are observed with Tur1, particularly for modern European and African goats. Modern European and African goats also show "Direkli4" allele sharing with the historic-era *Capra cylindricornis* Caucasus1, while ancient European goats show allele sharing with ancient Turkish wild Direkli1-2.

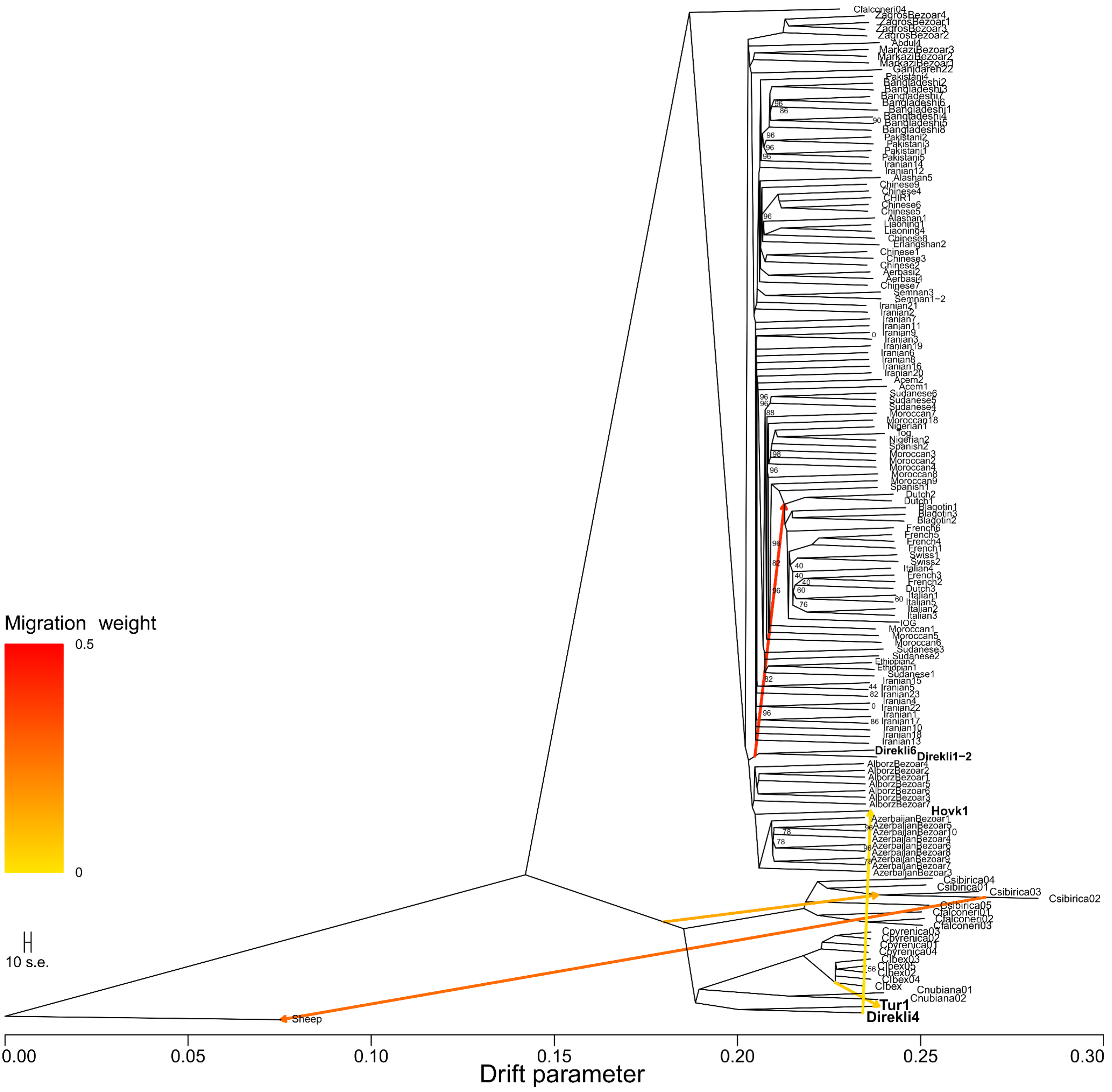


**Figure S15:** Treemix results for *m*=5, using 223,772 transversion variants ascertained on sheep, k=1000. Support values for nodes are indicated when <100 (50 bootstrap replicates).

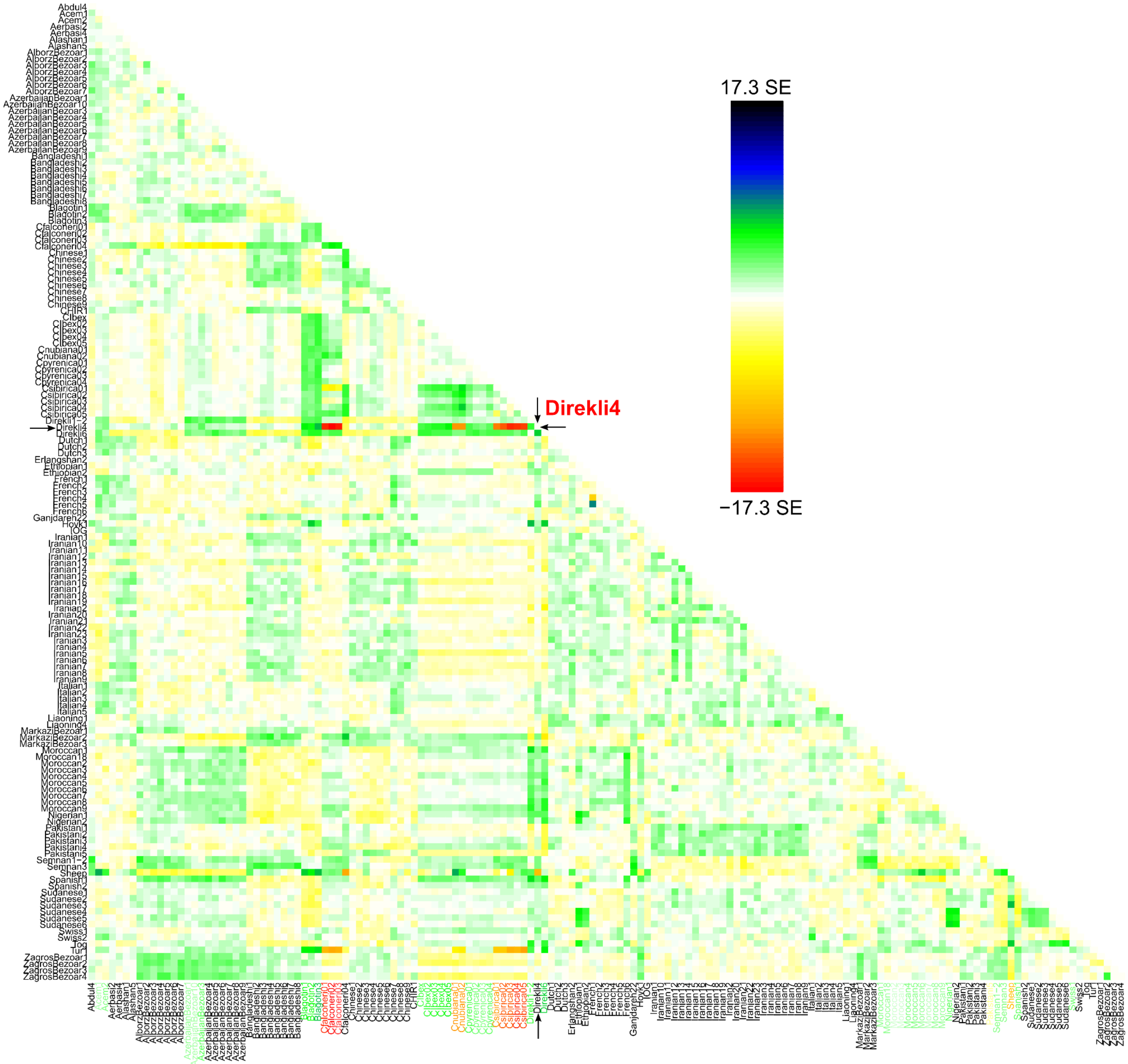


**Figure S16:** Residuals for Treemix m=5. Some genomes have unmodeled affinity with Direkli4: Neolithic Serbian genomes from Blagotin; wild Late Pleistocene bezoar (*Capra aegagrus*) from Direkli Cave; Alpine Ibex (*Capra ibex*); Pyrenean ibex (*Capra pyrenaica*); Moroccan goat; and the outgroup Sheep. Populations which have lower genetic affinity with Direkli4 than the model predicts: the markhor (*Capra falconeri*); Siberian ibex (*Capra sibirica*); and Nubian ibex (*Capra nubiana*). X-axis labels are coloured by their residual with Direkli4.

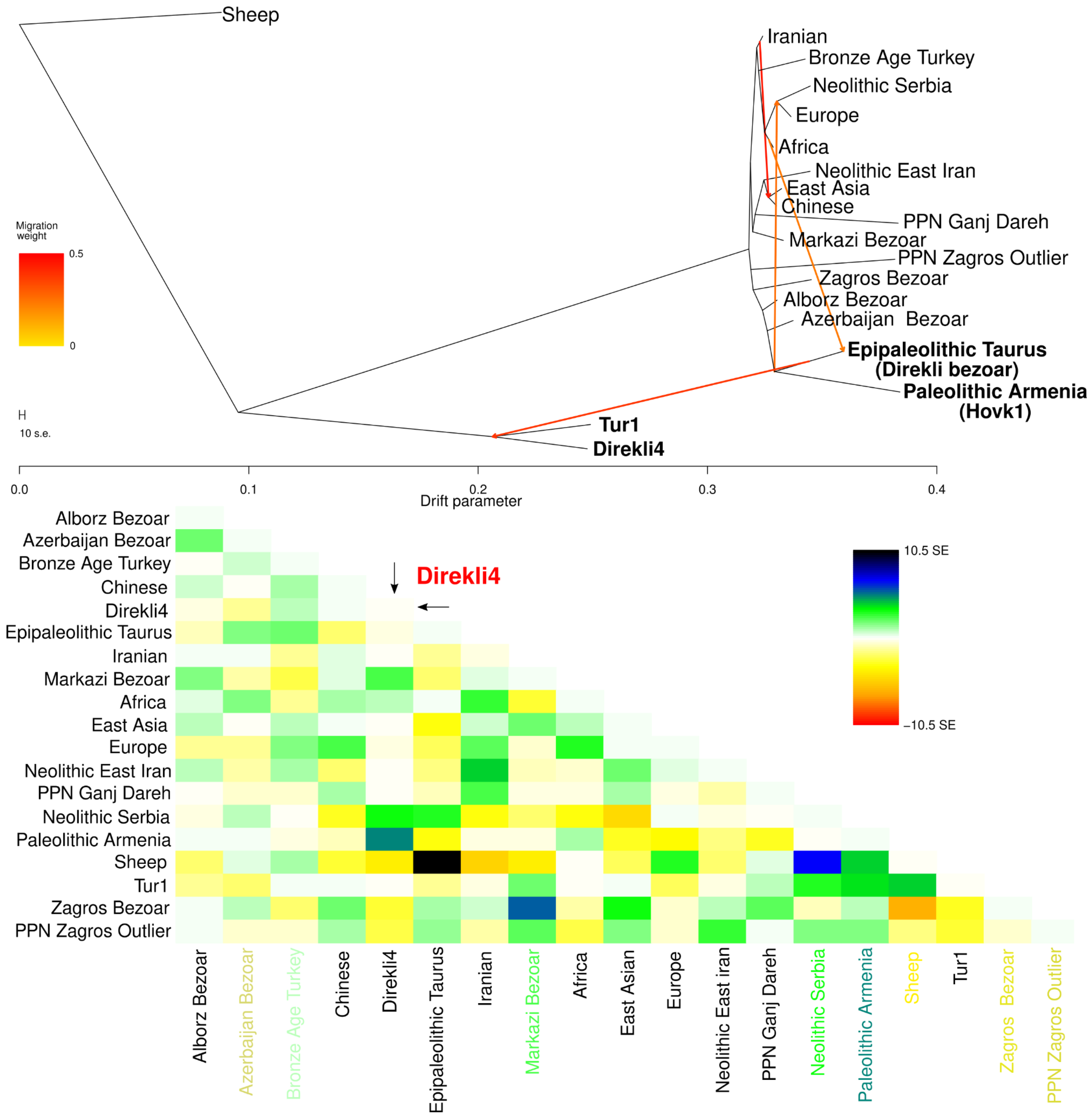


**Figure S17:** Orientagraph model and residuals for m=4, using 104,550 transversion variants and k=1000. Residual x-axis labels are coloured by their residual with Direkli4.

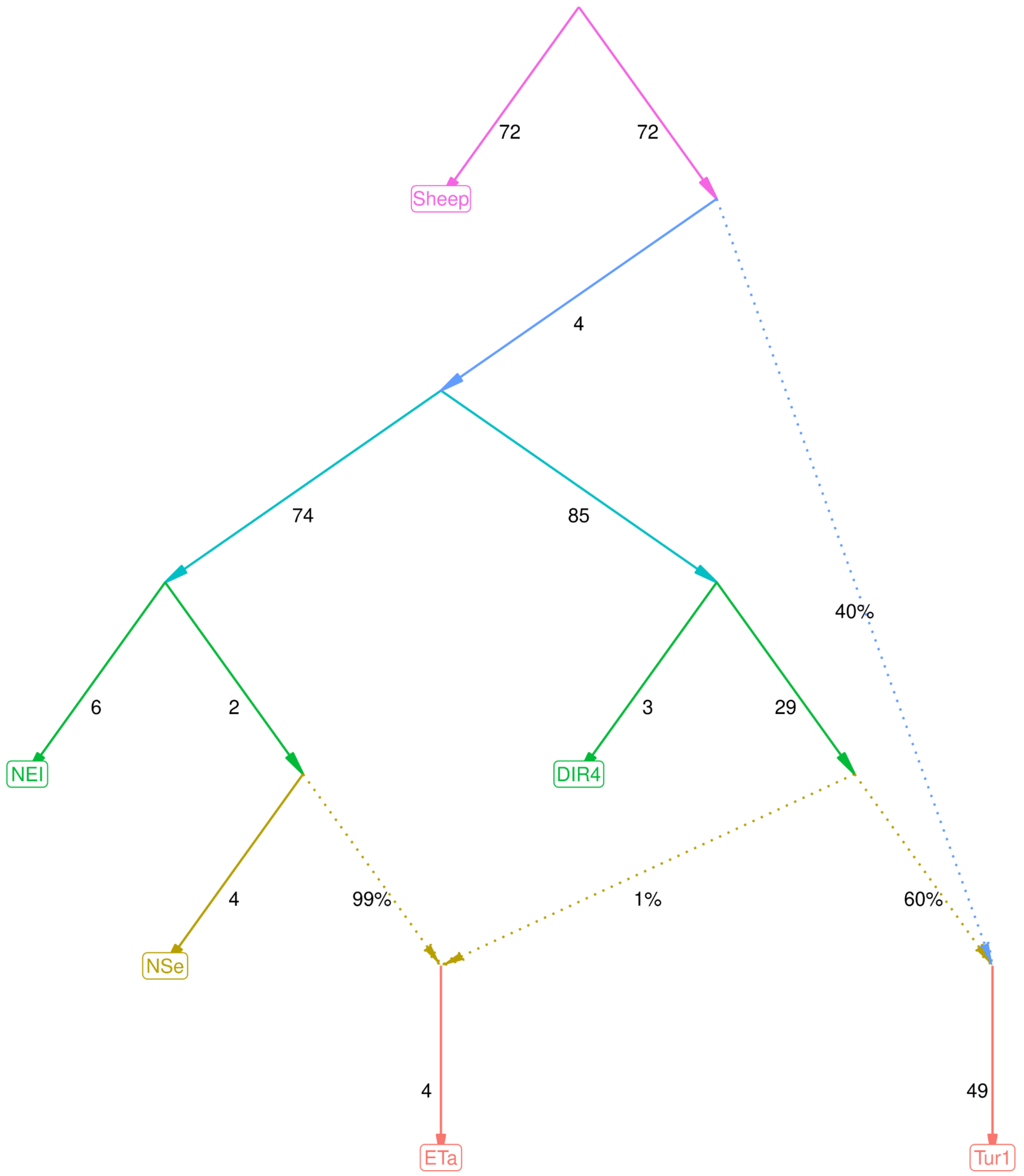


**Figure S18:** ADMIXTOOLS2 (Maier et al. 2022) automated graph exploration for m=2 (two admixture events), using 223,772 biallelic variants ascertained in sheep and presenting the best fitting graph as measured by log likelihood score (-0.5). Graphs which fit the data “as good as” the above graph are available at https://osf.io/3ecqd/, along with the best fitting graph for m=3. Abbreviations used for populations defined in Table S3: ETa=Direkli Cave bezoar (Epipaleolithic Taurus), DIR4=Direkli4, NSe=Neolithic Serbia, NEI=Neolithic East Iran, ur1=Tur1. Sheep was used as the outgroup.

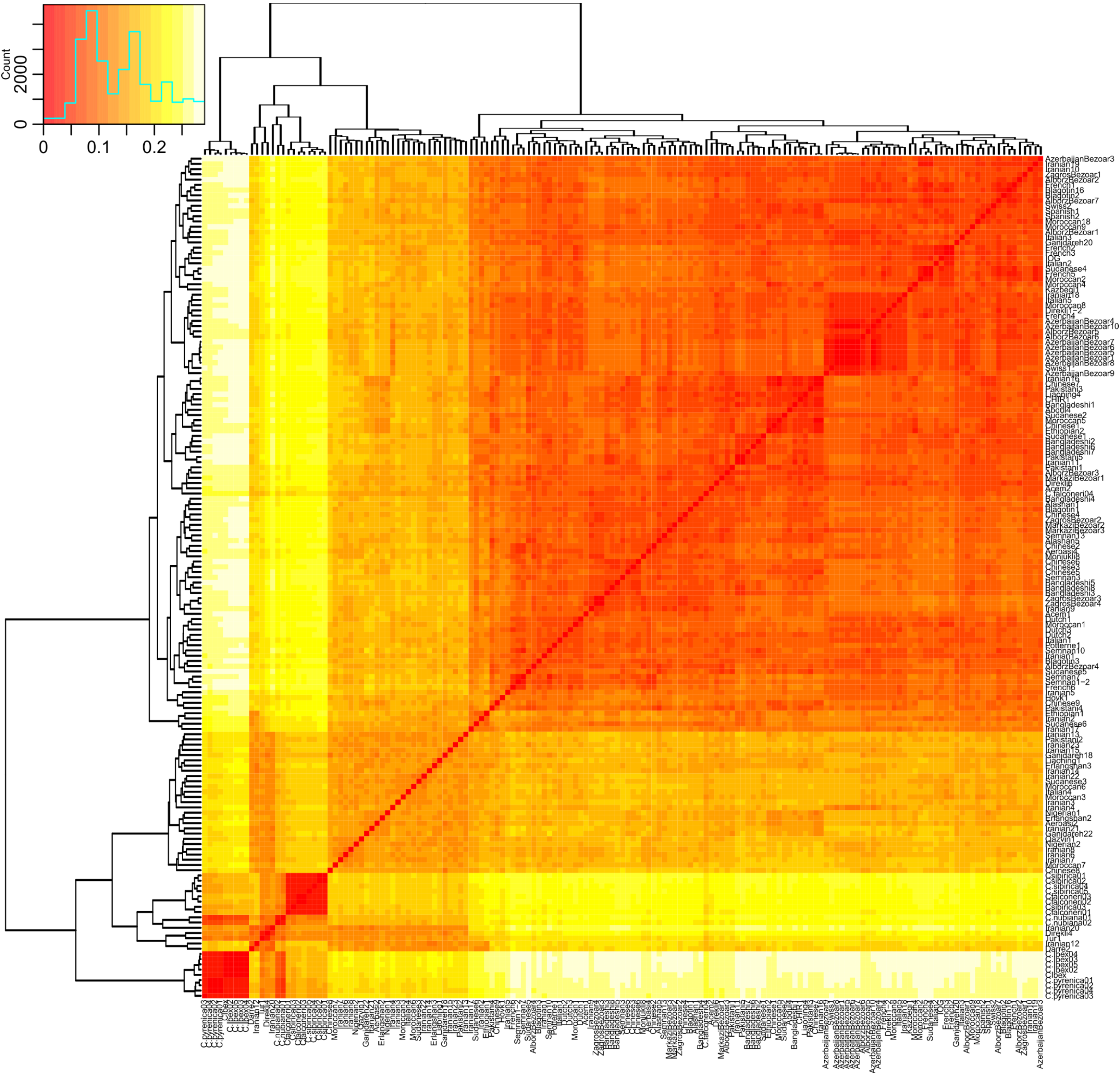


**Figure S19: Clustered heatmap of IBS data for ARS1 1:104,150,173-104,349,720.** Sheep and lower coverage Nubiana1, Caucasus1 are removed.

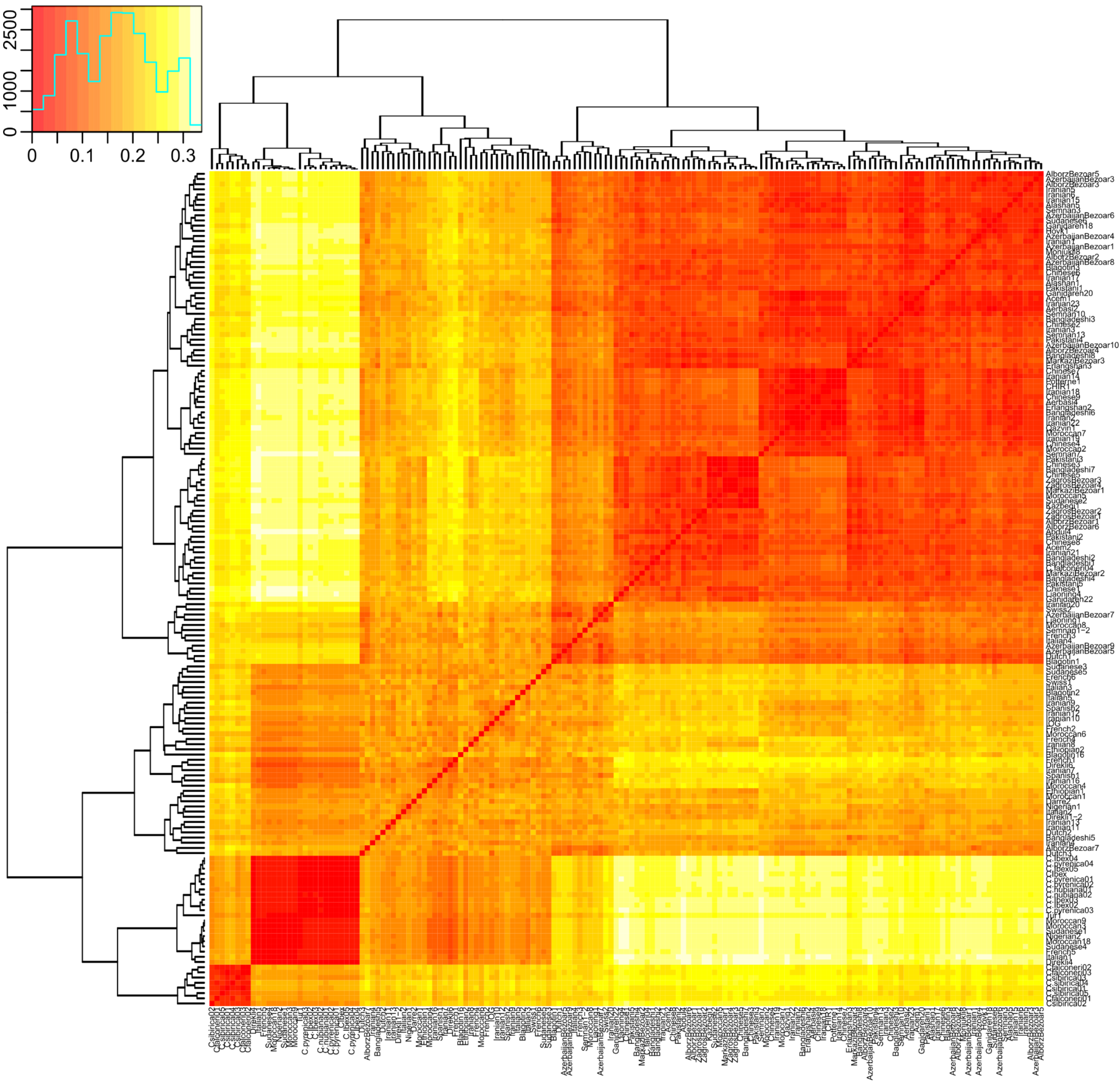


**Figure S20: Clustered heatmap of IBS data for ARS1 2:24,324,410-24,369,675.** Sheep and lower coverage Nubiana1, Caucasus1 are removed.

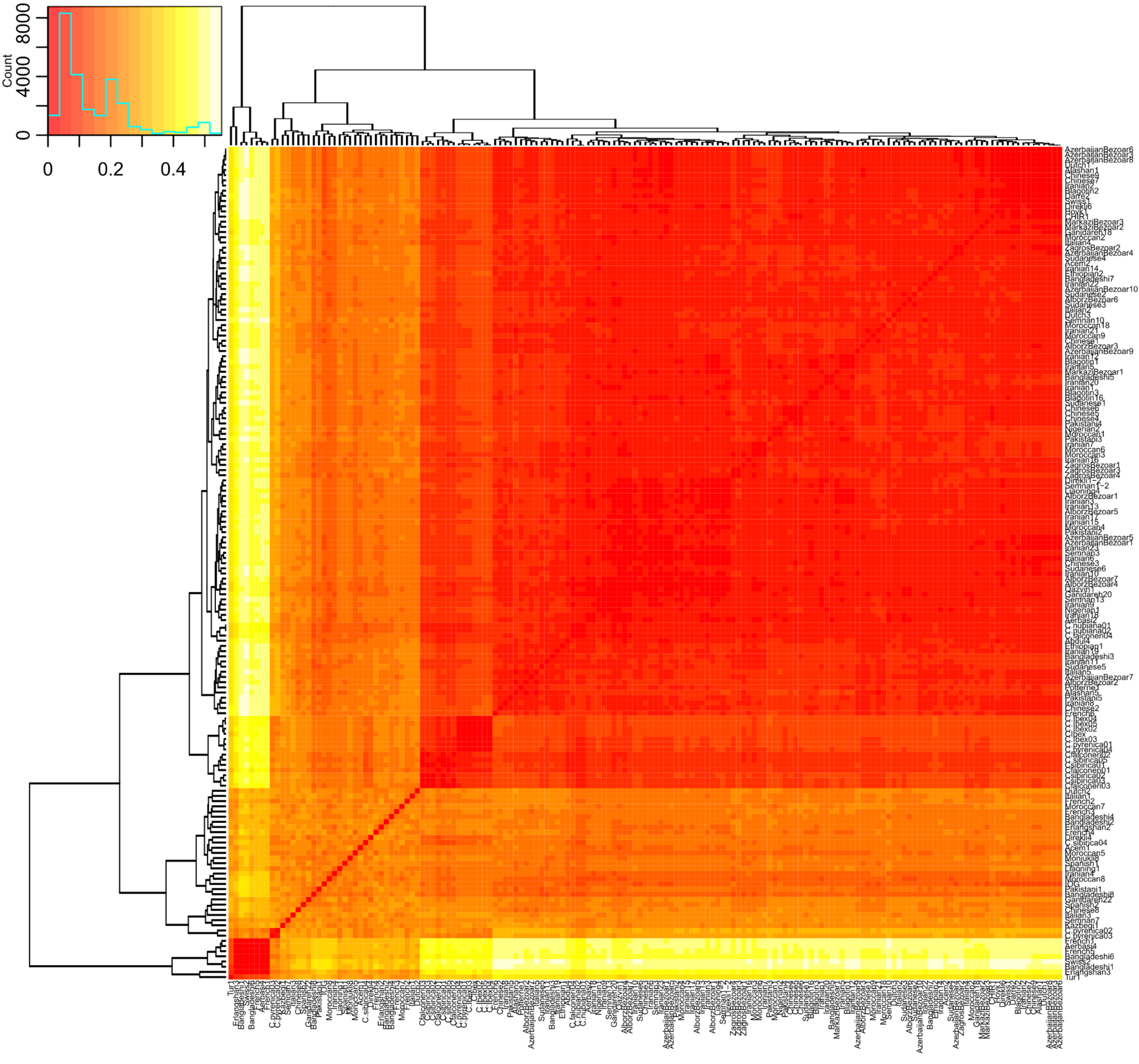


**Figure S21: Clustered heatmap of IBS data for ARS1 13:66,710,508-66,749,824.** Sheep and lower coverage Nubiana1, Caucasus1 are removed.

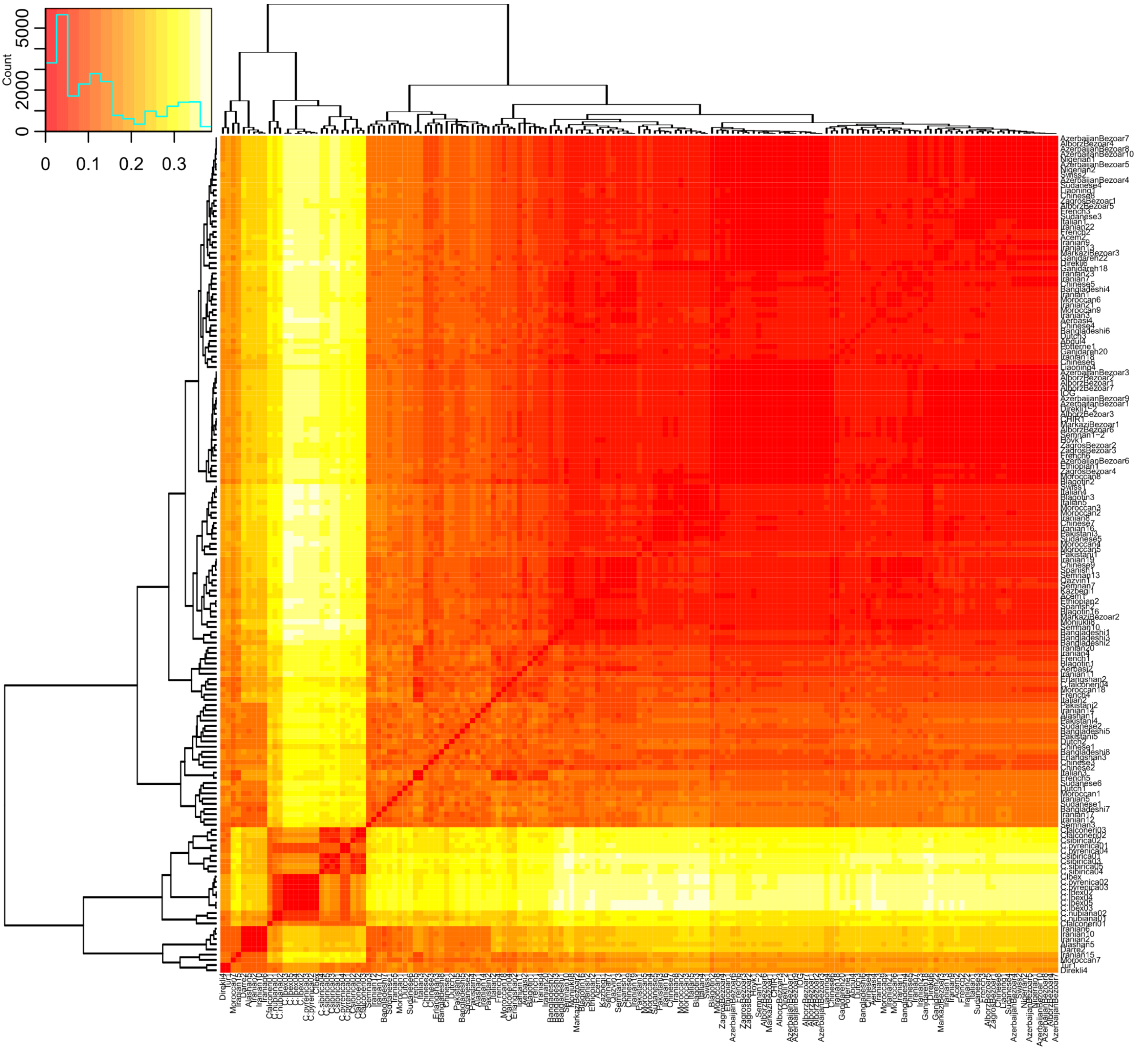


**Figure S22: Clustered heatmap of IBS data for ARS1 1:118,433,695-118,469,862.** Sheep and lower coverage Nubiana1, Caucasus1 are removed.

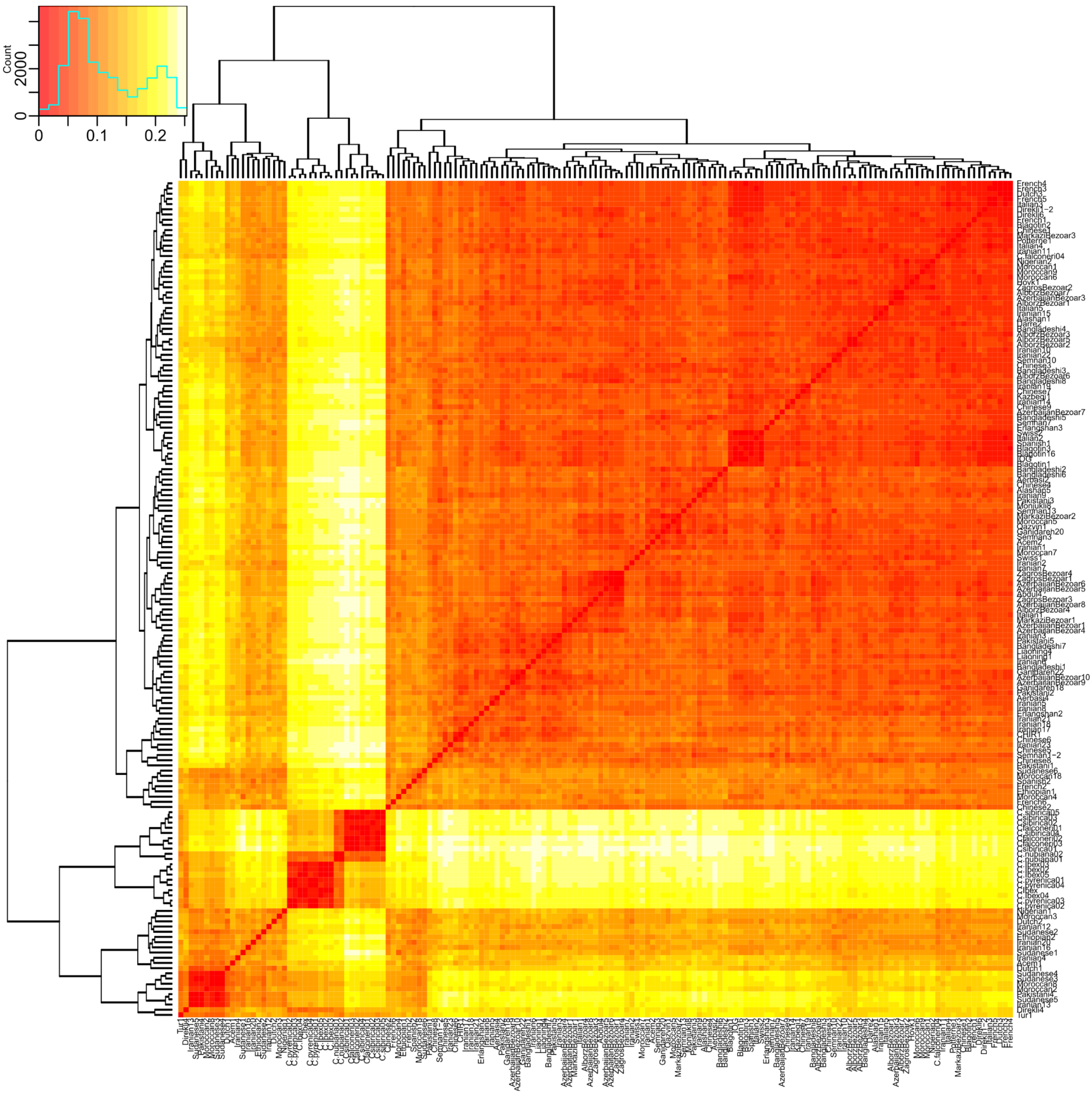


**Figure S23: Clustered heatmap of IBS data for ARS1 6:20,670,803-20,728,594.** Sheep and lower coverage Nubiana1, Caucasus1 are removed.

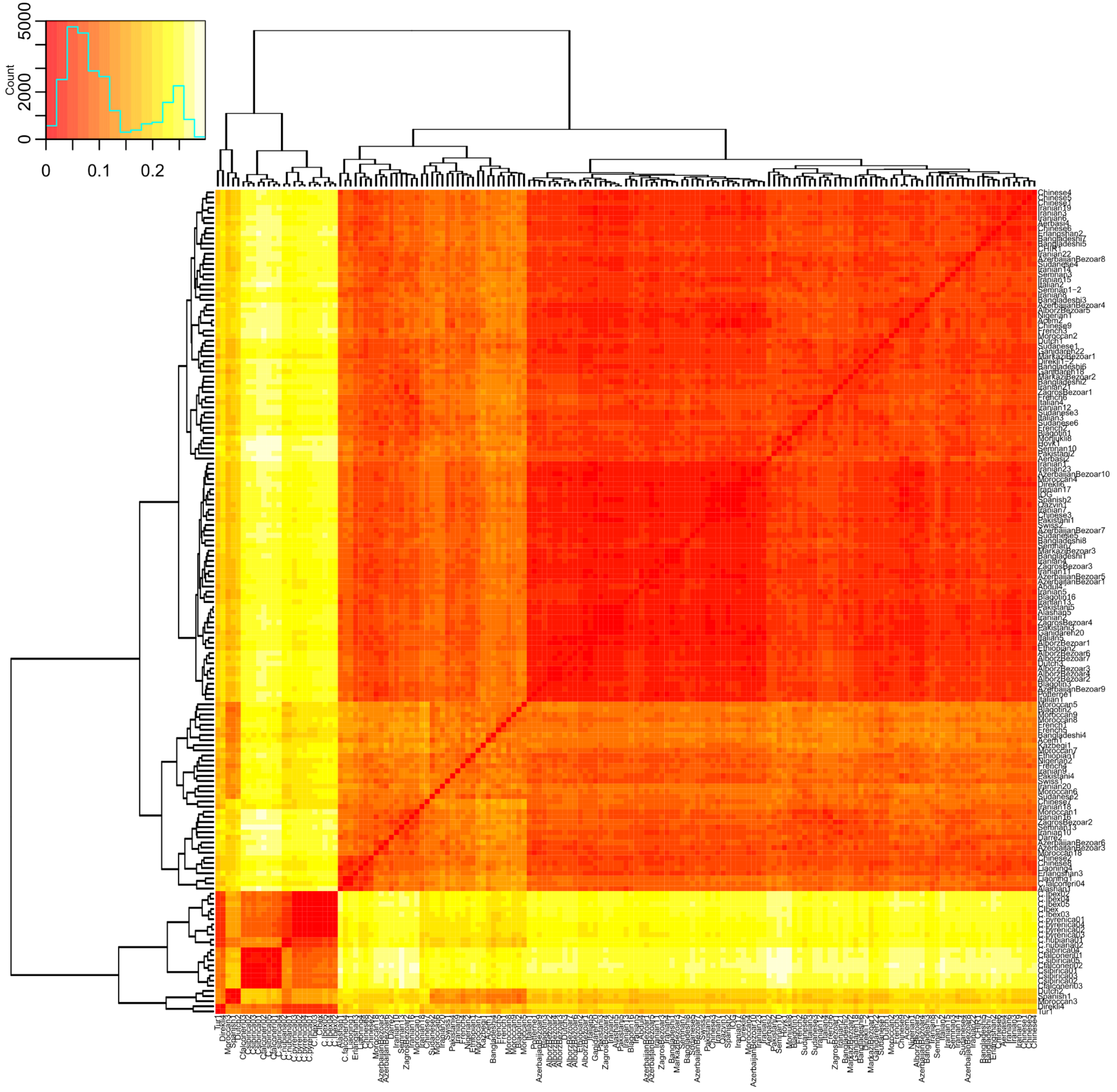


**Figure S24: Clustered heatmap of IBS data for ARS1 6-75,470,180-75,509,529.** Sheep and lower coverage Nubiana1, Caucasus1 are removed.

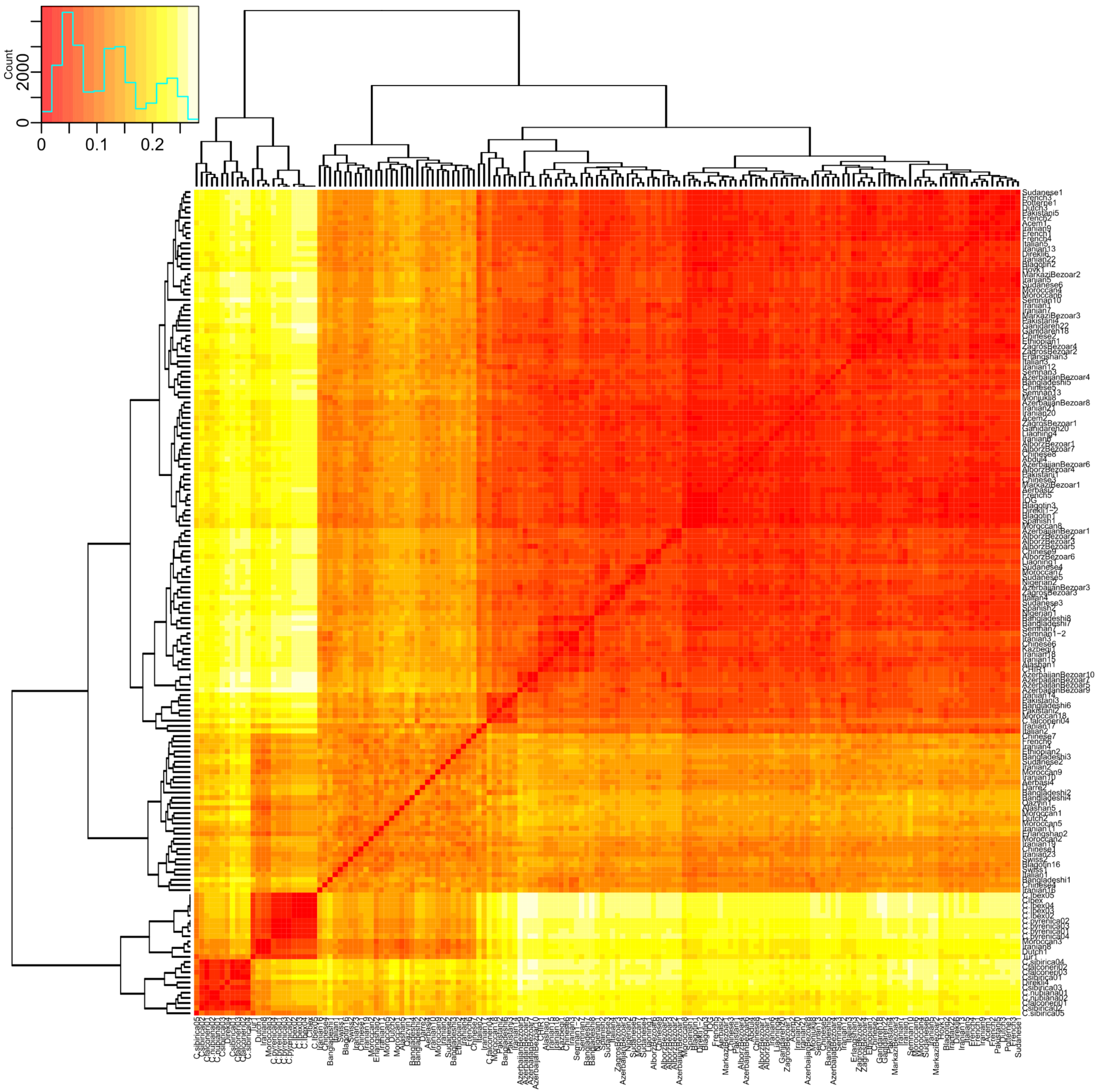

**Figure S25: Clustered heatmap of IBS data for ARS1 7:24,370,068-24,409,932.** Sheep and lower coverage Nubiana1, Caucasus1 are removed.

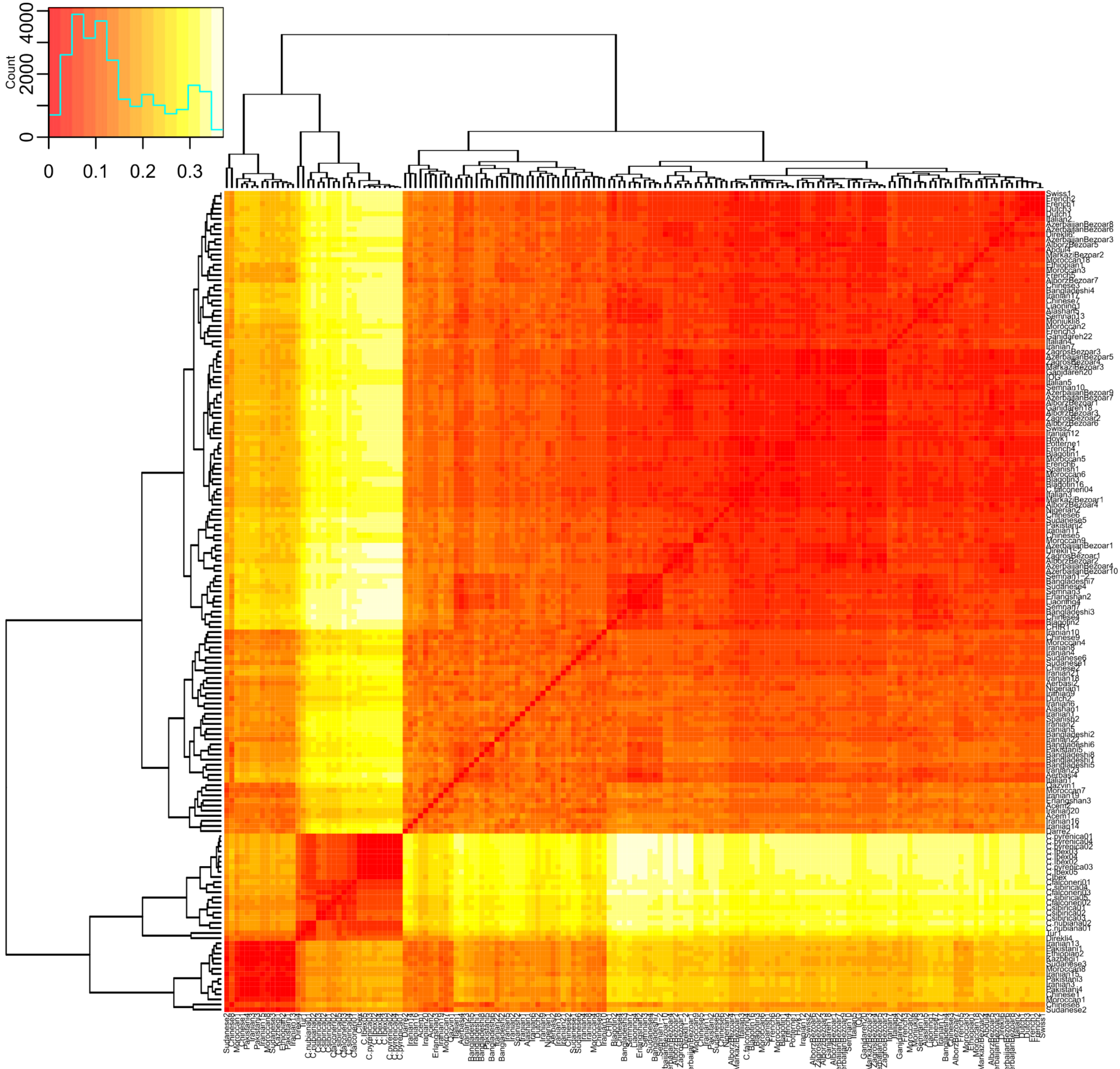


**Figure S26: Clustered heatmap of IBS data for ARS1 11:66,897,539-66,929,689.** Sheep and lower coverage Nubiana1, Caucasus1 are removed.

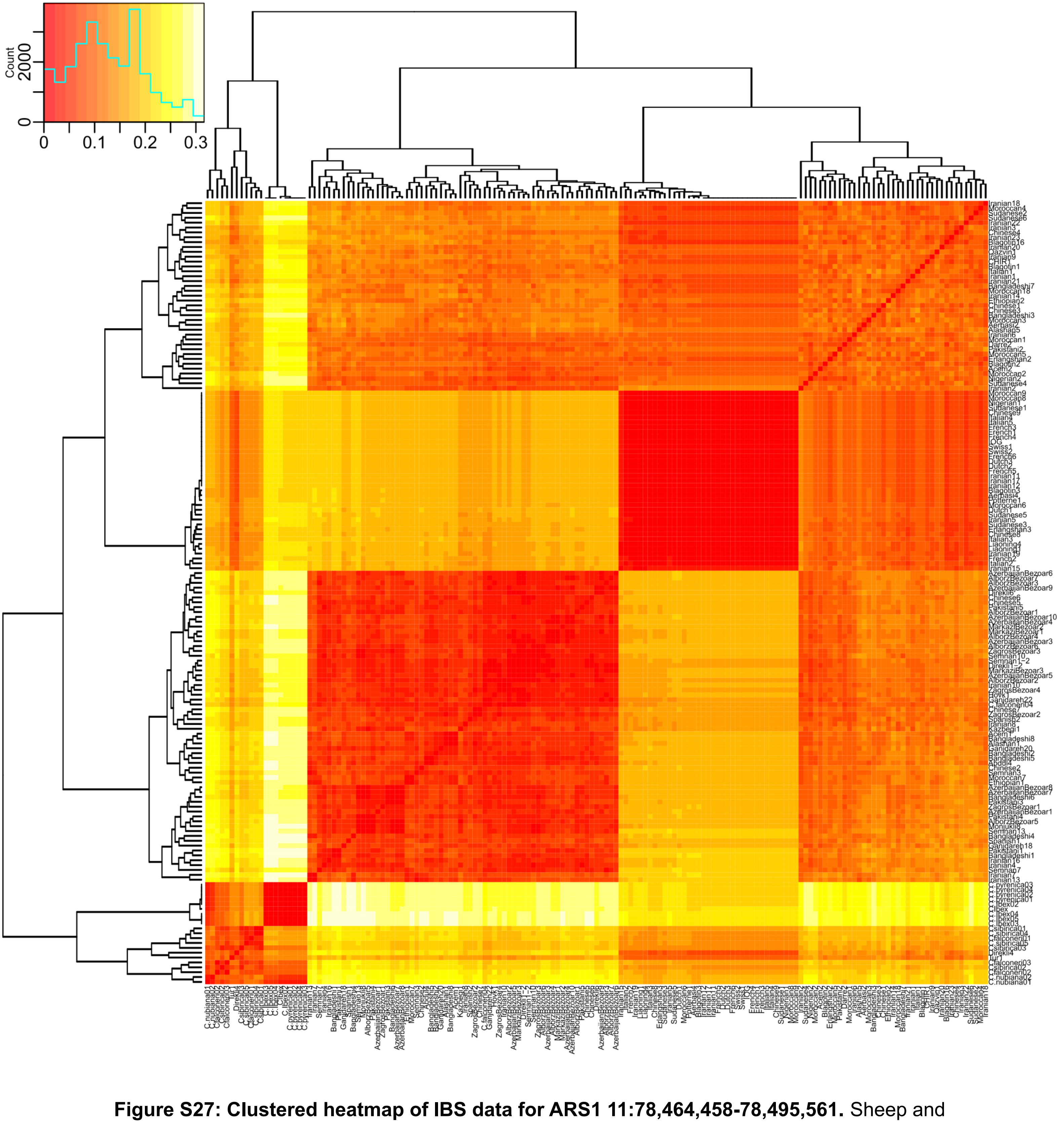


**Figure S27: Clustered heatmap of IBS data for ARS1 11:78,464,458-78,495,561.** Sheep and lower coverage Nubiana1, Caucasus1 are removed.

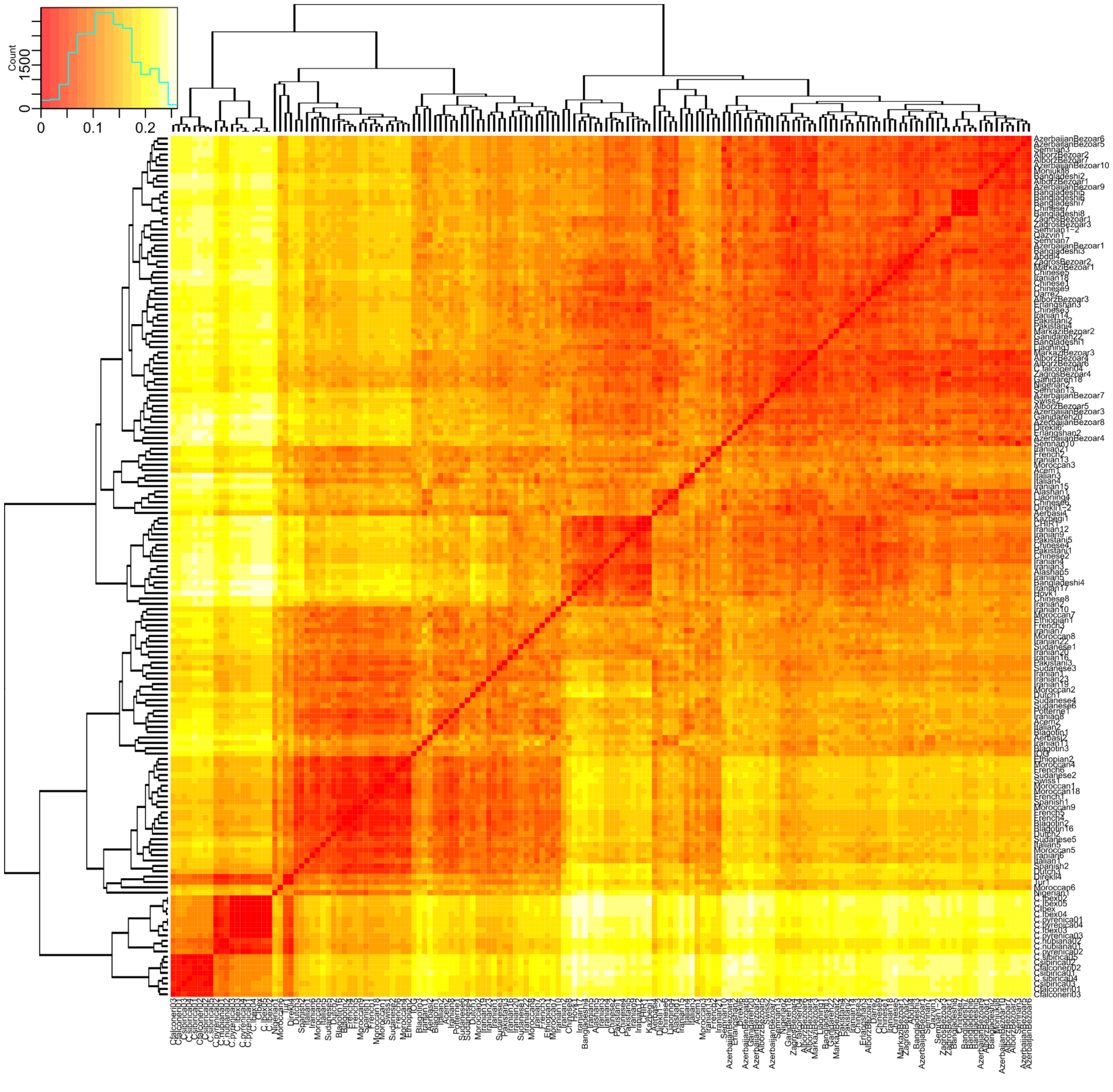


**Figure S28: Clustered heatmap of IBS data for ARS1 19:13,575,349-13,629,669.** Sheep and lower coverage Nubiana1, Caucasus1 are removed.

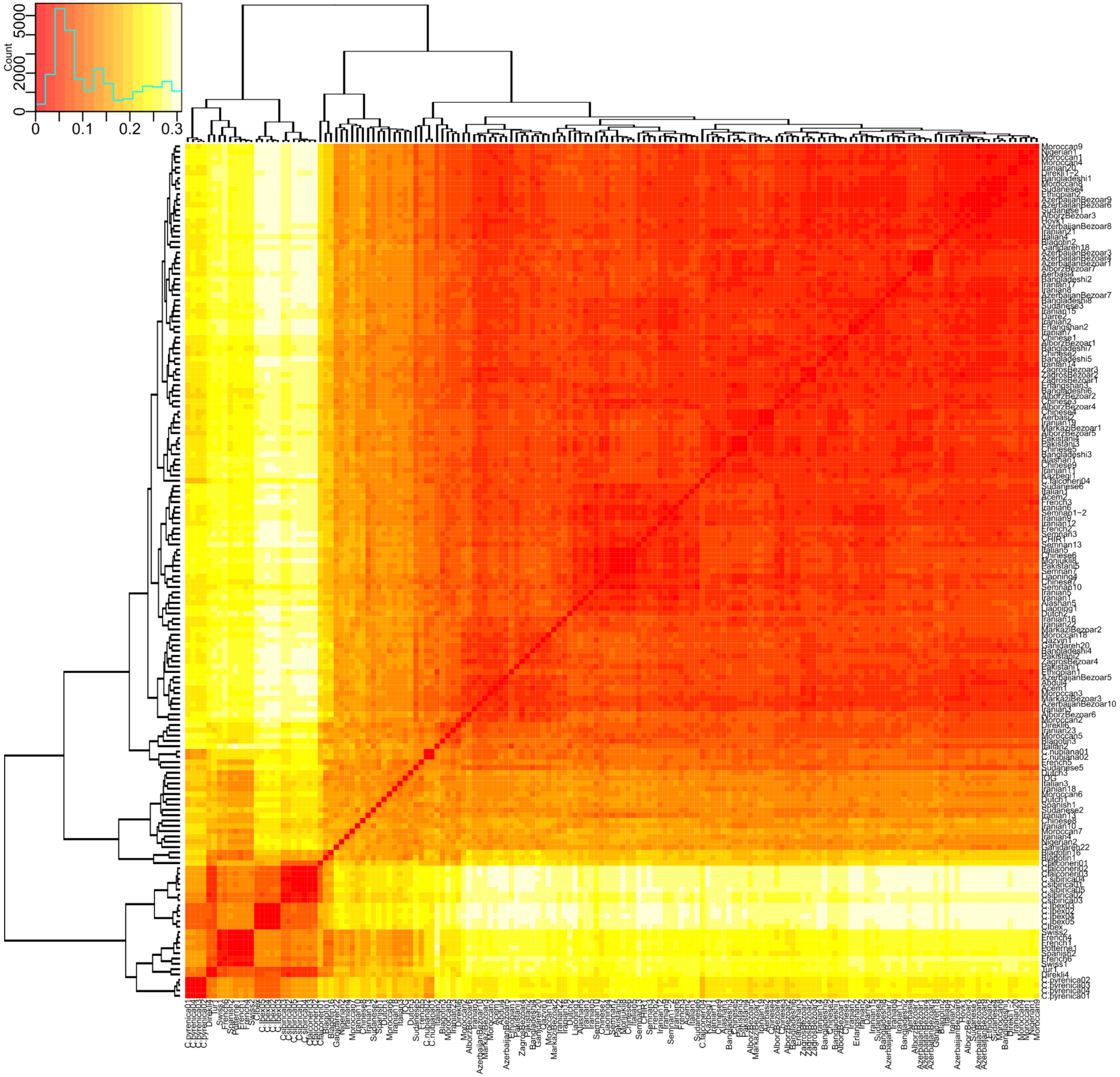


**Figure S29: Clustered heatmap of IBS data for ARS1 29:4,557,144-4,627,040.** Sheep and lower coverage Nubiana1, Caucasus1 are removed.

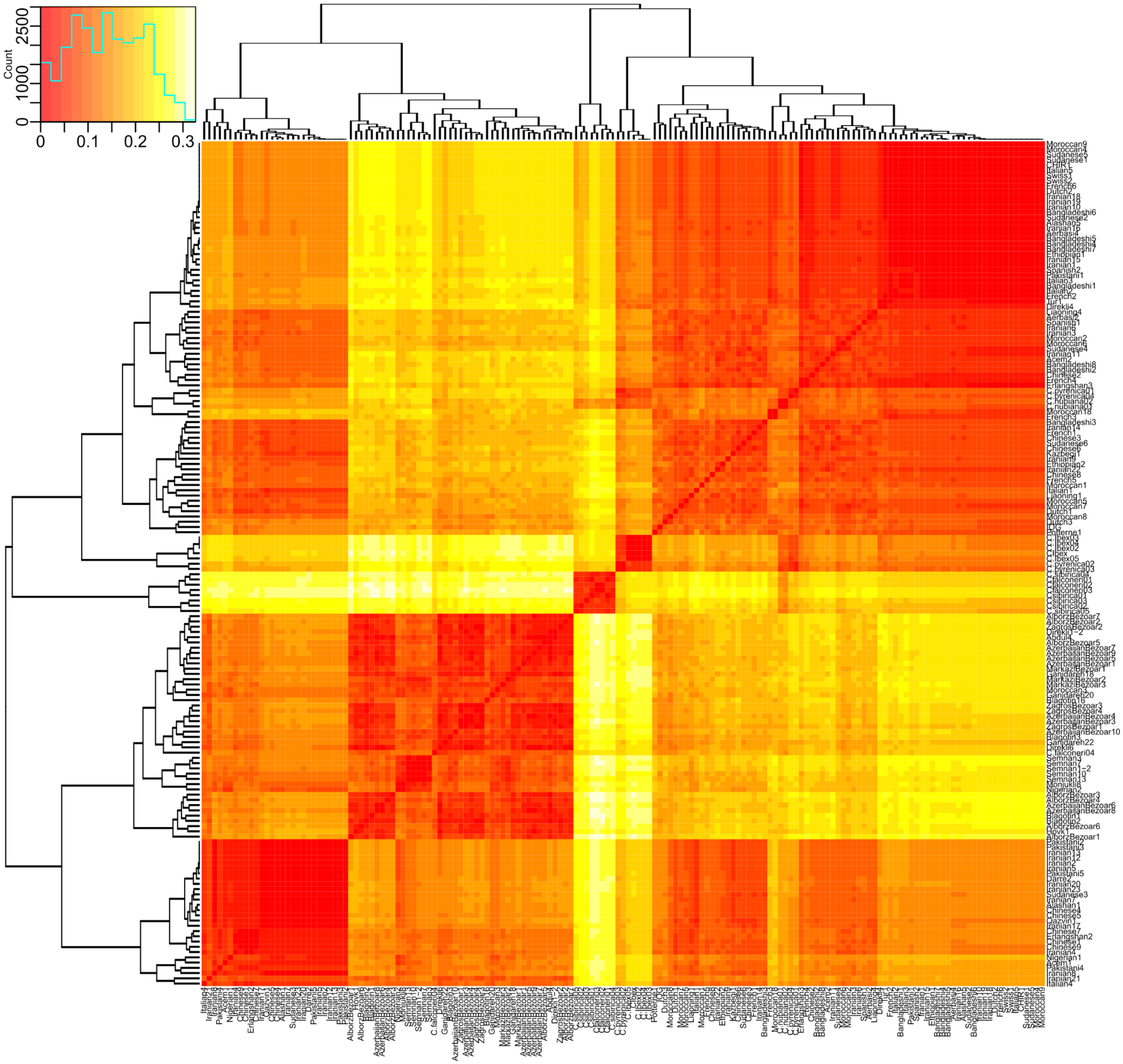


**Figure S30: Clustered heatmap of IBS data for ARS1 29:46,175,547-46,301,705.** Sheep and lower coverage Nubiana1, Caucasus1 are removed.

**Figures S31, S32: NJ trees of IBS data of introgressed regions.** Outgroups (Sheep, Table S3) are not shown. Provided as additional data file 1 and 2.
