## Supplementary figures and images for "A novel lineage of the Capra genus discovered in the Taurus Mountains of Turkey using ancient genomics"

### Supplementary Data File 1

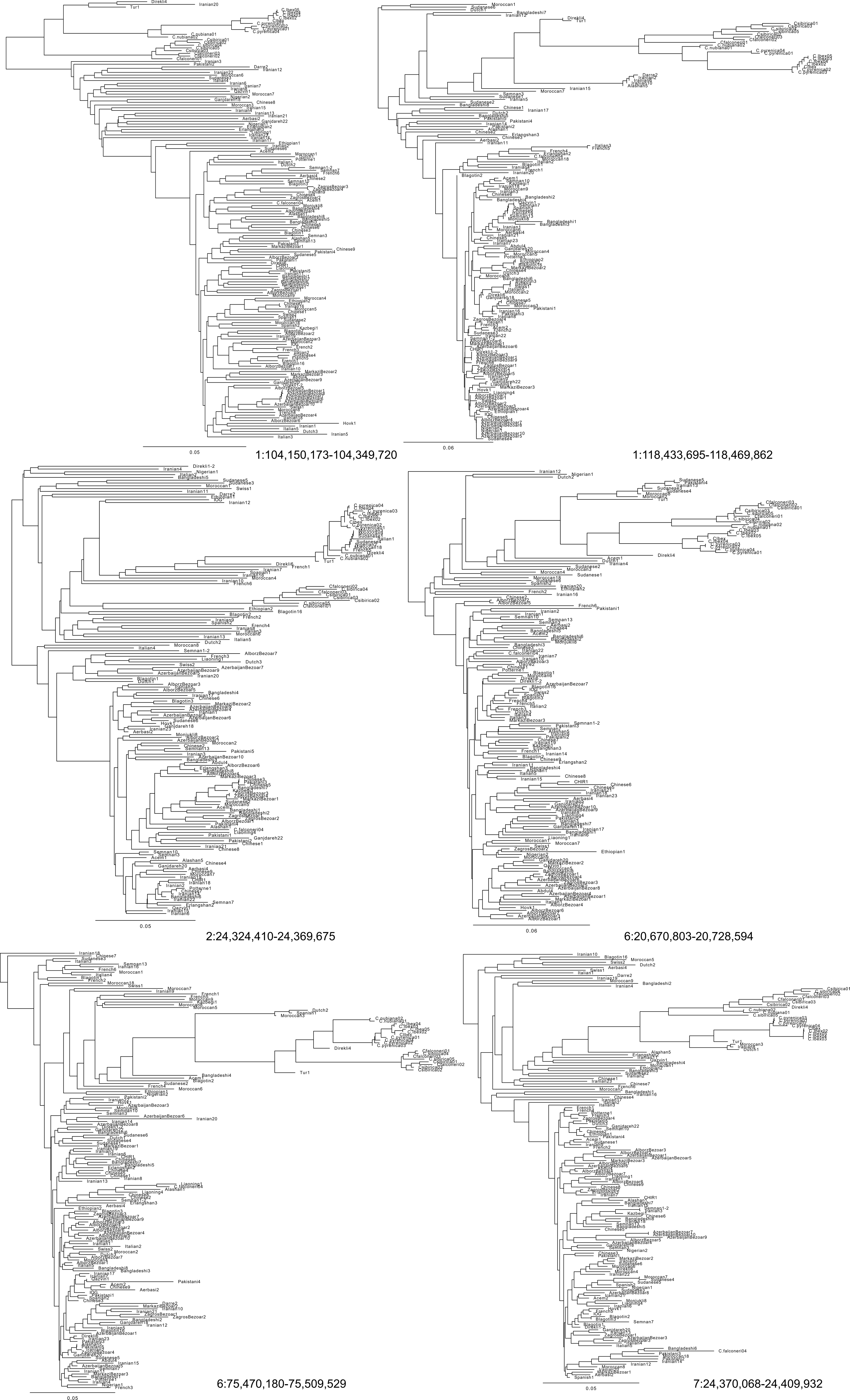

### Supplementary Data File 2

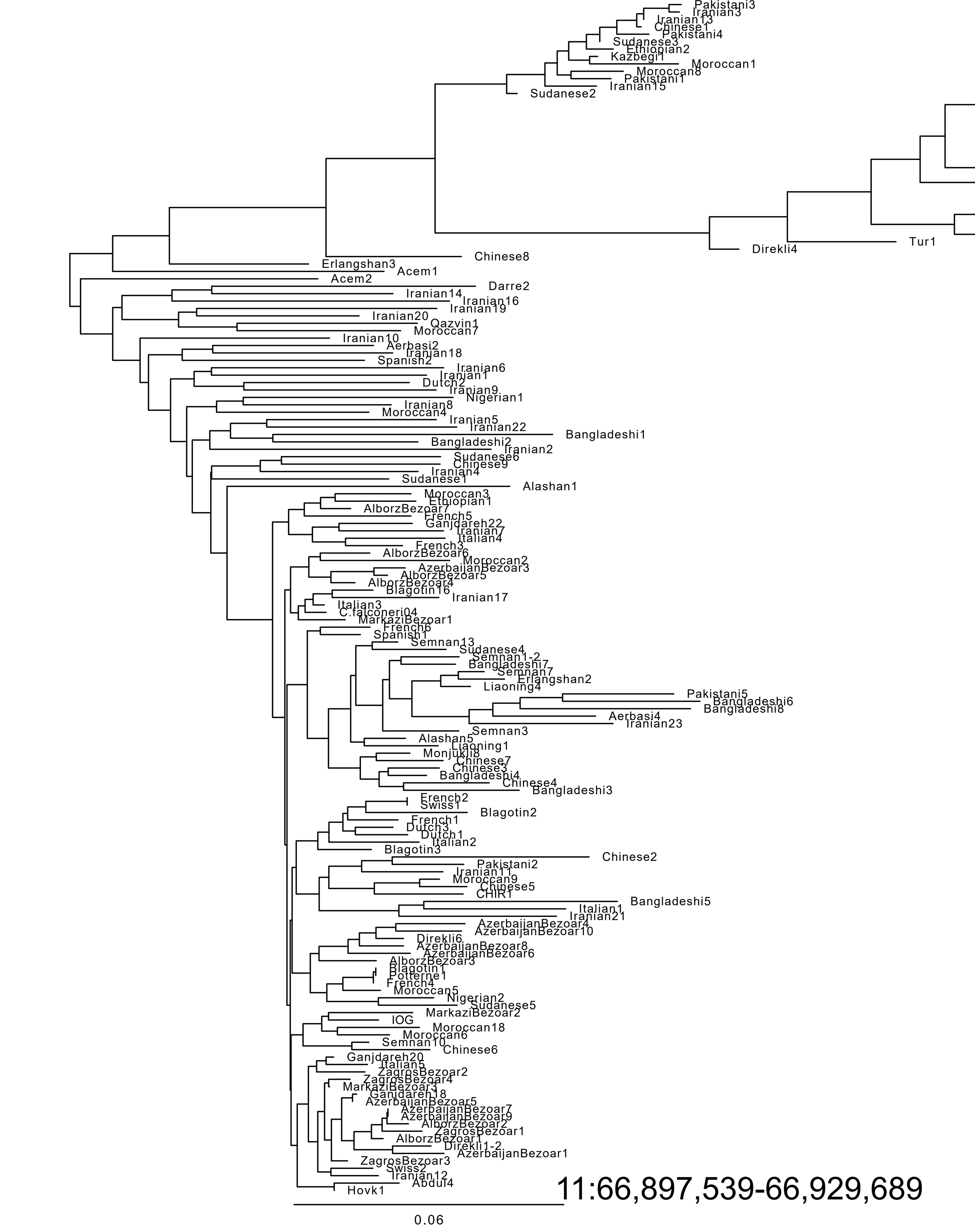

11:66,897,539-66,929,689

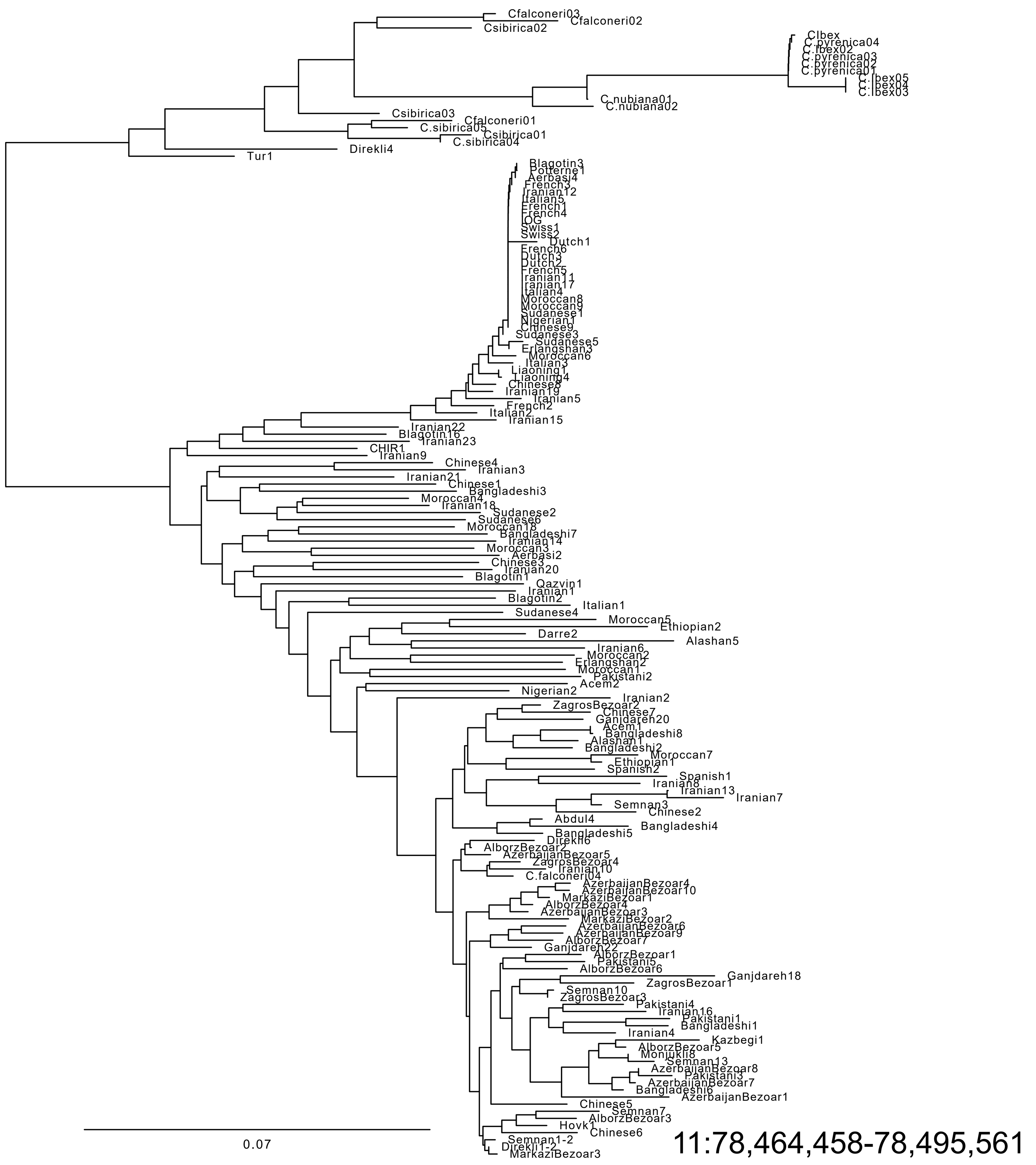

11:78,464,458-78,495,561

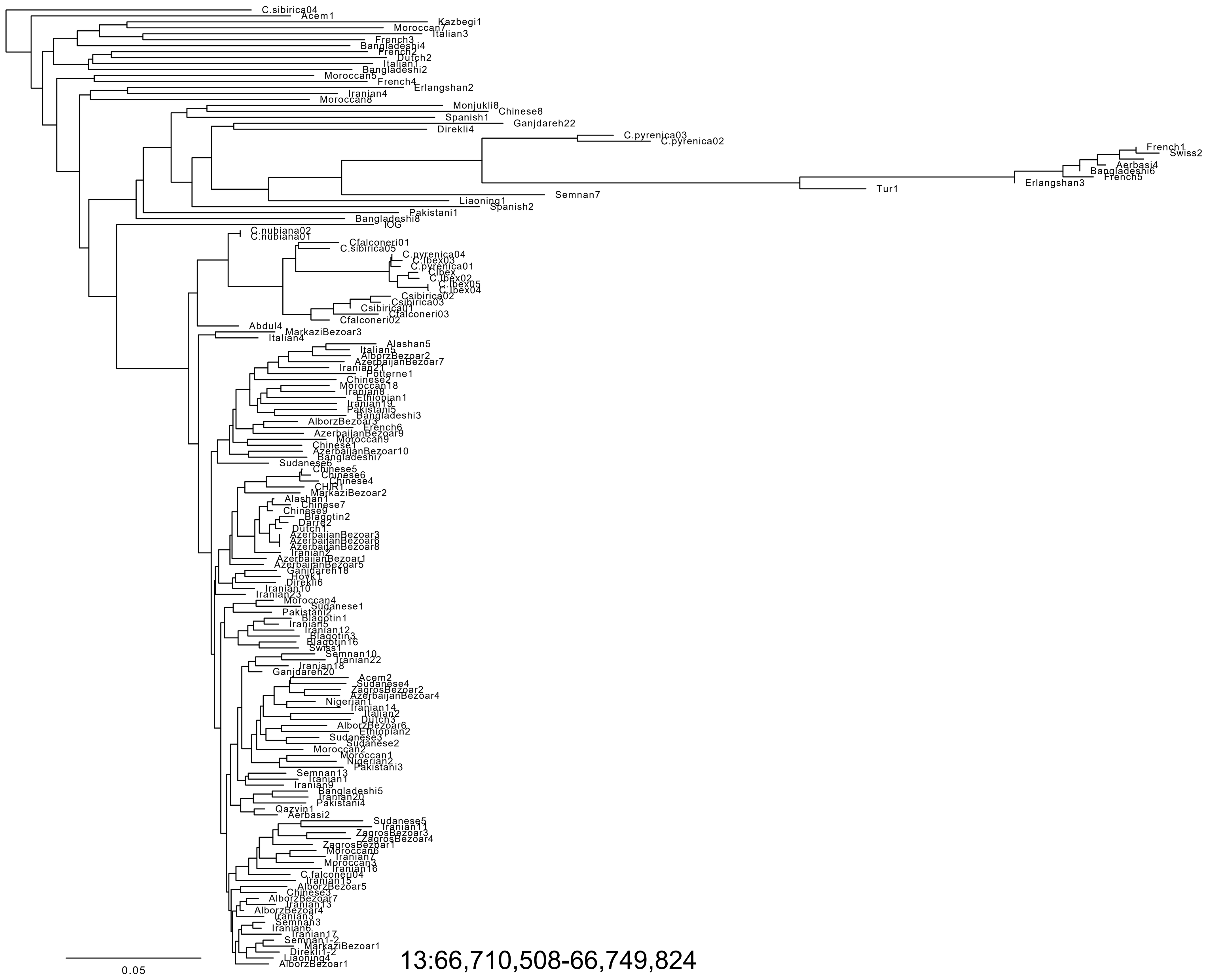

13:66,710,508-66,749,824

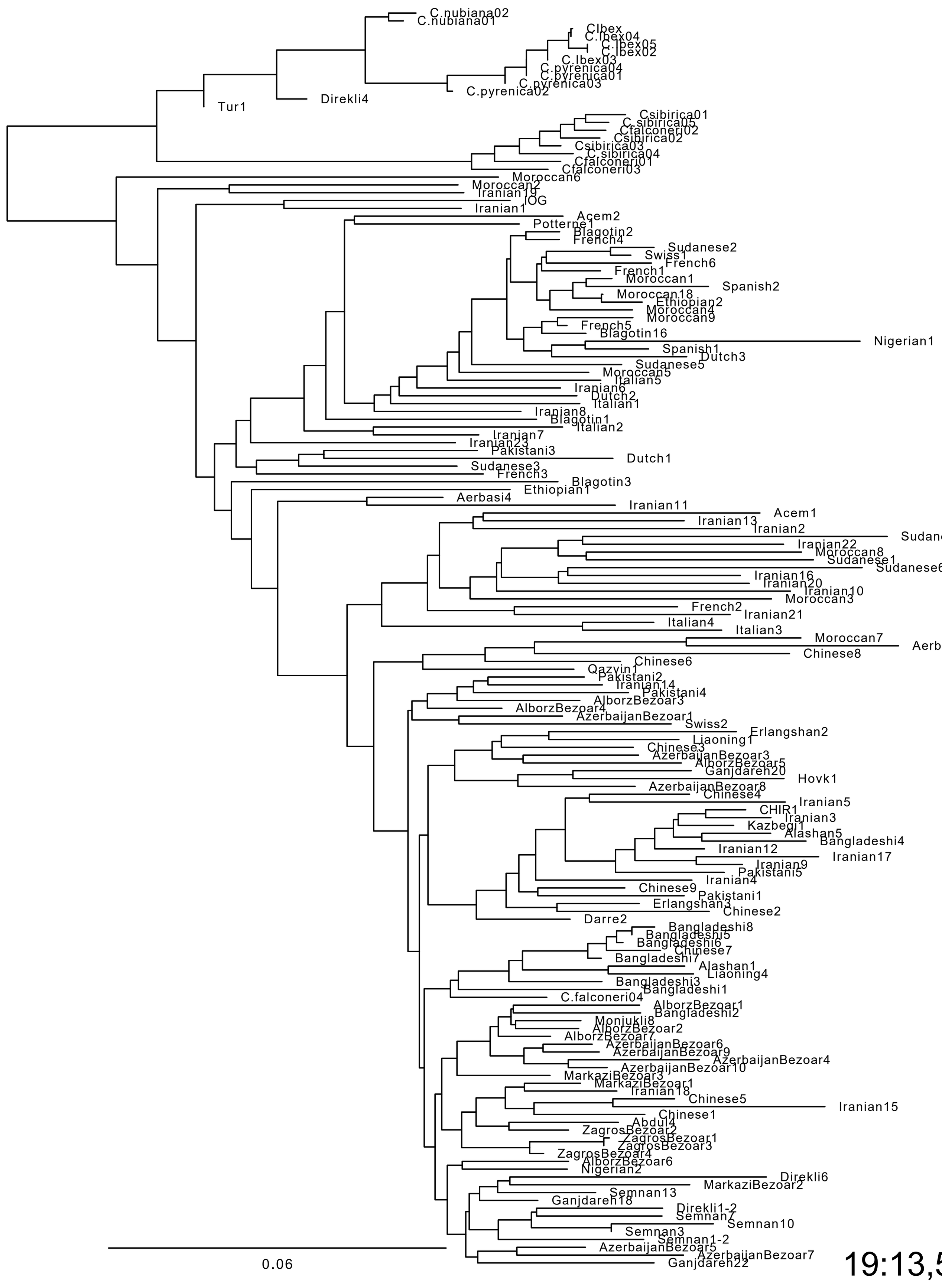

19:13,575,349-13,629,669

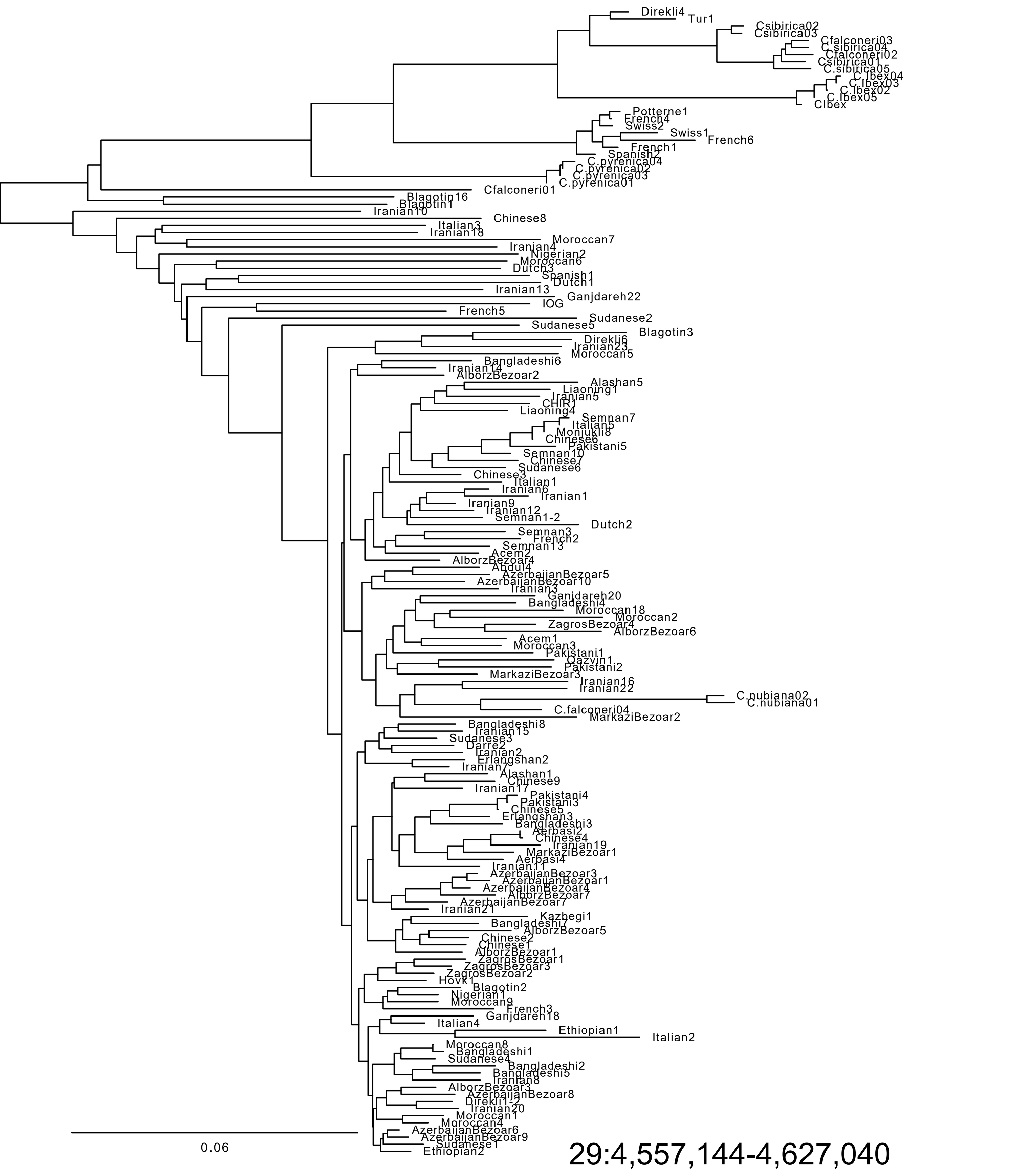

29:4,557,144-4,627,040

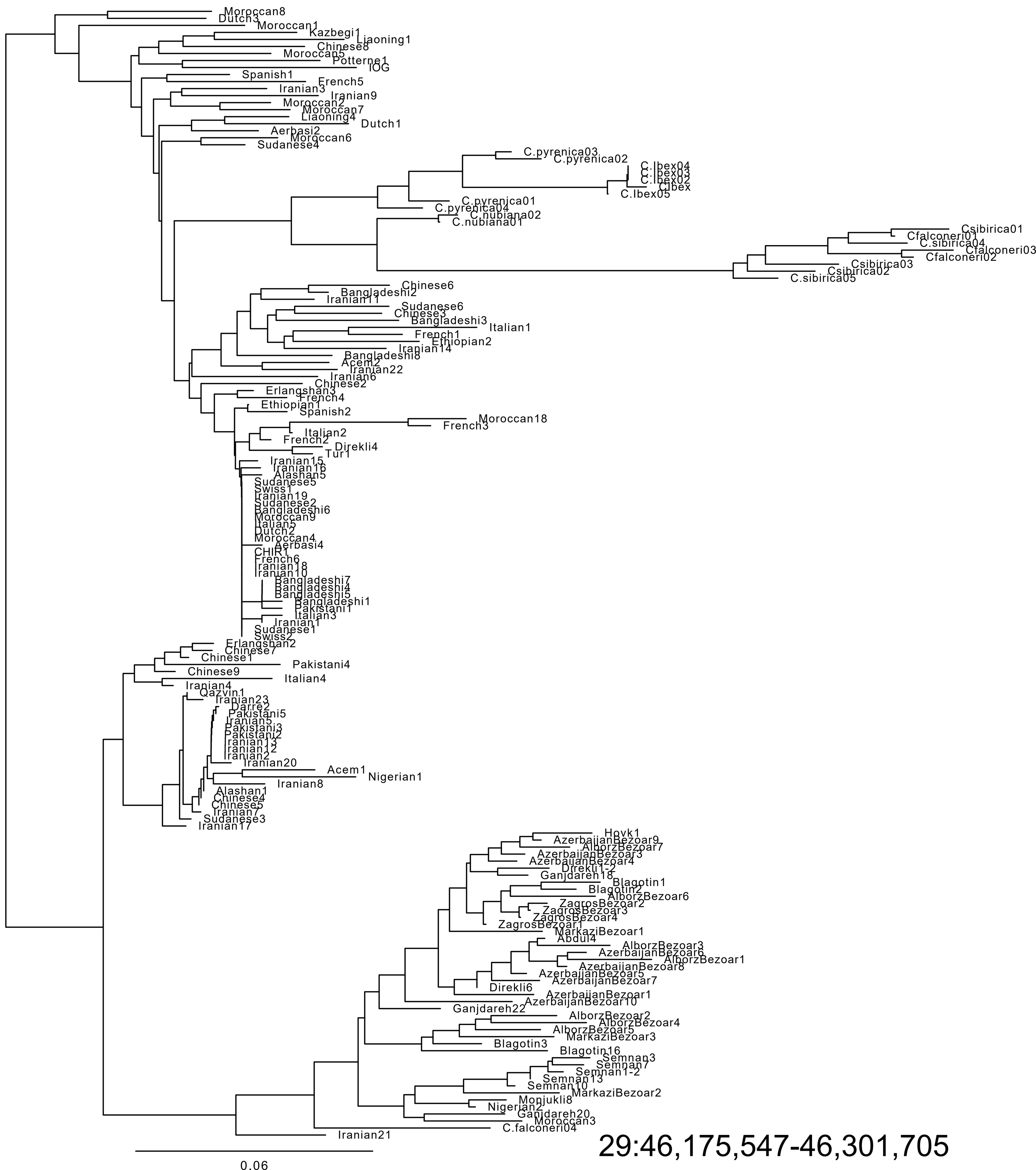

29:46,175,547-46,301,705
